# An autonomous robotic system for high-throughput phenotyping and behavioral studies of individual fruit flies

**DOI:** 10.1101/2024.08.21.607451

**Authors:** Seung Je Woo, Cheng Huang, Joan Savall, Benjamin Conrad, Junjie Luo, Mark J. Schnitzer

## Abstract

The fruit fly, *Drosophila melanogaster*, is a widely used model species in biomedical research. Despite its importance, conducting manual experiments with individual fruit flies can be challenging and time-consuming, especially for studies of individual fly behaviors. Such studies often involve cumbersome preparatory steps, such as manually tethering a fly and then positioning it within an experimental setup^1,2^. These procedures commonly require the fly to be anesthetized, and, before behavioral assessments begin, the fly must recover from anesthesia. Hence, the introduction of automated phenotyping and behavioral assays would expedite important aspects of fly research, by minimizing manual handling of flies and decreasing the net time needed for experiments. Here, we introduce FlyMAX (Fly Manipulation and Autonomous eXperimentation), an autonomous robotic system for manipulating adult flies without use of anesthesia. FlyMAX collects individual flies from a standard vial, analyzes them with computer vision, and achieves a throughput of >1,000 flies per day for high-throughput inspection and characterization assays. Robotic handling had no detectable adverse effects on fly longevity or our assessments of fly health. Moreover, the behavioral performance of flies, especially of males, was better and less variable than of flies handled manually. Our system employs deep learning-based machine vision for real-time assessments of picking quality and fly phenotypes. This enables fully pipelined, autonomous experimentation for behavioral assays with individual flies in controlled environments, which was previously infeasible. Overall, FlyMAX constitutes a promising technology to enhance the efficiency and reproducibility of research with flies and other insects in fields such as genetics, neuroscience, and drug screening.

## Introduction

*Drosophila melanogaster*, commonly known as the fruit fly, serves as a cornerstone in multiple biological disciplines^3–5^ and enables large-scale experiments, owing to its abundance, short life cycle, and cost-effectiveness^6^. The fruit fly is frequently used as a model organism for behavioral screening, drug discovery and other research assays due to the powerful genetic tools that exist^7–11^. In recent years, the pursuit of total lab automation has gathered substantial attention, for the sake of advancing the practice of fly research with high-throughput, automated experimental approaches that do not require human intervention^12,13^.

Although integrating a set of automated steps for fly experimentation is an important goal, prior efforts toward this did not yet achieve full automation. As a result, each step of a fruit fly experiment still typically relies on manual intervention. Although researchers can perform assessments of groups of flies in a vial or a custom chamber holding substantial numbers of flies (*e.g.*, hundreds), experiments focused on the behaviors of individual flies are still usually performed manually, with far fewer flies per experiment (*e.g.*, tens)^14–16^. Recent trends in fly research, particularly in neuroscience, underscore the increasing emphasis on experiments with individual flies, such as for relating complex behaviors to specific genes^17^ or circuits within the fly neural connectome^18–22^.

To conduct experiments on individual fruit flies, researchers usually resort to anesthesia, as from exposure to cold and carbon dioxide, to facilitate the handling of individual flies during setup of an experimental session. However, anesthesia may potentially harm fly health and behavior^23,24^. Moreover, the common practice of tethering flies often involves use of ultraviolet (UV) light-cured adhesives, which can also have adverse effects on fly health^25,26^.

Recently, researchers have developed some tools allowing high-throughput or automated experimentation on individual flies^27–32^. Examples of automated tasks include transferring a single fly from one place to another, monitoring an unrestrained fly^28,31^, and running experiments^27,29,30,32^. While these tools have made notable strides, they are limited to specific tasks and lack integration for performing pipelined experiments in a fully automatic way. To date, a comprehensive laboratory automation solution that prepares and performs experiments on individual flies without use of anesthesia has remained elusive.

In response to these challenges, we introduce Fly Manipulation and Autonomous eXperimentation (FlyMAX), a total laboratory automation system designed to collect adult flies without anesthesia and to conduct phenotyping and behavioral studies. With FlyMAX, we introduce an autonomous, pipelined operational framework to conduct behavioral experiments in a controlled environment, without manual intervention. Our use of deep learning-based machine vision underpins the autonomous system by facilitating real-time inference on high-resolution images. The system’s sophisticated features, including advanced phenotyping and behavioral assays with visual stimulation and thermogenetic control, underlie its capacity to precisely and efficiently handle and analyze *Drosophila* with high throughput.

We validated FlyMAX’s reliability for conducting behavioral assays by comparing its results with those from published literature. Robot-conducted fly behavior assays, including with thermogenic control, yielded results that fit with past results, affirming the system’s precision and ability to replicate known biological phenomena^33,34^. FlyMAX’s high-throughput capabilities accelerate the execution of behavioral assays, to about 50 flies in 15 hours, as compared to the approximately 60 hours needed for comparable manual experiments. These results verify the efficacy of our system and reinforce its potential as a key tool for studying the complex behavioral patterns of flies and other insect species.

## Results

### Total laboratory automation for fruit fly experiments

We present FlyMAX, an autonomous robotic system for streamlining fly handling, including fly collection, inspection, and experimentation (**Fig. 1a**). Achieving accurate and precise fly handling is crucial for the complete automation of such processes. In our earlier prototype^27^, we demonstrated a delta robot’s ability to pick flies with vacuum suction. Building on this earlier work, here we substantially enhanced the delta robot’s fly-picking capabilities. Improvements include a fly thorax detection system for more accurate and rapid target identification, increased encoder resolution to facilitate precise translations of 0.01 mm, durable robust translation mechanisms that function continuously for extended durations without failures, and redistribution of the robot platform’s mass to maintain high-speed translations without instability. Through precise and robust hardware, coupled with advanced machine vision, we engineered picking robotics components to efficiently collect a well-picked fly in about 1 min (**Fig. 1a**; **Fig. S1-S3**). Further, our new delta robot can navigate between different stations where it performs different tasks, thereby achieving full automation.

**Fig. 1.**
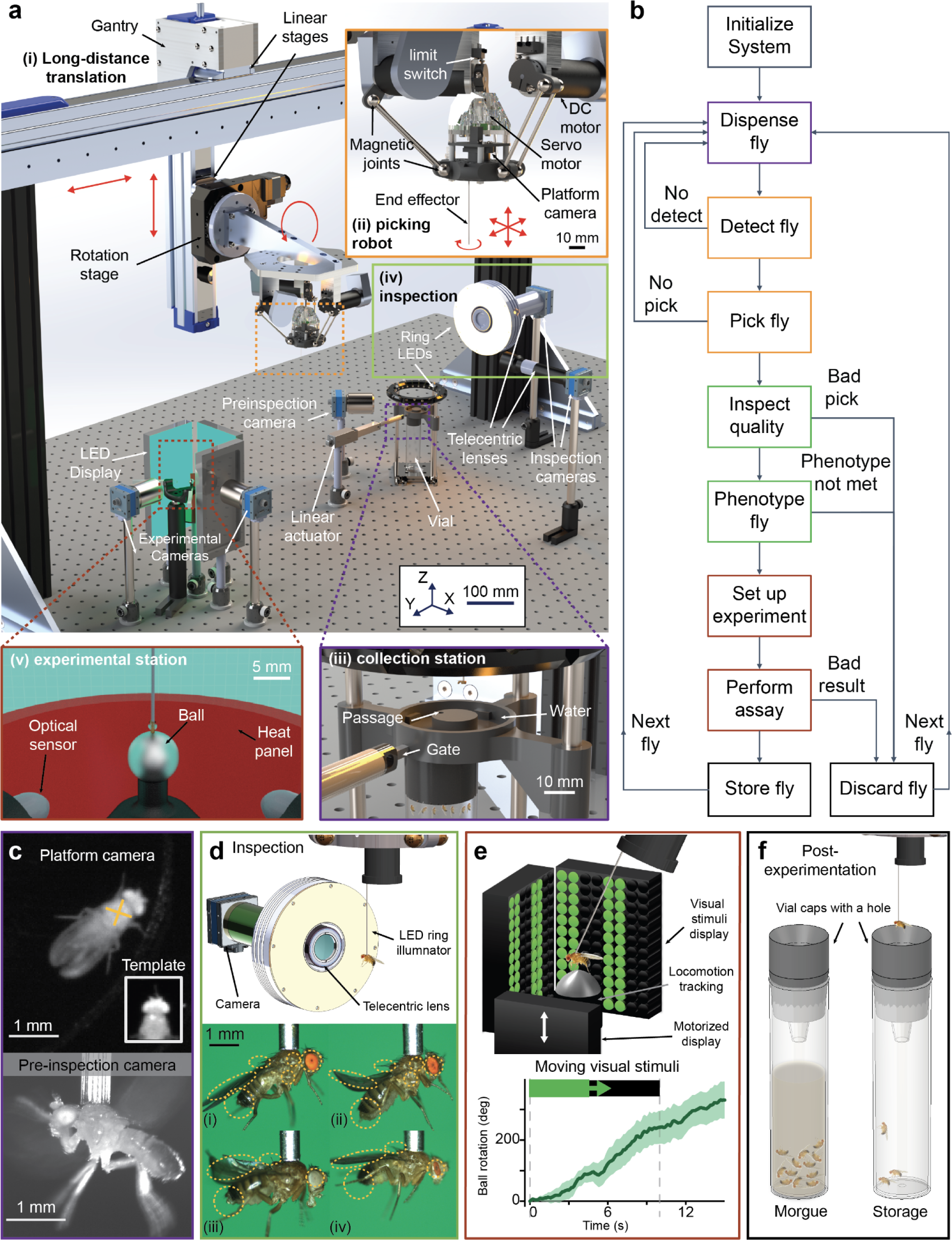
Overview of the Fly Manipulation and Autonomous eXperimentation (FlyMAX) system for autonomous preparation, phenotyping and experimentation with awake behaving flies. **a)** Computer-assisted design (CAD) schematic of the FlyMAX system. All movements, data acquisition, and computational data analytics are performed in a fully autonomous manner. Red arrows indicate translational and rotational degrees of mechanical freedom: fine translation along the X axis; coarse and fine translations along the Y and Z axes; and yaw and pitch rotations. Colored boxed areas and insets highlight the following stations and apparatus: *i*, Gantry system, which translates the delta robot between multiple different stations. The gantry has two linear translation stages (800 mm and 200 mm ranges for lateral (*Y*) and vertical translations (*Z*), respectively, and a rotational stage for controlling the fly’s pitch (±45 deg.). *ii*, Delta robot, designed to rapidly pick and transport a fly by applying suction to the thorax. The robot has a moveable platform onto which is mounted a camera, to locate the fly’s position, and an end effector, to apply suction. 3 DC rotary motors drive the movements of the robot’s 3 arms, each of which constitutes a pair of aluminum femurs capped with magnetic ball joints. A rotary servo motor actuates yaw rotations of the end effector and hence of the fly. *iii*, Collection station for picking an unanesthetized fly. A vial of flies resides underneath the picking stage, which is inside a dark chamber and surrounded by water, limiting flies’ abilities to escape by flying or walking away. A gated passageway from the vial allows flies to climb up to the picking stage but also can be closed to prevent flies from climbing up when collection is unneeded. A pre-inspection camera (shown in main figure) is used to quickly identify flies that have been incorrectly picked, so they can be released and so the robot can attempt another pick. *iv*, Inspection station for machine vision analyses of the picked fly. The station has ring LEDs for illumination and a pair of inspection cameras with telecentric lenses. *v*, Experimental station for collected flies. An LED display and a floating trackball (tracked by a pair of optical computer mice) enable tests of fly visuo-motor behavior. A heat panel beneath the trackball enables thermogenetic manipulations. **b)** Diagram of system workflow. Upon initialization, the control software acknowledges all devices and initializes them to standby mode. Next, the system starts automated task performance. Steps shown in yellow, green, and purple boxes are performed at the fly collection (**c**), inspection (**d**), and experimentation (**e**) stations, respectively. After experimentation with each fly, it is transferred to either a storage chamber or a morgue (**f**). At the initialization, FlyMAX initializes all the components for automation and checks the status of its sensors, cameras, and motors. The system begins with dispensing flies for collection (opening the gate of the collection station - passive), once it detects a fly, it will locate the target thorax and attempt to pick it with the end effector. If the robot does not successfully pick a fly, it retries the collection process by returning to the dispensing step. Once it captures a fly, at the inspection step, FlyMAX checks the quality of the picking. If it detects a poorly picked fly, it discards the fly that does not meet the desired roll and pitch angles and restarts the collection process. FlyMAX can also assess for desirable phenotypes, and if the phenotype criteria are not met, it will discard the fly and look for the next fly. After that experimental station is initialized, and the gantry and the delta robot translate to place fly on an air-suspended ball. Once everything is set up, FlyMAX performs a behavior assay. Depending on user preferences and/or the result of experiments, the fly can be discarded or stored. The delta robot then goes back to the collection station to collect another fly, and these processes iterate for autonomous experimentation. **c)** *Top*, Example image taken by the robot’s platform camera, illustrating that the angled, circularly symmetric illumination provided at the fly collection station yields high contrast images of flies. A template-matching algorithm performs fast, rotation- and scale-invariant detection of the thorax (marked with a yellow plus-sign), using the template shown in the inset. *Bottom*, Example image taken with the pre-inspection camera, showing a side view of a correctly picked fly. **d)** *Top*, CAD schematic showing the mechanical configuration after FlyMAX has transported a collected fly to the inspection station. The delta robot and gantry manipulate the fly’s yaw and pitch angles, respectively, to enable imaging of the fly from various viewpoints. *Bottom*, Sample images taken at the inspection station of flies with different phenotypes. The robot captures high-resolution images that distinguish features from different phenotypes. Yellow ellipses enclose distinctive phenotypic features, including wings, scutellar bristles, eyes, abdomen, and humeral. Flies shown in panels i–iv are wildtype, *Df(2L)-cl-h1*/*CyO*, *amos^Roi-1^*, *w^1118^*/*Dp(1;Y)y^+^*;*CyO*/*sna^Sco^*;*MKRS*/*TM6B,Tb^1^*, and *Bar^1^*. **e)** *Top*, CAD schematic showing a configuration in which FlyMAX has transported a collected fly to the experimental station. The robot has 5 degrees of freedom to accurately position the fly on an air-suspended ball that is surrounded by an LED display providing visual stimulation. One of the LED display panels can be slid vertically up and down, to provide one of the cameras a side view of the behaving fly before the start of optomotor experimentation, enabling verification that the fly is in the correct position. *Bottom*, Example data showing a fly’s locomotor responses on the ball to the presentation of a moving visual stimulus. The locomotion tracking setup measures a ball rotation caused by a fly on a ball, and the data represents the rotation with an ON visual stimulus. Dashed dark gray lines mark the start and end of the visual stimulus. The visual stimulus moved from left to right, and positive displacement on the graph indicates that the fly rotated in the same direction. **f)** Stations for holding flies after the end of experimentation. Gentle air pressure at the tip of the end effector of the delta robot transfers a collected fly to a vial. *Right*, A storage vial to keep flies for future experiments. *Left*, A morgue for discarded flies.

The autonomous workflow unfolds as follows (**Fig. 1b**): after initializations, including homing the motors of the delta robot, FlyMAX opens a gate of the collection station to dispense flies. The robot detects a fly’s thorax and picks the fly at the estimated position. These steps repeat until the robot successfully collects a fly. After collection, multiple stages of machine vision inspect the fly to ensure the picking is proper for subsequent experimental steps. Another inspection step ascertains the fly’s phenotypes. If the fly fails to meet user-defined criteria, such as having curly wings or white eyes, for qualification, FlyMAX discards it and begins the picking process again. Once a fly passes all inspections, the system transports it to an experimental setup for user-defined experiments. At the end of experiments, FlyMAX either stores the fly in a designated vial or discards it. This workflow iteratively runs until a specified number of user-defined experimental procedures are done.

The process begins at the collection station, where FlyMAX transports the delta robot module to pick flies designated for the experiments. The station comprises a stage arena, a fly vial, an infrared illuminator, a gate, and other subcomponents (**Fig. 1c**). Enclosed in a black box, the collection station provides a dark environment for flies. Leveraging the flies’ geotactic behavior^35^, they climb up to the arena when the gate is open. Water encircles the arena, preventing flies from escaping and keeping them less active. We fine-tuned the angle of the infrared lighting on the stage to achieve optimal contrast for the delta robot’s camera (**Fig. 1c**, *Top*; **Fig. S3**). Using a template matching algorithm, which is fast, rotation- and scale-invariant, the robot swiftly identifies the picking target—usually the thorax of a fly—and promptly lifts and holds the targeted fly using a gentle vacuum (**Fig. S1**). The system then manipulates the fly in 3 translational and 2 rotational degrees of freedom (DOF), facilitating precise and accurate transportation to inspection and experimental stations.

At the inspection station, each picked fly undergoes a machine vision analysis using convolutional neural networks that determine the quality of the pick and the fly’s phenotypes (**Fig. 1d**). Although FlyMAX accurately estimates the position of a fly’s thorax and can make a pick within ∼100 ms, this can still be challenging if the fly moves abruptly. To account for this, the quality assessment routine verifies a correct pick on the thorax, gauged by estimating the pitch and roll of the tethered flies. The success criteria for a suitable pick vary based on the type of the experiment. Generally, the pitch should be between –15 deg. and 45 deg., and the roll should be within –10 deg. to 10 deg. In addition to this quality check, the inspection algorithm uses deep neural networks to perform phenotypic assessments of fly morphology.

Next, FlyMAX transports the inspected fly to a station designated for experimentation. Our experimental station tested the fly’s visual motion driven behavior, but a broad range of other experimental tests would be feasible (**Fig. 1e**). Our experimental testing rig incorporates a display for stimulus presentation, an air-suspended ball tracking setup for behavior recording, and a temperature controller that allows thermogenetic manipulations. Users can also customize this station for other types of assays and tests and can synchronize the hardware for these rigs with the FlyMAX software, for instance to ensure precise timing for all data collection.

We used morgues and storage vessels for the disposal and preservation of captured flies, respectively. The air pressure from the end effector of the delta robot releases flies from the vacuum tether into vials with narrowly opened caps (**Fig. 1f**). Typically, flies with poor pick quality and those filtered out due to undesirable phenotypes were discarded, while those flies from which we took experimental data were preserved. Altogether, the robot effectively conducts automated experiments using methods akin to those performed by biologists, but with shorter durations and without need for CO_2_, chilling, or ultraviolet (UV) light.

### Robotic manipulation preserves fly health and facilitates experimentation

To assess the impact of robotic handling on fly health, we performed a comparative analysis of survival rates between flies handled by FlyMAX and those in a control group (**Fig. 2a,b**). FlyMAX sorted the robot-handled group within 0-1 day from eclosion; we manually sorted the control group of flies under CO_2_ anesthesia. The longevity curves for flies subjected to robotic handling were statistically indistinguishable from those of the control group, for females (*P* = 0.33; *n* = 94 and *n* = 132 for robotically and manually handled groups, respectively; log-rank test) and males (*P* = 0.97; *n* = 122 and *n* = 132 for robotically and manually handled groups, respectively; log-rank test).

**Fig. 2.**
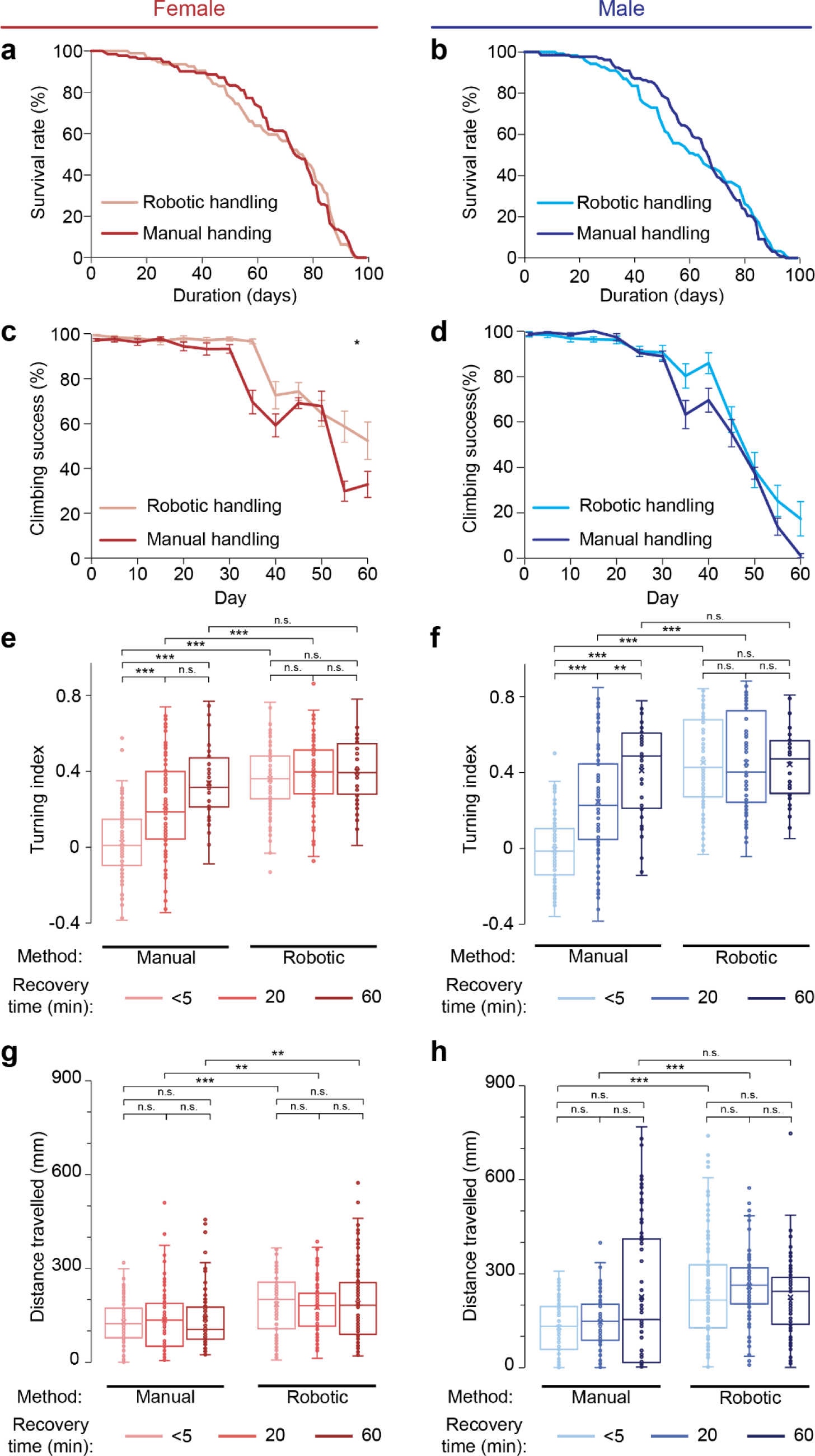
Robotic handling does not affect fly longevity or assessments of health. **a, b)** Lifespan curves of female (**a**) and male (**b**) Canton-S flies. We used FlyMAX to collect and store robotically handled (**Methods**) flies on Day 0 and then performed a lifespan assay [*n* = 94 and 132 female flies, respectively, in the robotically and manually handled groups (*P* = 0.33; log-rank test); *n* = 122 and 132 male flies, respectively, in the robotically and manually handled groups (*P* = 0.97)]. **c, d)** Results of a climbing assay performed with female (**c**) and male (**d**) Canton-S flies on different days after either robotic or manual fly handling. We set 175 mm for a target line, a climbing distance for measuring how many flies climbed further. We utilized FlyMAX to collect and store robot-handled flies on Day 0 and then performed the first climbing assay on the next day. We performed the climbing assays of the remaining days with the interval of 5 days (**Methods**). [*n* = 8 vials with a total of 125 female flies in the robotically handled group and *n* = 9 vials with a total of 154 female flies in the manually handled group (*P* = 0.0080; Kruskal-Wallis ANOVA; \**P* < 0.05). Error bars: s.e.m. across 8 vials and 9 vials for robotically handled and manually handled group, respectively; *n* = 8 vials with a total of 106 male flies in the robotically handled group and *n* = 9 vials with a total of 114 male flies in the manually handled group (*P* = 0.95; Kruskal-Wallis ANOVA). Error bars: s.e.m. across 8 vials and 9 vials for robotically and manually handled groups, respectively]. **e, f)** Visual Sensory assay of female (**e**) and male (**f**) Canton-S flies. We applied chilling and UV light exposure to one group of flies for manual tethering, while we handled the other group using the robot without exposing them to these conditions. For the group with manual tethering, we waited less than five minutes, twenty minutes or an hour to recover before performing experiments. The box-and-whisker plot shows the turning indices of the flies handled manually or robotically with waiting 3 different periods prior to experiments. Boxes span the 25th-75th percentiles, horizontal lines denote median values, whiskers span 1.5 times the interquartile distance, and open circles are individual data points. This description applies to all the box-and-whisker plots in this paper. For comparison, we also waited the same amount of time for the robotically handled flies. For each **(e)** and **(f)**, flies with *n* = 100, 100 and 40 for manual methods with <5 minutes, 20 minutes, and 1 hour recovery time, respectively. Flies with *n* = 100, 100 and 40 for robotically handled methods with 15 minutes and 1 hour recovery time, respectively. (\*\**P* < 0.01, \*\*\**P* < 0.001; n.s.: not significant; Kruskal-Wallis ANOVA followed by post-hoc rank sum tests with Holm-Bonferroni correction). **g, h)** Distance traveled of female (**g**) and male (**h**) flies. Of flies (**e**) and (**f**), each data point indicates the average distance each fly traveled (**Methods**) (\*\**P* < 0.01, \*\*\**P* < 0.001; n.s.: not significant; Kruskal-Wallis ANOVA followed by post-hoc rank sum tests with Holm-Bonferroni correction).

To assess the potential impact of robotic manipulations on fly climbing ability over time, we conducted a longitudinal climbing assay to evaluate the geotactic behaviors of robotically handled flies (**Fig. 2c,d**; **Fig. S6**). On day 0–1 from eclosion, we used FlyMAX to sort the sex of the robotically handled flies without anesthesia; we manually sorted the control group of flies using CO_2_ anesthesia. We initiated climbing assays on the following day. We conducted subsequent climbing assays without anesthesia every 5 days and observed a statistically significant superiority in the climbing performance of robotically handled female flies as compared to those handled manually (*P* = 0.0080; *n* = 8 vials with a total of 125 flies in the robotically handled group and *n* = 9 vials with a total of 154 flies in the manually handled group; Kruskal-Wallis ANOVA). Male flies showed no such differences across the two groups (*P* = 0.95; *n* = 8 vials with a total of 106 flies in the robotically handled group and *n* = 9 vials with a total of 114 flies in the manually handled group; Kruskal-Wallis ANOVA). We conducted these climbing assays on the flies that survived each day of the experiment; we therefore recognize the possibility that a survivor bias might, in principle, have influenced the results. Nonetheless, the findings suggest that robotic handling did not adversely affect climbing, and, in female flies, may have even led to better performance than manual handling.

Next, in experiments with tethered flies, we observed that flies tethered by FlyMAX were more active in their behavior than those tethered manually. To quantify these differences, we conducted locomotor behavior experiments involving ON-OFF moving gratings, in which a square-wave pattern that alternates between light (ON) and dark (OFF) edges moves across the visual field (**Methods**). Within each robot-handled and control group, we divided the flies into three subgroups based on their recovery time before the start of experiments: <5 min, <20 min or <60 min. Notably, flies in the manually handled control group exhibited the poorest turning index^34^—a unitless metric that represents the ratio of yaw velocity to pitch and roll velocities **(Methods**)—when we conducted experiments immediately after tethering, as compared to flies that recovered for 60 min (**Fig. 2e,f**; \*\**P* < 0.01, \*\*\**P* < 0.001; Kruskal-Wallis ANOVA followed by post-hoc rank sum tests with Holm-Bonferroni correction). By comparison, the performance of robot-tethered flies remained consistent across all three subgroups, with no significant differences observed among them. This suggests that FlyMAX can perform experiments with picked flies immediately, without need for a recovery interval after picking.

Because the turning index does not account for the extent of fly movement, we also analyzed the distances that flies traveled during the experiments (**Fig. 2g,h**; \*\**P* < 0.01, \*\*\**P* < 0.001; Kruskal-Wallis ANOVA followed by post-hoc rank sum tests with Holm-Bonferroni correction). The robot-handled flies displayed significantly higher activity levels compared to conventionally tethered ones, potentially yielding more accurate and reliable results. Overall, our experiments on fly survival, climbing and activity levels after robotic picking demonstrate that FlyMAX does not harm, and may even benefit, the health of picked flies, as compared to manual handling.

### High-throughput processing of over a thousand flies per day

To evaluate the throughput capacity of our system in preparing suitable flies for experiments, we devised a workflow with picking of flies at the collection station and assessment of their quality at the inspection station (**Fig. 3a**; **Fig. S7**). We considered picked flies to be ‘qualified’ for this assessment if their roll was within the ranges of -10 deg to 10 deg and their pitch was within -15 deg to 45 deg. Within this pitch range, we categorized -15 deg to 5 deg as ‘low pitch’, 5–25 deg as ‘medium pitch’, and 25–45 deg as ‘high pitch’ (**Fig. 3b**). By autonomously collecting and inspecting individual flies, FlyMAX identified >1,000 qualified flies for experiments within 17 h (**Fig. 3c**). The average time to collect a qualified fly was about a minute, much briefer than the recovery time for manually prepared flies^1,22,36^ (**Fig. 3d**; **Fig. S8**).

**Fig. 3.**
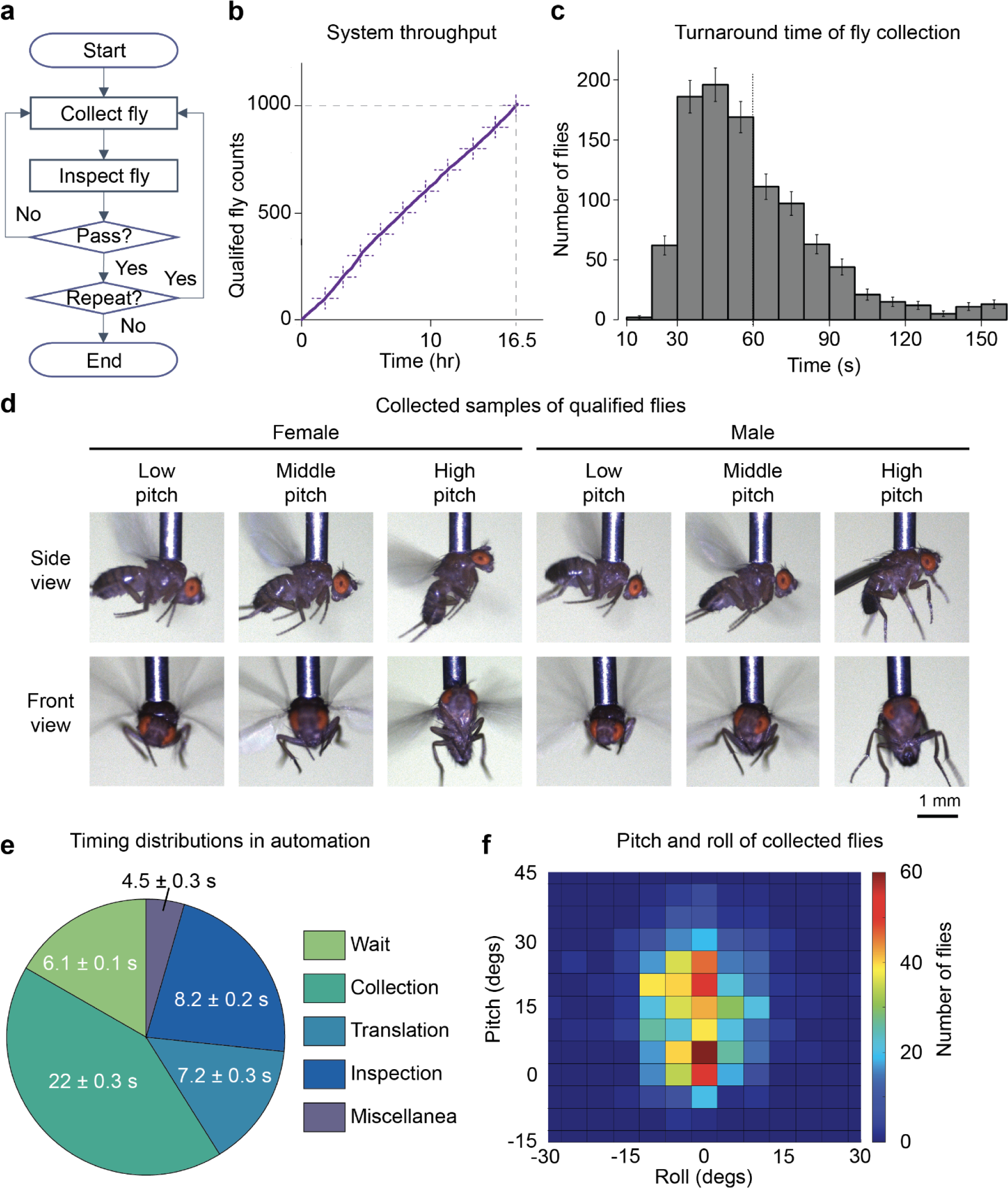
The automated collection system can select >1,000 flies per day, suitable for incorporation into high-throughput assays and screens. **a)** Workflow of collecting qualified flies for experiments. FlyMAX collects a fly and inspects it to check it captures the thorax with good roll and pitch dimensions (**Fig. S7**, **Methods**). The system repeatedly collects flies until the collected amount reaches users’ desired number of flies. **b)** Numbers of flies collected as a function of time elapsed during robot operation in collecting qualified flies. A dotted cross displays every 100^th^ collected fly. **c)** In fly collection, turnaround time measures the duration needed to repeatedly detect, pick, inspect, and discard flies until it picks a qualified fly. This histogram displays the distribution of the turnaround time. The red dashed line indicates the average of the turnaround time. **d)** Samples of collected flies. The first and third rows show side-facing views of flies, and the second and fourth rows show front-facing views. We classified pitches ranging from -15 to 45 degrees into three categories: low (-15 to 5 degrees), middle (5 to 25 degrees), and high (25 to 45 degrees). **e)** Timing distribution of qualified fly collection. ‘Wait’ is the time taken to detect a fly on the collection stage. ‘Collection’ is the time taken to track a detected fly and pick it. ‘Translation’ is the time taken to move along stations to transfer a collected fly from the collection station to the inspection station. ‘Inspection’ is the time taken to check the quality of a picked fly. ‘Miscellanea’ is the remaining time taken in running the automation including homing, error-handling, cleaning, etc. Error bars: s.e.m. calculated from the time data of 1,007 qualified flies. **f)** Pitches and rolls of collected and qualified flies. *n* = 1,007. Each bin ranges 5 deg. with [-2.5, 2.5] deg. around the centers. The color indicates the number of the flies whose rolls and pitches are within the bin.

Uninterrupted and continuous operation for extended durations without manual intervention is crucial for achieving high-throughput experimentation. Prior to quantifying our final results, we conducted multiple runs of automated tasks, identifying errors or cases of prolonged delays, such as flies not climbing up for a long duration, device communication glitches, and clogs inside the end effector. We addressed these issues by implementing vial stimulation, error-handling algorithms, and automatic end effector cleaning tasks.

Aiming for higher throughput, we installed a pre-inspection module at the collection stage to discard poorly picked flies at an early stage (**Fig. S9**). Using separate infrared light illumination, FlyMAX verifies whether the delta robot accurately picked the thorax of a fly. A dedicated neural network model for infrared thorax verification rejects non-thorax-picked images only if it is highly confident, *i.e*., when the probability of a bad pick exceeds 90%. This module led to early filtration of 70% of inadequately picked flies, substantially reducing the processing time compared to filtering these flies out at a later stage. Out of 1,615 collected flies, FlyMAX properly retained 1,007 flies that met the criteria. The distribution of time per fly (**Fig. 3e**) indicates that a majority of the time elapsed during the collection phase (46 ± 0.7 %, 22 ± 0.3 s), representing the duration taken by the robot to pick a fly’s thorax. This duration encompassed the time the robot tracked a target fly until it became stationary. The wait phase in the distribution constitutes the time taken for the robot to detect the first fly in each trial (13 ± 0.3 %, 6.1 ± 0.1 s). The inspection phase combines the time for pre-inspection at the collection station and for the inspection station (17 ± 0.3 %, 8.2 ± 0.2 s). The translation phase accounts for the time to transport the delta robot between the collection and inspection stations (15 ± 0.3 %, 7.2 ± 0.3 s). The miscellaneous phase includes the time for actions other than the above phases, such as homing motors, error handling, and cleaning. The pitch and roll distributions of collected flies show two local maxima, with pitch values displaying greater variability than roll (**Fig. 3f**). We allowed a wider range of pitch, from -15 to 45 deg, because FlyMAX has an additional rotation stage to correct for this variation by precisely aligning flies in the pitch dimension.

### Real-time phenotyping with deep learning-based machine vision

We employed deep neural networks for image classification to sort files based on their phenotypic features. At the inspection station, flies undergo initial quality inspection, and upon passing this, they proceed to phenotypic classification (**Fig. 4a**; **Fig. S10**). For quality assessments, one option was to use standard convolutional neural network (CNN) models^37^, resizing the original image to 224 × 224, a commonly used set of image height and width dimensions for these models (**Fig. 4b**). However, this resizing can compromise image resolution, potentially leading to a loss of crucial details needed to determine phenotype, particularly for phenotypes expressed in specific body parts like eyes and bristles on scutellum. The images we captured at the inspection station have a resolution of 2,448 × 2,048 pixels. Using models compatible with this image size presents challenges, as does downsampling the images to match standard input sizes. Using the original size may cause memory and computational issues, while downsampling risks losing subtle differences in phenotypes. To address this, we devised a two-stage model. The first stage detects positions related to phenotypes, and the second stage classifies the cropped image centered on the detected positions at a full resolution (**Fig. 4c**). The first stage employs a pose estimation model, where the CNN model’s output indicates predicted positions. We utilized the DeepLabCut framework^38^, deep learning software for animal pose estimation, to label and train the model for identifying phenotypic locations. For the second stage, we used EfficientNet^39^, which enabled us to perform phenotypic classification on larger image sizes without losing phenotype details. This two-stage approach combines the efficiency of pose estimation with the precision of high-resolution classification.

**Fig. 4.**
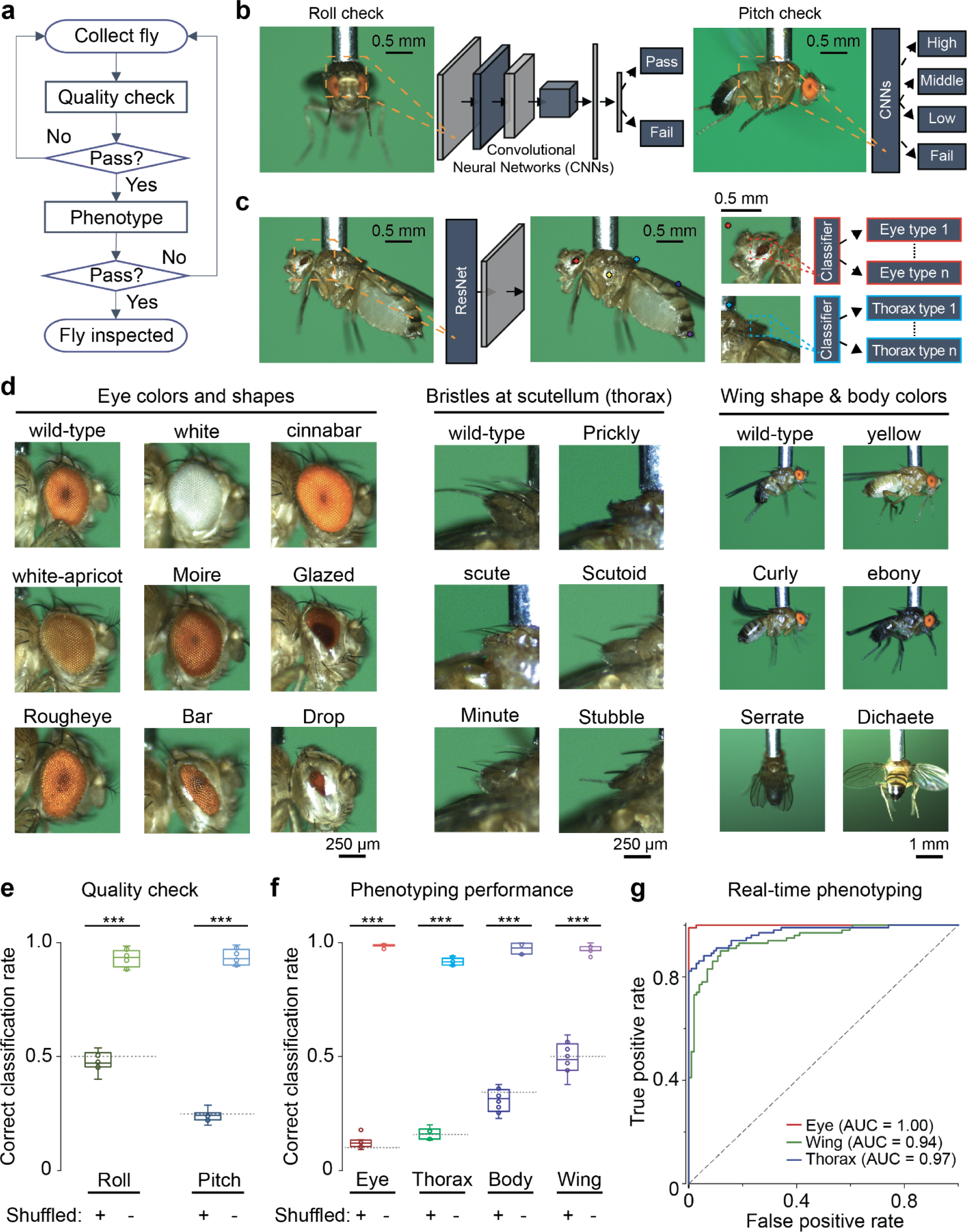
Deep-learning based machine vision, combined with robotic manipulation, allows the inspection of quality and phenotypes of collected flies. **a)** Workflow of real-time inspection for experimentation. The inspection involves two steps: picking-quality checking and phenotyping. At the quality checking process, the system determines if a picked fly has good roll and pitch values. At the phenotype process, it classifies phenotypes of the fly. **b)** Quality checking with convolutional networks (CNNs). The collected front view and side view go through the CNNs trained with classifying roll or pitch quality of flies, respectively (**Methods**). We trained a dedicated classifier for each roll or pitch classification. **c)** Phenotyping with CNNs. To accurately identify features for phenotypic classification without sacrificing resolutions, the system first identifies key features with the CNNs whose output goes through transposed convolutional networks to spatially locate the positions. Each identified keypoint goes through CNN classifiers for eye, thorax, wing, and body color (**Methods**). **d)** Sample images of phenotypic features captured by FlyMAX. Various genotypes exhibit distinctive eye colors and shapes, bristles, body colors, and/or wing shapes. Phenotypes denoted by capitalized letters indicate dominant phenotype. For eye colors and shapes, Bar has a rectangular eye shape, cinnabar has a bright red eye color, Drop has a small eye shape, Moire has brownish eye color, Glazed has no photoreceptors in the eye and darker red eye color, Rougheye has rough and irregular eye facets, white has white eye color, and white-apricot has orange eye color. For bristles at the scutellum, Prickly has very short and twisted bristles, scute has no bristle on the posterior scutellum, Scutoid has either no or very few on the scutellum, Minute has shorter bristles, and Stubble has short thick bristles. For wings, Curly has a curled wing, Serrate has notches at wing tips, and Dichaete has 45 deg. extended wings. For body colors, yellow has a brighter body color, and ebony has a darker body color compared to the wild type. **e)** Performance in quality checking. For roll checking, the labels were ‘bad’ and ‘good’; *n_train_* = 2,156 and *n_test_* = 308. For pitch checking, the labels were ‘bad’, ‘low’, ‘middle’, and ‘high’ as indicated in Fig. 3b; *n_train_* = 2,541 and *n_test_* = 363. We trained all networks and tested with 8-fold cross validation. The box-and-whisker plot shows the accuracy of picking quality classification in roll and pitch using the images captured by FlyMAX at the inspection station. Gray dotted lines indicate chance levels (\*\*\**P* < 0.001; Wilcoxon rank sum test). **f)** Performance in phenotyping. We used nine classes of eye phenotypes indicated in Fig. 3d as labels for eye classification; *n_train_* = 1,798 and *n_test_* = 526. For bristle at scutellum classification, we used six classes indicated in Fig. 3d labels; *n_train_* = 1,118, *n_test_* = 261. For body color classification, we used yellow and ebony along with wild type as labels; *n_train_* = 2,121 and n_test_ = 886. For wing classification, we used Curly and wild type as labels; *n_train_* = 1,110 and *n_test_* = 357. We trained all networks and tested with 8-fold cross validation. The box-and-whisker plot shows accuracy of phenotype classification in eye, thorax (scutellum), body color, and wing using the images captured by FlyMAX at the inspection station. Gray dotted lines indicate chance levels (\*\*\**P* < 0.001; Wilcoxon rank sum test). **g)** Real-time phenotyping experiments. FlyMAX performed phenotyping wild-type flies (BDSC# ID 64346; CS-Canton) and flies with white eyes, curled wings, and scutoid thorax (BDSC# 3703; w[1118]/Dp(1;Y)y[+]; CyO/nub[1] b[1] sna[Sco] lt[1] stw[3]; MKRS/TM6B, Tb[1]). ROC curves of real-time classifications between two groups. *n* = 100 for each group. Purple dotted line indicates 0.5 of area under the curve.

Among numerous phenotypic features, we selected eye colors, eye shapes, bristles at the scutellum, body color, and wings based on their common usage as genotype markers (**Fig. 4d**)^40,41^. For example, compared to the wild type, scute has no bristles on the posterior scutellum, Scutoid has no or very few bristles, Minute has short and thin bristles, and Stubble has short and thick bristles.

To validate the inspection performance, we trained the roll and pitch classifiers using previously collected data. For roll classification, we used 2,156 images for the training set and 308 for testing. For pitch classification, we used 2,541 images for training and 363 for the test set. In both roll and pitch classifications, the average classification rates were about 95% and were significantly different from those trained with randomly labeled data (**Fig. 4e**; \*\*\**P* < 0.001; Wilcoxon rank sum test).

To assess the performance of our phenotyping system, we trained classifiers for eye, bristle, body, and wing phenotypes using previously collected data. For eye classification, we used 1,798 images for the training set and 526 for the testing set; for bristle classification, 1,118 for the testing set and 261 for the testing set; for body classification, 2,121 for the training set and 886 for the testing set; and for the wing classification, 1,110 for the training set and 357 for the testing set. To prevent classifiers from relying on incorrect features, we diversified the fly lines to include various combinations of phenotypes (**Table 2**). In phenotype classifications, the rate of successful classification for eye phenotype was 99%, 92% for thorax, 97% for body color, and 97% for wing, all of which were significantly different from success rates of classifiers trained with randomly labeled data (**Fig 4f**; \*\*\**P* < 0.001; Wilcoxon rank sum test).

To validate the real-time phenotyping capability, we automated FlyMAX using these trained models to perform phenotype classification. We mixed a fly line with white eye, Scutoid bristles, and Curly wings (w[1118]/Dp(1;Y)y[+]; CyO/nub[1] b[1] sna[Sco] lt[1] stw[3]; MKRS/TM6B, Tb[1]) with a wild type line into a single vial. FlyMAX randomly collected a fly, inspected it, and then classified its phenotypes. A receiver operating curve (ROC) analysis indicated that eye classification had the highest fidelity (area under the curve (AUC) of 1.0), whereas wing (AUC=0.94) and bristle (AUC=0.97) classifications were around 95%. These results demonstrate the robot can effectively perform real-time classification.

### Autonomous assessments of stimulus-response behavior

To automate assessments of visual sensory-motor behavior, the robot securely positioned individual flies on a trackball that tracked their locomotor behavior as they responded to a video display (**Fig. 5a**). We equipped this station with a temperature control unit for thermogenetic manipulations that was capable of heating, using a thermistor, and cooling, using a fan (not shown in the figure). The station also allowed automated configuration of experimental settings, including fly positioning, temperature regulation, trackball and visual display setup. After initializing these components, FlyMAX started the experiments (**Fig. 5b**; **Fig. S11,12**).

**Fig. 5.**
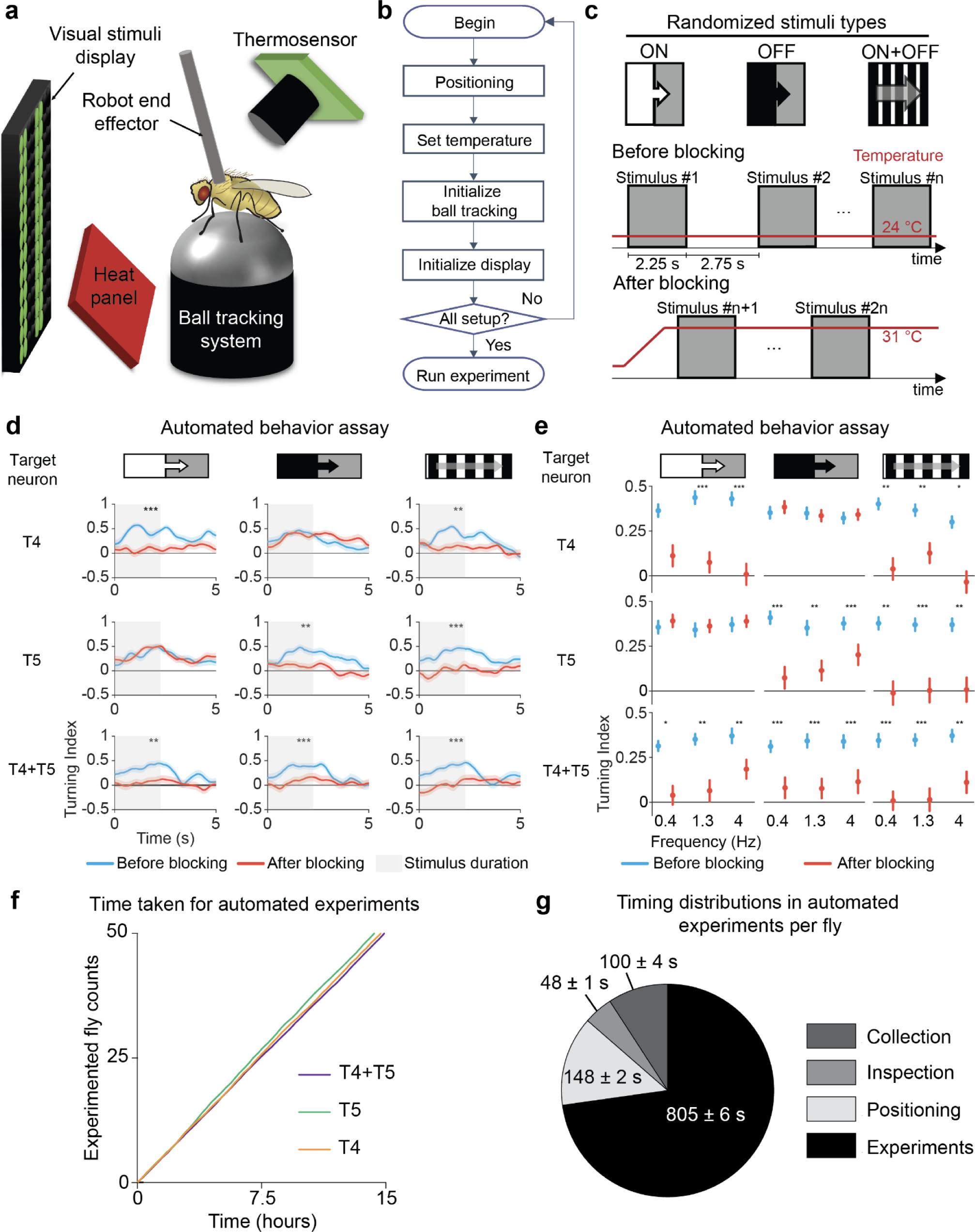
Automated assessments of stimuli-response behavioral assays on individual flies. **a)** Diagram of the experimental setup for visual sensory motor behavior experiments. FlyMAX manipulates a picked fly vacuumed by the end effector for positioning. An LED display surrounds the fly with 270 deg. A thermosensor monitors fly body temperature for themogenetics control. The temperature measurement feedbacks to the system to determine whether to turn on or off the heat panel to maintain a certain temperature. A ball tracking setup holds an air-suspended ball, and a pair of optical computer mice records the locomotion with infrared sensors. **b)** Workflow of setting up automated experiments. In the beginning, FlyMAX positions a collected fly on a ball (**Fig. S12a,b**). Then, the temperature controller sets a desired temperature (**Fig. S12c**). The pair of optical computer mice initializes to record ball tracking. After that, the visual display initializes with one of its panels translated to enclose the behavior rig. FlyMAX syncs all devices before proceeding to conduct experiments. **c)** Automated experimental sequence for visual sensory motor behavior assay. We used three types of stimuli, ON, OFF, and ON-OFF. Each stimulus pair comprises the same stimulus pattern with opposite directions of stimulus movement. The experimental sequence begins with a control group session, performed at a permissive temperature (22°C), followed by a blocking group where thermogenetics blocks the target neuron with a restrictive temperature (31°C). The system monitors temperature, so the experiments with thermogenetic perturbations begin once it reaches the restrictive temperature (**Methods**). **d)** Turning index (TI) curves along with time. FlyMAX performed experiments on control group sessions at the permissive temperature and on blocking group sessions at the restrictive temperature. The plot displays visual sensory behaviors under moving grating with 1.4 Hz as a temporal frequency. The robot applied stimulus at time = 0 s and stopped at time = 2.25 s (gray shading: stimulus duration). Lighter shaded areas include s.e.m. of the turning indices over all 50 flies at each time point. We used 50 flies per group with targeted neurons: T4 (*R54A03>shi^ts^*), T5 (*R42H07>shi^ts^*), and T4+T5 (*R42F06>shi^ts^*). Each fly underwent different stimulations with both permissive and restrictive temperature as described in Fig. 5c, and each datapoint includes standard error of mean (\*\**P* < 0.01, \*\*\**P* < 0.001; Friedman ANOVA followed by post-hoc Wilcoxon signed-rank tests with Holm-Bonferroni correction on average turning index for each fly over the first 2.25 s after each stimulus onset). **e)** Mean±s.e.m. turning indices of 50 flies with varying stimuli and temporal frequencies. Each column represents a different stimulus type, and each row represents a different target neuron. Each plot displays turning indice values across different frequencies. Blue and red plots indicate behaviors before and after blocking neurotransmissions at permissive and restrictive temperatures, respectively. Each dot represents the mean value of the turning index. We used 50 flies per group with targeted neurons: T4 (*R54A03>shi^ts^*), T5 (*R42H07>shi^ts^*), and T4+T5 (*R42F06>shi^ts^*). Each fly underwent different stimulations with both permissive and restrictive temperatures as described in Fig. 5c (\**P* < 0.05, \*\**P* < 0.01, and \*\*\**P* < 0.001; Friedman ANOVA followed by post-hoc Wilcoxon signed-rank tests with Holm-Bonferroni correction on average turning index for each fly over the first 2.25 s after each stimulus onset). **f)** Numbers of flies experimented as a function of time elapsed during FlyMAX’s operation in automatic behavior assays. For each target, we experimented on 50 flies using the automated experiments protocol described in (**c**). **g)** Timing distribution of automated behavior assay. ‘Collection’ is the time taken to collect a fly. ‘Inspection’ is the time taken to perform a quality check. ‘Positioning’ is the time taken to move along stations to transfer and position a collected fly. ‘Experiment’ is the time taken to perform a behavior assay. ‘Miscellanea’ is the remaining time taken in running the automation including homing, error-handling, waiting, etc. Error bars: s.e.m. across 150 flies, with 50 flies per genotype (T4, T5, and T4+T5).

To validate our approach, we focused on T4 and T5 neurons in the fly’s visual system and used the FlyMAX system to verify their function in motion detection. Prior studies revealed that T4 and T5 neurons in the medulla optical lobe of *Drosophila* are critical components of the elementary motion detectors, responding to the direction of movement in the flies’ visual field^33,42,43^. T4 neurons are selectively sensitive to moving ON-edges (transitions from dark to light), whereas T5 neurons respond preferentially to moving OFF-edges (transitions from light to dark)^44^. When a bright light pattern moves across the fly’s visual field (an ON stimulus), the T4 neurons detect this change, prompting the fly to move in the same direction as the light pattern. On the other hand, if a dark pattern moves across its field of view (an OFF stimulus), the T5 neurons detect this motion, prompting the fly to move in the same direction of the moving darkness. When an ON-OFF stimulus, a square-wave pattern that alternates between ON and OFF visual stimuli, moves across the fly’s field of view, both T4 and T5 neurons detect this, prompting the fly to move in the same direction as the moving pattern. Hence, we designed the FlyMAX experimental station so that it can thermogenetically perturb the activity of T4 and T5 cells and track the effects of these perturbations on fly behavior. We expected that, during thermogenetic perturbations of T4 or T5 neural activity, flies would not properly follow moving light or dark stimuli, respectively.

The experimental protocol involved first presenting the fly, while it was at room temperature, a randomly arranged sequence of 3 different moving visual stimuli (ON, OFF and ON-OFF), each of which was displayed with one of 3 different temporal frequencies (0.4 Hz, 1.3 Hz, or 4 Hz), also selected randomly, as the trackball recorded the fly’s evoked turning responses. These frequencies indicate how quickly the visual pattern sweeps across the display windows per second. Specifically, 0.4 Hz, 1.3 Hz, and 4 Hz correspond to visual motion speeds of 30 deg/s, 90 deg/s, and 270 deg/s, respectively (**Methods**). We displayed all three stimulus-types with motion in the right-to-left and left-to-right directions^34^.

Next, we took behavioral data during thermogenetic manipulations, in which we used the temperature-sensitive *UAS-shi^ts1^*to suppress neurotransmission in the genetically targeted neurons^45^ (*i.e.*, either the T4 or T5 cells or both, targeted using a T4 driver, *R54A03-GAL4*, a T5 driver, *R42H07-GAL4*; or a T4/T5 driver, *R42F06-GAL4*; 50 female flies per group). During this part of the session, the same set of randomly ordered visual stimuli were again shown to the fly with motion in the right-to-left and left-to-right directions (**Fig. 5c**).

To determine the effect of thermogenetic perturbations, we quantified flies’ visual-induced behavior using a unitless metric, the turning index^34^, and weighted the yaw velocity of the ball according to whether the motion was in the same or opposite direction as the stimulus (**Methods**). We compared flies’ mean turning indices, averaged over the 2.25 s duration of each visual stimulus, before *vs.* after thermogenetic perturbation. At the permissive temperature (22°C), all genotypes of flies were able to follow the motion direction of ON, OFF or ON-OFF stimuli and showed increased turning indices compared to periods when there were no stimuli shown. Thermogenetic blocking of neurotransmission in the targeted neurons significantly altered the turning indices (**Fig. 5d**; *P* = 0.0338 for T4, *P* = 2.24×10^-5^ for T5, and *P* = 2.28×10^-5^ for T4+T5 neurotransmissions; *n* = 50 flies per genotype, comparing before *vs*. after blocking neurotransmission, across all visual stimulus types and frequencies; Friedman ANOVAs). After blocking T4 neurons, flies had lower average turning indices for the ON and ON-OFF stimuli but similar responses to the OFF stimulus as before thermogenetic perturbation (**Fig. 5d**; \*\**P* < 0.01 for the ON-OFF stimulus, \*\*\**P* < 0.001 for the ON stimulus; *n* = 50 flies comparing before *vs.* after blocking T4 neurotransmission; post-hoc Wilcoxon signed-rank tests with Holm-Bonferroni correction). Conversely, after blocking neurotransmission in T5 neurons, flies had lower average turning indices for the OFF and ON-OFF stimuli but responded normally to ON stimuli (**Fig. 5d**; \*\*\**P* < 0.001 for the OFF and ON-OFF stimuli; *n* = 50 flies, before *vs.* after blocking T5 neurotransmission; post-hoc Wilcoxon signed-rank tests with Holm-Bonferroni correction). When we thermogenetically blocked *both* T4 and T5 neurotransmission at the restrictive temperature (31°C), flies reduced their turning responses and had reduced mean turning index values in response to all three classes of stimuli, suggesting they were unable to detect the visual motion (**Fig. 5d**; \*\**P* < 0.01 for the ON stimulus, \*\*\**P* < 0.001 for the OFF and ON-OFF stimuli; *n* = 50 flies, comparing before *vs*. after blocking neurotransmission; post-hoc Wilcoxon signed-rank tests with Holm-Bonferroni correction). These results aligned with published findings that T4 and T5 neurons are essential for directional selectivity and motion detection and respond, respectively, to ON and OFF visual stimuli^33,34^.

Further, during experimental sessions conducted autonomously with FlyMAX, we varied the visual motion speed (temporal frequency) of the ON, OFF and ON-OFF stimuli, and tested how flies responded across different speeds. (**Fig. 5e**; \**P* < 0.05, \*\**P* < 0.01, and \*\*\**P* < 0.001 for each combination of 3 different target neurons, 3 different visual stimuli, and 3 different frequencies; *n* = 50 flies comparing before *vs.* after blocking the target neurons; post-hoc Wilcoxon signed-rank tests with Holm-Bonferroni correction). We observed an overall trend that flies showed stronger directional responses to faster-moving stimuli. Additionally, thermogenetic inactivation revealed that T4 and T5 consistently responded to ON and OFF stimuli across all frequencies, respectively.

To evaluate the automated performance of our fly behavioral assays, we quantified the time required for the entire process, from collecting the flies to performing the visual motion experiments. Using a single vial of flies for each set of experiments with the same target neurons, it took approximately 15 hours to process 50 flies, with the entire 3 sets of experiments (150 flies total) requiring about 45 hours of FlyMAX operation (**Fig. 5f**). The timing distributions for these automated experiments showed that FlyMAX spent a majority of time in the experimentation phase, *i.e.*, testing visual motion behavior (**Fig. 5g**). This suggests that, unlike conventional experiments that often have many trials for each fly, the robot could in principle perform fewer trials with each individual fly while increasing the total number of flies, potentially providing greater overall statistical power.

## Discussion

Here we introduced the Fly Manipulation and Autonomous eXperimentation (FlyMAX) system, which offers unprecedented levels of lab automation with live adult flies. The system integrates state-of-the-art deep learning-based computer vision with advanced robotic manipulation, enabling autonomous and precise handling of large numbers of individual flies for complex behavioral experiments. FlyMAX conducts these experiments in a high-throughput way without using anesthesia, enabling a potential paradigm shift in how behavioral experiments are done in research fields using *Drosophila*.

### Advancement in fly research automation

FlyMAX serves as a model for automating laboratory processes that require handling of small biological specimens that are not in a liquid state, or cultured in liquid. Traditional laboratory automation systems mainly handle specimens in liquid, transporting them using vials or pipette tips and labeling them with identifiers on their respective containers^46–48^. However, extending this type of automation to small animal species is challenging, as animal manipulation requires precise orientation and positioning of each animal subject for controlled experimentation. FlyMAX addresses this by capturing a specific part of the fly, the thorax, enabling accurate orientation and positioning using a suction tube and computerized detection algorithms. This innovation suggests a path toward lab automation for other small animal species, by adapting the suction tube dimensions and computer vision parameters to make them suited to other species.

The real-time phenotyping capabilities of FlyMAX present a unique method for classifying alert flies based on morphology. Conventional fly preparation methods using anesthesia do not require such phenotyping, since biologists usually manually perform phenotyping under a microscope. However, to achieve automation without anesthesia or human intervention, a novel approach to phenotyping becomes important. Thus, we created ‘on-the-fly’ sorting methods. Our phenotyping approach differs from other CNN-based phenotyping methods that perform object classifications on the object as a whole^49–51^. Our method for identifying key body parts for phenotyping, before running detailed classifiers, allows efficient and detailed phenotyping. The resulting precision and accuracy in fly phenotyping might also be applied to other species, perhaps even humans, based on various traits such as eye color, hair color, *etc*.

In biological research, many automated tools have emerged, yet researchers still often perform manual tasks related to setting up devices for behavior, stimulation, imaging, *etc*. Moreover, ensuring that experiments run according to standardized protocols is crucial for ensuring data quality. With FlyMAX, users are free from manually maneuvering experimental components, since the software takes control of all modules, synchronizes data collection, and streamlines the entire experimental process.

FlyMAX has the potential to enhance scientific experimentation, making it more systematic and repeatable. Consistency in experimental results is crucial, yet difficulties in replicating the results of prior publications often arise due to experiment protocols that were not fully described or unintended human biases^52–54^. By eliminating human steps in experimental setup and execution, FlyMAX should thereby reduce experimental variability caused by human factors. Further, the experimental setup procedures and settings performed by FlyMAX can be recorded digitally, facilitating the replication of experiments in the future by using the same set of digitally logged parameters.

Another advantage of FlyMAX over conventional tethering preparations is its capacity to effectively conduct experiments with male flies. Typically, in tethering experiments, researchers mainly use female flies, due to their tendency to exhibit greater vitality and quicker recovery from anesthesia^55,56^. Our system accommodates flies of both sexes, enabling a broader range of experimental possibilities, such as the exploration of subtle behavioral differences between the sexes. For instance, we observed that male flies seemed more sensitive to visual motion than females, which might potentially be obscured by the use of anesthesia (**Fig. 2e-g**).

### Comparison to current tools

FlyMAX not only significantly improves fly collection over our previous version but also expands the capabilities for automatic experimentation^27^. Our prior robot excelled in picking alert flies without anesthesia, but had limited versatility post-collection. Biologists had to manually control the robot and repeatedly swap the collection stage with other rigs for each experiment. This involved repeated manual calibrations, as the position of the stage varied with each swapping. FlyMAX offloads these manual tasks, allowing biologists to focus on other tasks.

Compared to other current tools, FlyMAX stands out as an autonomous system for sophisticated experiments. MAPLE, for instance, used a similar gantry with handling tools based on air suction^28^. Its capturing method is to enclose a fly inside a pipette tube. Regardless of health concerns, which the paper does not fully investigate, the system cannot fully manipulate the flies for performing experiments^28^ but instead merely transports fly subjects from one place to another. While this is useful for automation, biologists can transport flies without using anesthesia. MAPLE cannot automatically perform tasks that normally require anesthesia for manual handling, such as experiments with tethered flies. By comparison, FlyPEZ was designed to help biologists capture natural fly behavior^31^. It uses open space and the geotactic behavior of flies to bring the fly to the imaging stage. Because the fly on the stage is not captive, detailed experiments or behavioral evaluations remain challenging. Moreover, controlling its environment is difficult, and once it flies away, no further imaging or behavior analysis is possible.

### Limitations and mitigation strategies

FlyMAX effectively leverages the geotactic behavior of flies, using it to naturally guide the flies to the arena at the collection station. However, for certain fly lines that are intentionally made unhealthy, or for mutants with geotactic deficits, prompt fly collection may not be feasible, as these flies cannot climb to the arena. In such cases, we recommend expressing genes that impair normal behavior only under specific experimental conditions, such as with thermogenetics and optogenetics, to ensure normal behavior initially while the flies are undergoing robotic picking and handling. We installed a temperature control at our experimental station for thermogenetic manipulations, as needed.

Another limitation regarding the collection of flies is that the collection stage is in an open space, which does not prevent flies from escaping. While the number of flies used for experiments is typically abundant, prepared flies, such as those that have undergone prior procedures like microsurgery, have significant experimental value, since each preparation takes considerable time and labor. As FlyMAX does not confine the flies, biologists might have concerns about losing these flies during the collection process. To address this, the robot could be encapsulated, or one could render the flies unable to escape by cutting their wings during microsurgery. Alternatively, in the future, FlyMAX might be directly integrated with a microsurgery station^57^.

### Technological outlook

Equipped with a gantry enabling long-distance translation, FlyMAX offers future possibilities for adding more experimental stations with additional capabilities, such as for brain imaging. Freely moving flies tethered by the thorax pose a challenge for brain imaging due to the flexibility of their head movements. Preparing flies through microsurgery and gluing their heads to the thorax beforehand might allow imaging and long-term studies, including with brain imaging^36^. Another possibility might be to integrate FlyMAX with fly rearing, such as with fly-flipping robots^58^. This integration with our phenotyping feature could potentially further automate genetics.

FlyMAX’s capacity for large-scale experimentation holds promise for applications in drug screening and reverse genetics. For instance, studies of a fly model of Alzheimer’s disease might involve labeling neurons responsible for specific behaviors, systematically permuting possible neural gene knockout combinations, and then examining all the combinations with FlyMAX^59,60^. Additionally, one might screen insecticides or insect repellents in an automated way, potentially gaining insights into environmental or agricultural issues related to insects^61,62^.

In the future, the application of deep reinforcement learning algorithms to optimize the robot’s initial positioning of the fly for behavioral experiments may help to preclude human variability or errors. Robot training via reinforcement learning procedures might allow one to train FlyMAX to maximize a tethered fly’s range of movement^63–65^. By optimizing parameters such as the roll, pitch and distance from the fly’s body to the rig, the system can potentially achieve precise positioning without human bias or intervention, leading to fully reproducible, robotically standardized experimental protocols.

FlyMAX’s versatility probably extends beyond *Drosophila*, opening new experimental possibilities with a range of insect species. One could customize the vacuum end-effector’s power and use species-specific template images for handling, for gentle manipulations of other insect species without health impairments. This approach might enable individualized experiments with a variety of insect models.

Overall, the FlyMAX system streamlines the process of conducting environment-controlled behavior experiments on *Drosophila* and serves as a versatile platform with a potential to reshape laboratory automation for insects and other small model organisms.

---

**Fig. S1.**
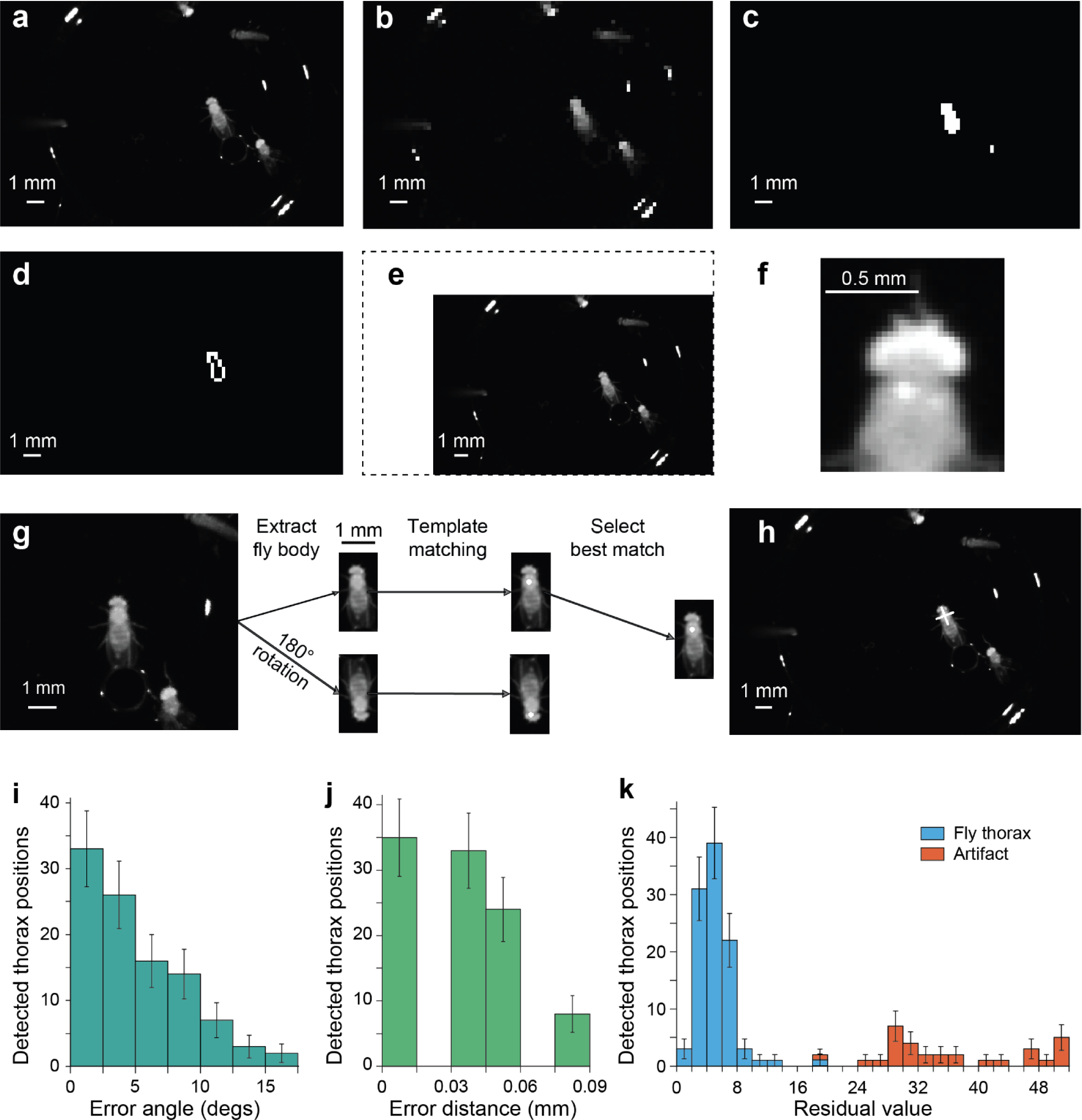
Fast, scale- and rotation-invariant template matching for thorax detection. **a–h)** Fly thorax detection algorithm that estimates the fly thorax position. From the original image taken from the platform camera at the delta robot (**a**), the algorithm downsampled the image for faster machine vision (**b**). Then, it performs Otsu’s automatic thresholding to binarize the image between matched fly pixels and background (**c**). Then, it performs blob detection to identify fly bodies and selects the largest detected blob as a target fly (**d**). Once the computer vision finds the target, it scales the original image, (**a**), based on its blob size to match the standard blob of the template (**e**). The dashed outline indicates the original size of the image. The template for matching contains a head and a thorax of a fly which are not much variant among different genders and/or phenotypes of flies (**f**). In the template, the algorithm utilizes a predefined position of the center of the thorax to estimate the target position based on the template matching result. As it can compute the orientation of the detected fly in (**e)**, it can rotate the cropped part for template matching (**g**). Because the computer vision cannot accurately identify whether head orientation aligns with that of the template in (**e)**, it rotates the image 180 degrees to include the case when the fly’s head is downwards. The template matching computes residual values and determines the thorax position based on the lowest value (**h**). **i)** The orientations estimated at (**d**) are compared to manually estimated orientations from (**a**). *n* = 101. Error bars: s.d. calculated as 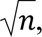 where *n* is the number of observations in each histogram bin. **j)** The estimated positions at (**h**) are compared to manually annotated positions from (**a**). *n* = 100. Error bars: s.d. calculated as 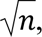 where *n* is the number of observations in each histogram bin. **k)** Distribution of residual values of fly thoraxes and artifacts. Artifacts were light reflections, unexpected materials, or image modifications. *n* = 101 for fly thoraxes, and *n* = 31 for artifacts. Error bars: s.d. calculated as 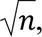 where *n* is the number of observations in each histogram bin.

**Fig. S2.**
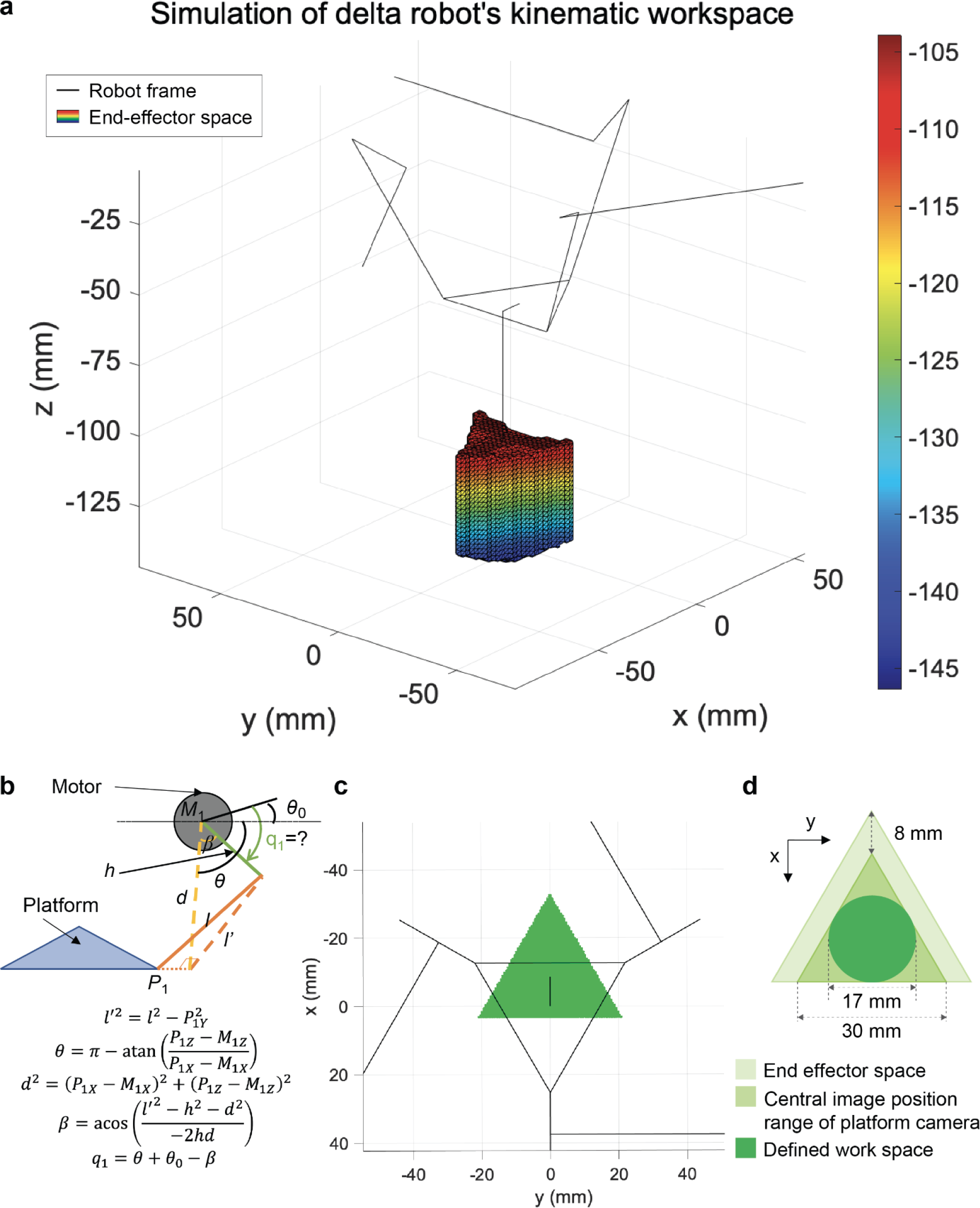
Simulations of the delta robot’s kinematic workspace. **a)** Simulation of delta robotics kinematics. The dimensions of the delta robotic frame determine the trajectories of the end effector. We aimed to have longer Z translation among XYZ translations to facilitate deeper robotics manipulation for experimental purposes. The mesh in the diagram displays the trajectories of the end effector based on the dimensions of the robotic frame. We calibrate the kinematics of the end effector with the physical mechanism, ensuring that the simulated movements aligned accurately with the actual robot’s movements. The color bar indicates the Z depth of the end effector, while the black line frame indicates the robotic components. **b)** Delta robot inverse kinematics. The task of the inverse kinematic algorithm is to compute the actuator shaft angles *q*_1_ to position the picking needle at the desired XYZ location. Relying on the delta mechanism to enforce pure translation of the picking platform and selecting actuator 1 for explication, we first translate the desired needle position to the associated platform joint *P1*. Projecting this point into the plane swept by the actuator’s radial arm, (*l* -> *l’*), allows solving the projected triangle for the actuator angle. Repeating this calculation for the other actuators gives the required actuator angles for the desired position. **c)** Top-down view of the space of the end effector spanning on XY space. The black lines indicate the robot frame and the green shape indicates the XY workspace of the end effector. The black line indicates the position of the end effector from the center of the platform. Notice the trajectory forms the inverted triangular shape of the platform whose center is on the end effector position when the platform is homed. **d)** The working XY space computation calculated from (**c**). The ‘defined workspace’ shows the space that the end effector can safely reach. This computation takes the needled position and camera space into account to deduce the circular space for the fly collection station.

**Fig. S3.**
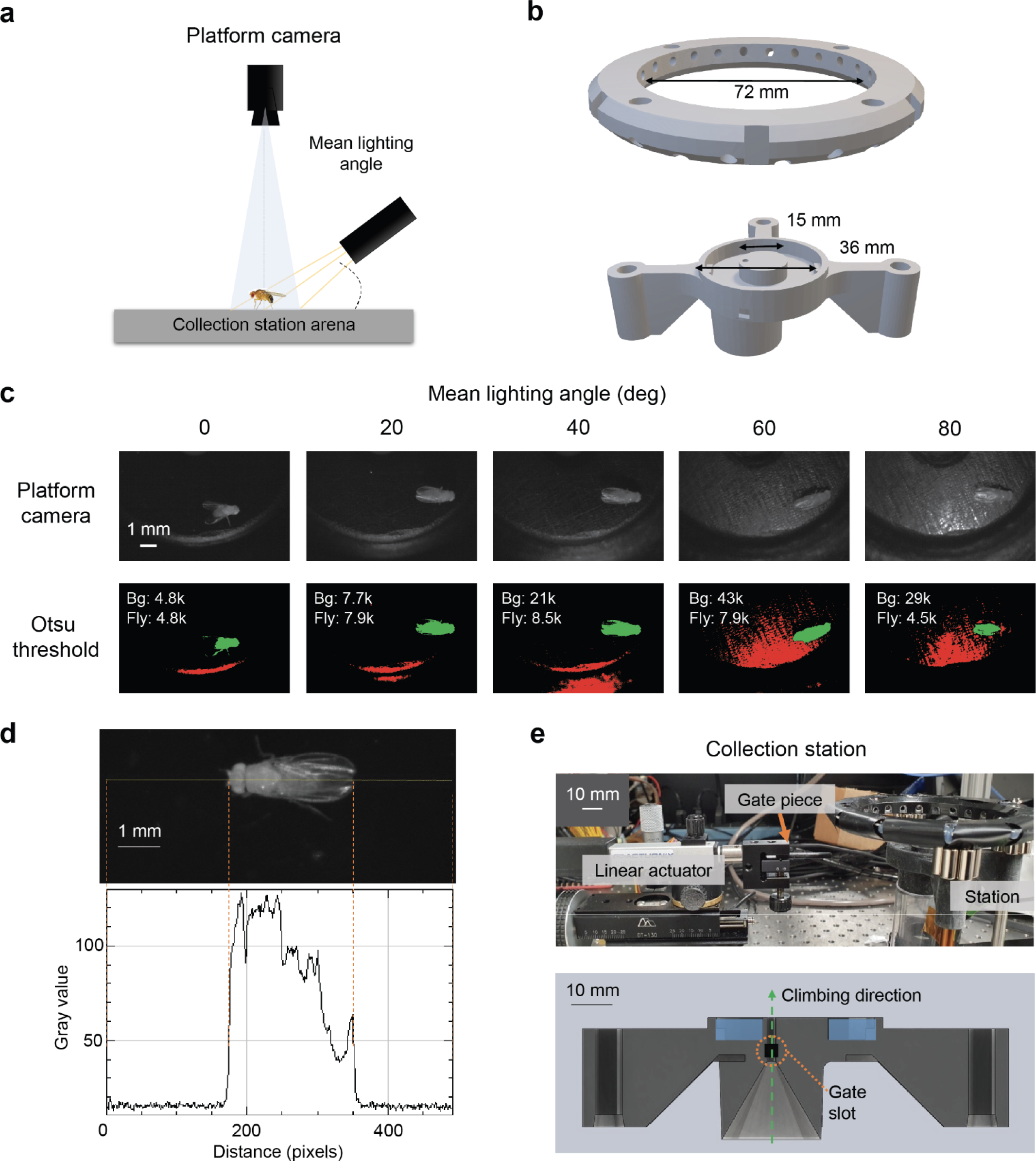
Collection station design for providing good contrast lighting for machine vision. **a)** Setup for controlling the angle of illumination. To get a top-down view image from the platform camera of the delta robot, we engineered the lighting angle of infrared illumination. We varied the lighting angle to determine the desired angle that can make the image of a fly salient compared to the background. **b)** CAD design for fly collection station. *Top*. CAD design for the fly collection station. We incorporated a custom ring light setup to illuminate the arena. Based on the lighting setup in (a), we developed a ring-structured lighting setup to provide uniform light distribution. It was essential to ensure that the frame did not interfere with the moving parts of the end effector. *Bottom*. The custom arena design for the collection station. Based on trajectory information derived from the simulation (Fig. S2d), we configured the arena space with a diameter of 15mm. We surrounded the arena with a sunken area dedicated to water filling. Beneath the arena lies a slot for a regular vial, and a hole in the arena serves as a passage for flies to climb up. **c)** Contrast checking. We adjusted the angles to optimize the lighting for illuminating the stage. Placing a fly on the arena, we varied the angles to assess their impact. To evaluate the contrast, we used the Otsu method to threshold the images captured by the platform camera. We quantified the number of high-valued pixels for both the fly and the background to determine the proportion of fly pixels and the ratio between high-value fly and background pixels. In the images, ‘Bg’ represents the number of high-valued pixels for the background (colored in red), while ‘Fly’ represents the number of high-valued pixels for a fly (colored in green). **d)** An example of cross-sectional pixel values of a fly image with good contrast. Compared to the background, most of the fly body parts register values above 50, while background pixels remain close to zero. The lighting angle for this image is 20 degrees. **e)** *Top*, The software controls a linear actuator holding a gate piece to manage the gate at the collection stage. This mechanism closes or opens the passage, ensuring that not all the flies exit the vial. *Bottom*, A sagittal view of the collection stage. The green dotted line shows a passage for flies with gate position. The gate slot indicates the space blocked by the gate piece when closed.

**Fig. S4.**
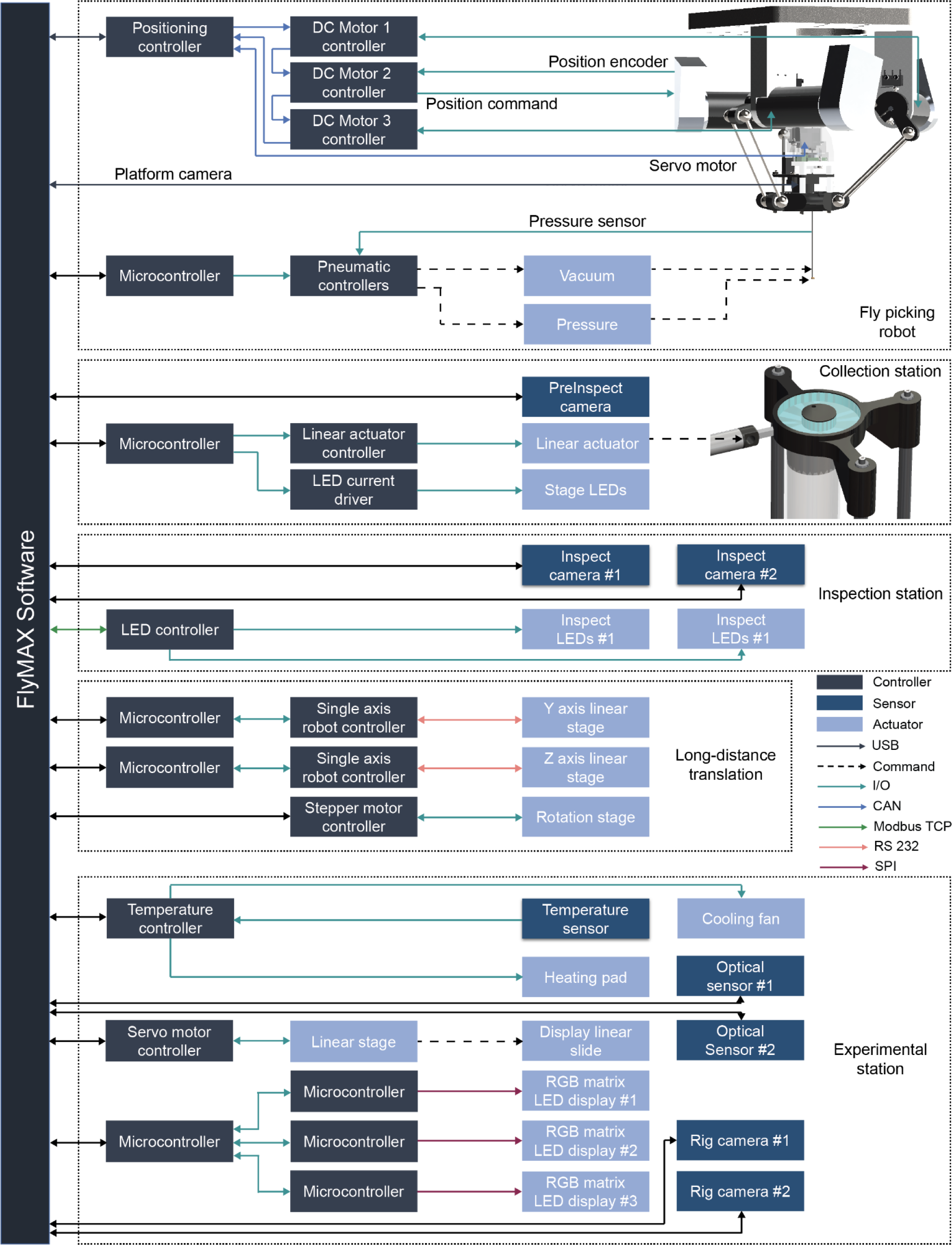
Software/hardware communications in FlyMAX. Communications between hardware and software. We used different protocols to confirm seamless communication between the software and individual devices. *Fly picking robot*. Three DC motors configuring the delta robot are communicating via the main Maxon EPOS position controller with daisy-chained CAN network. Each motor has a dedicated controller for high accuracy and speed. The system sends position commands to each motor controller, and all motors simultaneously perform the commands upon receiving the execution signal. The motor for rotating the yaw of the end effector is also connected to the main EPOS position controller. Each controller reads position data from its encoder and commands the position. With the inverse kinematics of XYZ position to angular rotations of motors, the robot can translate to desirable position with accuracy (See Picking robot in Methods). The end effector is a hypodermic tube that can control pneumatics. With the pneumatic setup, it can perform vacuuming and pressuring. *Collection station*. We installed a pre-inspection camera to perform machine visioning tasks at the collection station. Other communications related to the collection station are lighting and gate controlling. The main software commands the LED current driver to turn the lights on or off and to change their intensities. For gate controlling, a linear actuator translates to control the openness of the passage from vial to the stage. *Inspection station*. The inspection station consists of lighting and cameras for high-level inspection. The LED driver supports a C++ library, enabling direct communication with FlyMAX software to configure detailed lighting options such as specific intensity and duration of illumination. *Long-distance translation*. FlyMAX supports long-distance translation via communicating with linear stages. Through microcontrollers (Arduino Mega2560 R3), FlyMAX software directly writes commands to writers. These support long Y and Z translations. The gantry also has a rotation stage which can rotate the picked fly to users’ desirable angles. *Experiment station*. For behavior assay, we installed a temperature controller, locomotion tracker and display panels. For temperature control, we used an infrared remote temperature sensor to estimate the body temperature of the fly running experiments. We used Teensy v4.1 for this process for low latency in temperature controlling. We used a thermistor as a heating pad and a fan for cooling. The microcontroller accepts the data from the sensor to compute the magnitude of the heater or cooler to maintain the desirable temperature. To facilitate full automatic experimentation, we installed a linear stage to translate one of the panels to allow two perpendicular view points for machine visioning for detailed positioning. We used two rig cameras for this purpose. We used two optical sensors to quantify the locomotive behavior of a fly walking on an air-suspended ball. We used a high-performing microcontroller (Teensy v4.1) for controlling the display to keep a high refresh rate. Each microcontroller controls the switch of individual LED pixels to control the light stimuli.

**Fig. S5.**
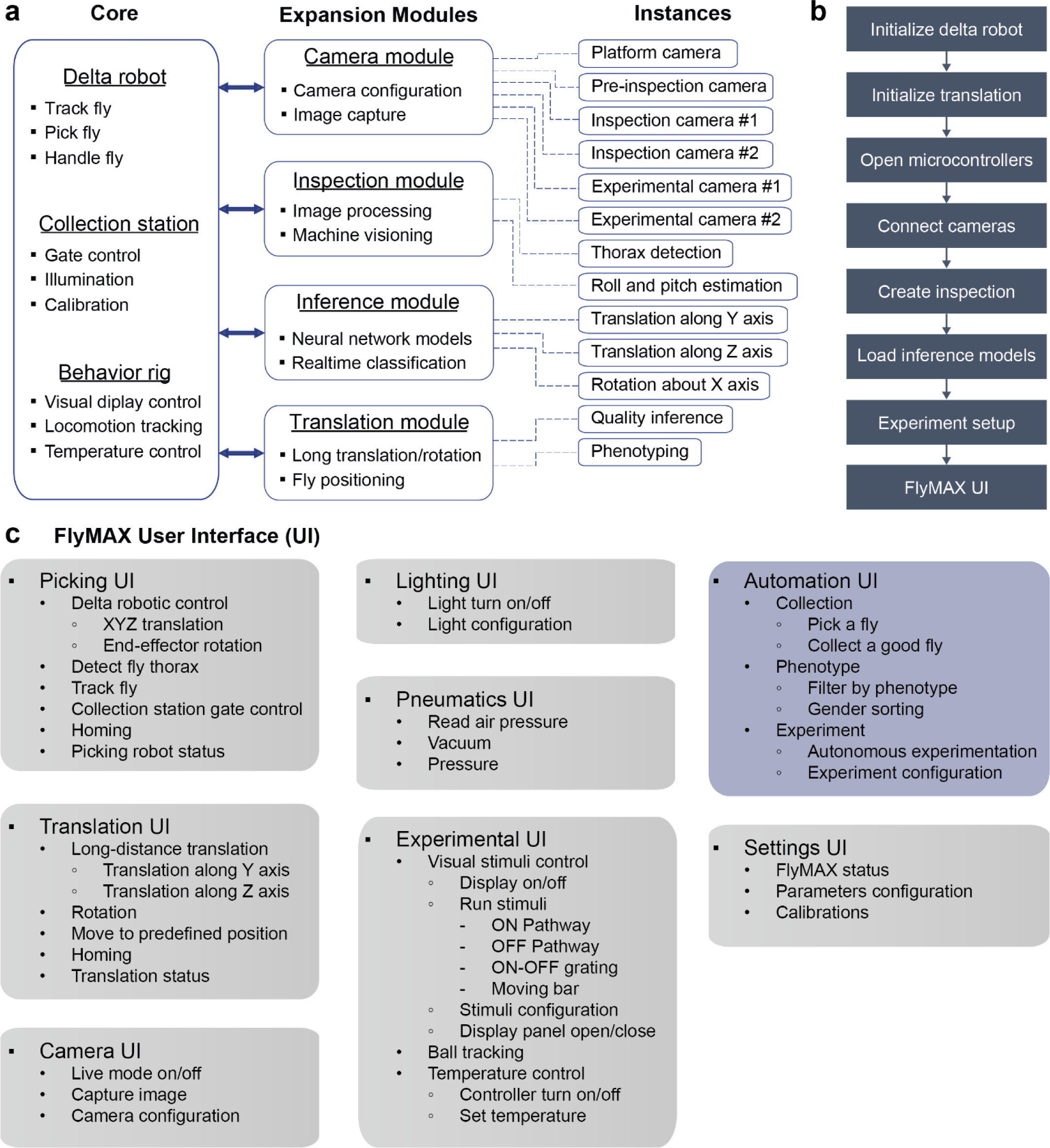
FlyMAX software. **a)** Software architecture of FlyMAX. We implemented core parts in the main software application, expansion modules are dynamically linked libraries that support the core program to create instances at initialization. The core components comprise of delta robotics, collection station, and behavior rig. For the delta robot, it has fly thorax detection, picking, and manipulation. The thorax detection software has methods for detecting a thorax target of a fly. This frame calls subordinate functions that are responsible for capturing image from the camera at delta robot, performing image processing and machine visioning to locate the target. The picking methods are robotic control functions that commands motors to schedule fast motion and execute. The manipulation part also contains fine robotic control and pneumatics control to suck or push air flow. The collection station has methods related to controlling the stage for the delta robot to capture flies. The software sends signals to a linear actuator to control the gate of the stage. It also has a lighting command that turns on/off the infrared light of the stage. It also supports functions necessary for calibration. This calibration consists of calibrating Z height of the end effector, image space from the camera at delta robot, stage space available for picking, and so on. For the behavior rig, we built a setup for visual sensory motor behavior. For this, the software has methods for controlling display that gives visual stimuli/cues to a subject. The software also has methods that read locomotive data received from infrared optical sensors that read ball movements. It also has functions that read and write the environment temperature. The expansion modules consist of camera, inspection, translation, and inference modules. We implemented them in C++ libraries to make the software modularized and easy to develop and maintain. Also, each library works as a class such that multiple instances that share the same functionalities such as cameras, can be implemented with ease. The camera module is responsible for configuring cameras and capturing images. The camera instances used for FlyMAX are one located at the platform of delta robot, one camera for pre-inspection located at collection station, two cameras for inspection, and two cameras for experimental station. The inspection module performs image processing and machine visioning on the images captured from the camera modules. Translation modules are responsible for long-distance translations and rotation. Currently, FlyMAX supports Y and Z axes translations and one rotation for pitch control. This module is also responsible for fine positioning fly to the behavior rig with the resolution of 0.01mm in translation. The inference module is similar to the inspection module, but this module is mainly for feedforwarding image input to neural network models. This supports quality inference and phenotyping. **b)** Initialization workflow when FlyMAX program starts. All the initialization steps confirm if any communication or device has any error before proceeding. At first, the picking robot initializes by translating motors for the delta robot to homing positions. Also, the rotating motor for the end effector is initialized to 0 degree. These steps make sure the robot always starts at a consistent position. Then, the translation module home to the inspection station. This ensures that FlyMAX knows the position of the delta robot as well as to make sure the robot does not collide with other objects. Once all motors are confirmed, the software opens communication channels for other microcontrollers that are responsible for controlling peripherals such as temperature and pressure sensors, lighting controllers, and display controllers. Afterwards, FlyMAX initiates all cameras; it connects the cameras to check its status and then turns off the live mode for the bandwidth purpose. Then, FlyMAX creates and loads the inspection modules and neural network models sequentially. The system initializes experimental setup by turning off the temperature control and stimuli display initially. FlyMAX checks the locomotion tracking status and turns off the live tracking mode. If the system completes all initializations, the program transitions to a user interface for controlling and commanding the robot. **c)** FlyMAX offers a comprehensive suite of automated tasks, including fly collection, phenotyping, and experimentation. It also provides a fine-grained UI for robot-assisted control. The Translation UI facilitates XZ linear translation and rotation around the X-axis. The ‘Picking UI’ supports the delta robotic control, detecting a fly and its thorax, tracking a target fly in real time, and controlling the gate of the collection station. The Camera UI supports configuration camera parameters and image capturing. Users can control all the lights via the Lighting UI. The Pneumatics UI is for controlling air flow of the end effector. The Experiment UI consists of controlling display, ball tracking and temperature for behavior assays. The major automated tasks are collection, phenotyping, and experimentation. From the panel, one can see that these autonomous features employ all the underlying functions supported by other UIs. The UI also supports displaying the status of all components of FlyMAX, setting its parameters, and calibrating the system.

**Fig. S6.**
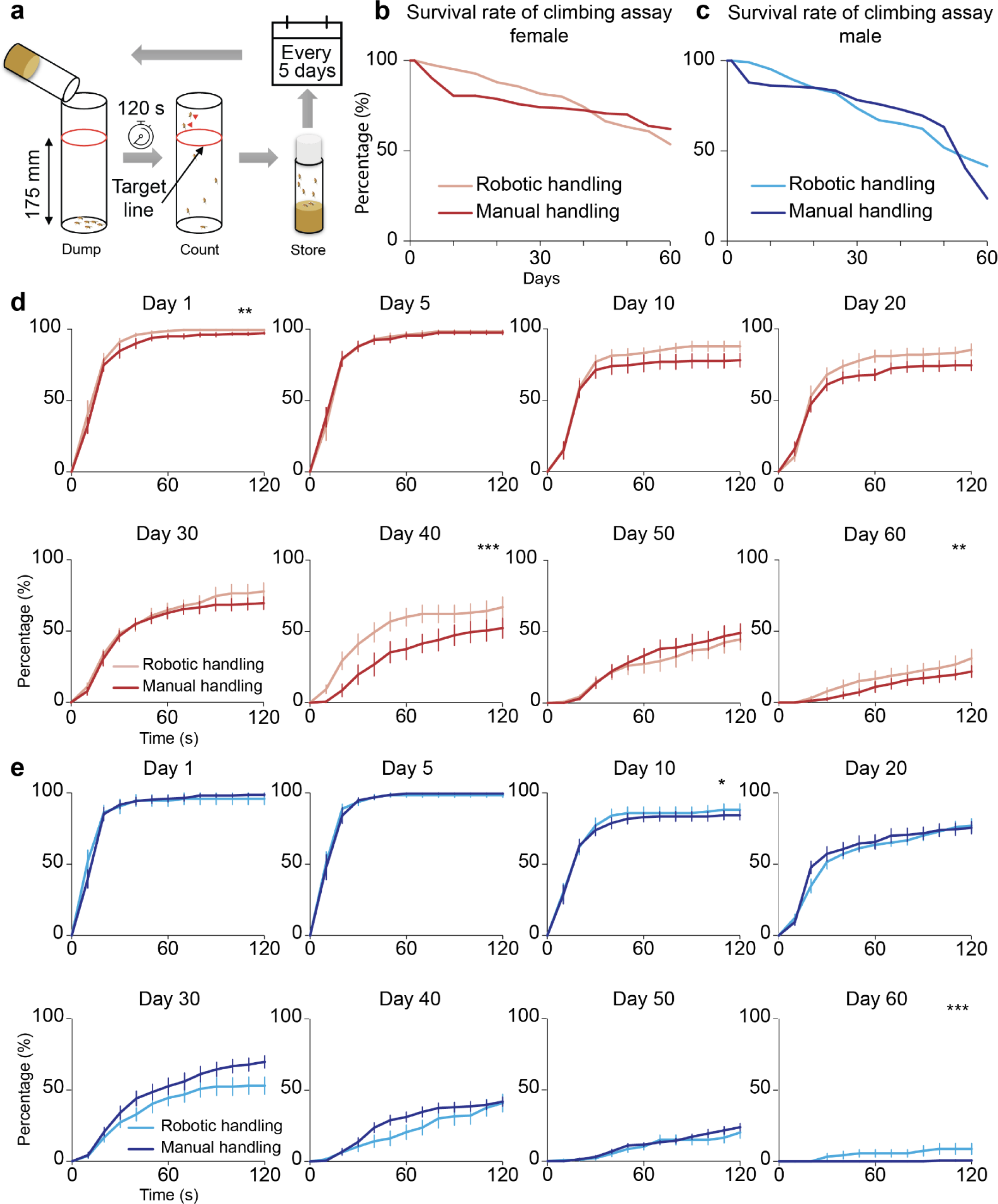
Longitudinal studies of fly climbing abilities. **a)** Protocol for longitudinal climbing assay. For each assay, we tapped a fly vial on the ground to ensure that flies remained on the ground. We set the target line at 175mm to count flies that crossed over it during the 120-second duration of each assay. Unlike conventional climbing assays, where biologists sacrifice subjects after the test, we kept flies after the experiments to assess their performance over longer periods. To allow for recovery time, we repeated the experiments with a five-day interval. **b, c)** Survival rate of the flies performed longitudinal climbing assay. The graphs depict the survival rate of the flies following the longitudinal climbing assay. On Day 0, FlyMAX captured the robotically handled flies when they climbed to the collection station and then sorted them by gender, whereas we exposed manually handled flies to CO2 and then sorted them. These graphs illustrate the percentage of flies that performed on each day compared to the number of flies experimented on Day 0. For (**b**), the total number of robot-handled flies on Day 0 was 125, and for control flies, it was 174 (*P* = 0.38; Kruskal-Wallis ANOVA). For (**c**), the total number of robot-handled flies on Day 0 was 106, and for control flies, it was 174 (*P* = 0.85; Kruskal-Wallis ANOVA). **d, e)** Longitudinal climbing assay of (**a**) female and (**b**) male flies that is presented in Fig. 2cd on a daily basis. Each day, we calculated the percentage of flies that crossed the target line (175 mm) by dividing the number of successful flies by the total number of surviving flies on that day, as described in (**b**) and (**c**). For (**d**), we used 8 vials with a total of 125 female flies in the robotically handled group and 9 vials with a total of 154 female flies in the manually handled group. Error bars: s.e.m. across 8 vials and 9 vials for robotically and manually handled groups, respectively. For (**e**), we used 8 vials with a total of 106 male flies in the robotically handled group and 9 vials with a total of 114 male flies in the manually handled group. (\**P* < 0.05, \*\**P* < 0.01, \*\*\**P* < 0.001; Kruskal-Wallis ANOVA). Error bars: s.e.m. across 8 vials and 9 vials for robotically and manually handled groups, respectively.

**Fig. S7.**
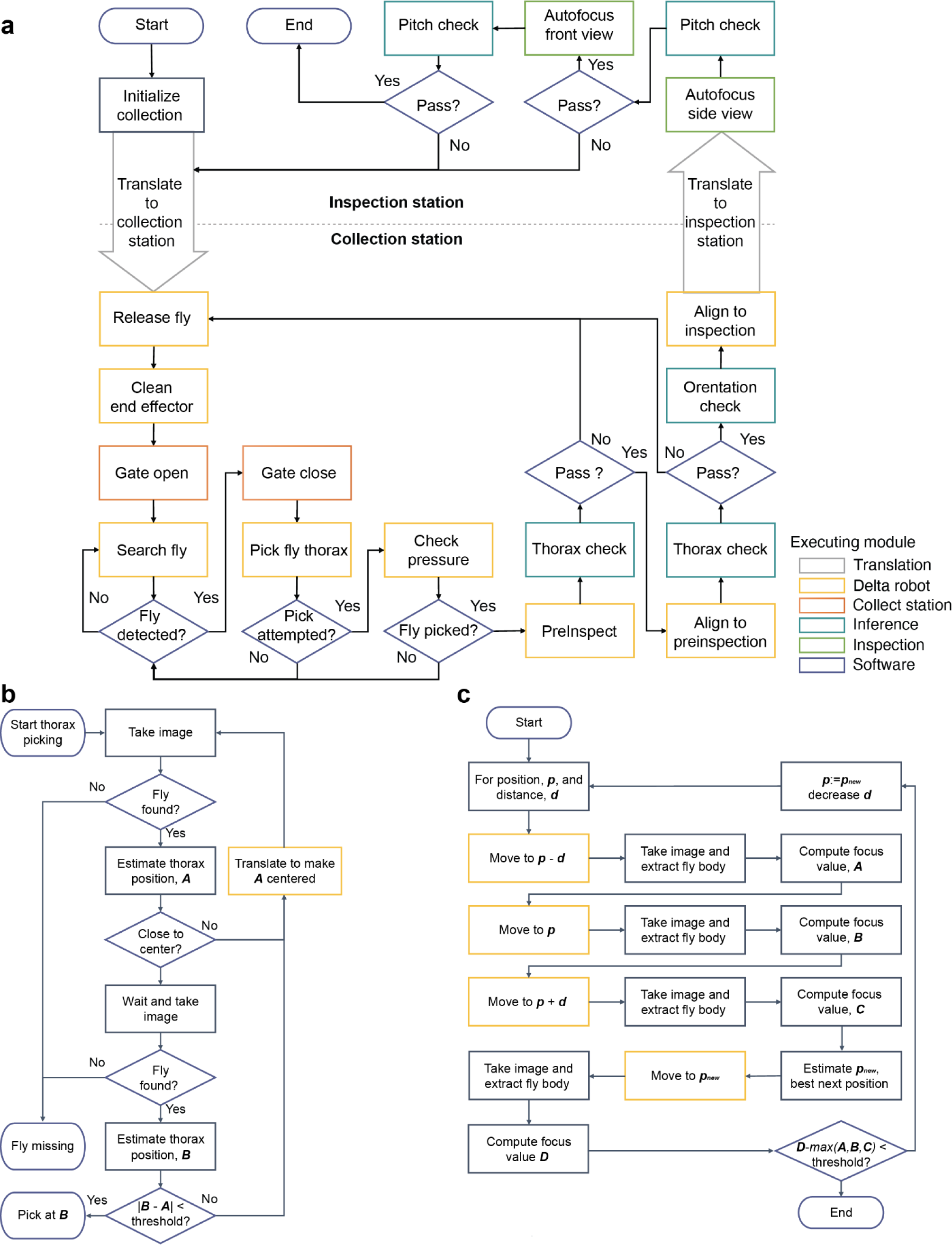
Fly collection workflow. **a)** Once FlyMAX is initialized as described in **Fig. S5b**, users can select to collect a good fly in the Automation UI diagrammed in **Fig. S5c**. Initially, the robot prepares for collection. After FlyMAX initializes, the delta robot positions itself in the inspection station. To begin collection, the robot moves to the collection station. It then performs the release of any previously held fly and proceeds with cleaning and checking the air pressure of the end effector. This step ensures consistent air pressure when performing vacuuming. Subsequently, it opens the gate of the collection station to allow flies to climb up to the arena. Using the platform camera, the robot checks if there are any flies in the arena. Upon detection, the gate closes, and the robot proceeds with picking the detected fly by performing thorax detection and fast approaching the target. After the attempted picking, the robot initially checks the air pressure sensor to verify if it has picked up a fly and further confirms this with the pre-inspection camera. During this step, machine visioning checks if the robot has appropriately picked up the thorax, and depending on the inference result, it either repeats the initial picking process or proceeds to the next step. Once the robot aligns the fly with the pre-inspection camera, it performs another thorax check as a double check. If pre-inspection passes, the robot moves on to the inspection station for further checking. it checks the fly’s orientation to ensure the head or tail faces the camera correctly and rotates the fly accordingly. Using the side view camera, it auto-focuses to obtain a crisp image (as described in (**c**)) and then assesses pitch quality. Subsequently, with the front view camera, it auto-focuses again to obtain a crisp image and estimates roll quality. If both criteria are met, FlyMAX confirms that it has collected a good fly. **b)** Fly thorax detection algorithm performed ‘Pick fly thorax’ process in (**a**). The Fly thorax detection algorithm enhances accuracy by translating the platform position consistently for pick attempts. It also verifies if the target fly is stationary before attempting to pick it. To illustrate, it captures an image from the platform camera and utilizes machine visioning to check if a fly is present in the arena. If found, thorax detection begins. Once computed, it checks if the first detected position, detonated as ***A***, is close to the camera center. If not, the delta robot moves to bring it closer to the center. This ensures a relatively uniform trajectory for fast and optimized movement. After locating ***A***, it waits for 50ms to determine if the captured fly is stationary or in motion. Subsequently, FlyMAX takes another image and measures the newly estimated thorax position, denoted as ***B***. If the distance between ***B*** and ***A*** is within ten pixels, it considers the fly stationary and begins the picking attempt. **c)** Autofocusing algorithm at the inspection stage. Compared to conventional autofocusing methods where the lens in a camera translates to obtain a crisp image, the end effector of the delta robot translates to perform autofocusing. To illustrate, FlyMAX takes three images with a positional difference of *d*. It computes the focus value for each captured image using the difference of Gaussians method. Based on these computed values, it computes the next probable position for achieving the best crisp image and updates the new position accordingly. This iterative process continues with smaller d increments until the change in focus value becomes subtle.

**Fig. S8.**
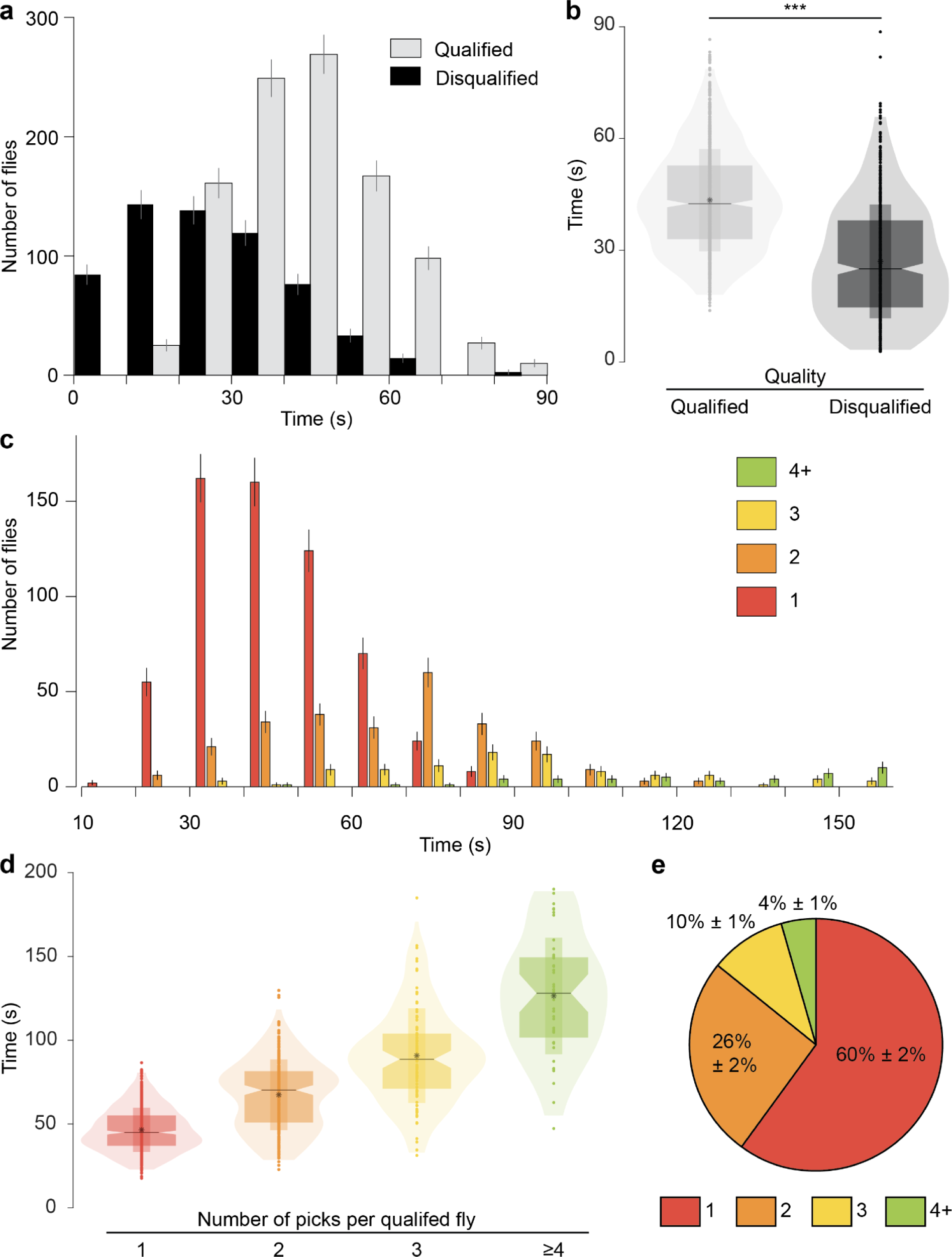
Characterizations of fly collection. **a)** Distribution of the time taken for automated fly collection, as illustrated in Fig. 3. A qualified fly has its thorax correctly picked up within a good range of roll and pitch angles. We consider flies meeting this criterion as qualified, while we classify those not meeting these conditions as disqualified (refer to the “Picked Quality Checking" section in Methods for detailed criteria). Distribution of time taken to pick qualified and disqualified flies during automated fly collection performed in Fig. 3. Error bars: s.d. calculated as 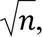 where *n* is the number of flies in each histogram bin. **b)** Violin plot of (**a**) shows the distribution of time spent in picking qualified and disqualified flies, with kernel density representing the timing spread. Boxes span the 25th-75th percentiles, whiskers span 1.5 times the interquartile distance, horizontal lines denote median values, notches reflect the 95% confidence interval for the median. Asterisks denote mean values, narrow bars indicate the standard deviation from the means, and dots represent individual data points. This description applies to all the violin plots in this paper. (\*\*\**P* < 0.001; Wilcoxon rank sum test). **c)** Time distribution of time taken to pick qualified flies. Numbers in the labels indicate the number of pick tries until to collect qualified flies. Error bars: s.d. calculated as 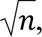 where *n* is the number of flies in each histogram bin. **d)** Violin plot of (**c**) shows the distribution of time spent picking qualified flies, categorized by the number of picks per qualified fly. Note the average time difference between each number of picking tries is linearly proportional to the average time for picking disqualified flies. **e)** Pie chart of (**c**). This chart displays the proportion of how many pick attempts the robot made to collect a qualified fly. Error bars: s.d. calculated as 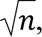 and then converted to a percentage, where *n* is the number of observations in each pie slice.

**Fig. S9.**
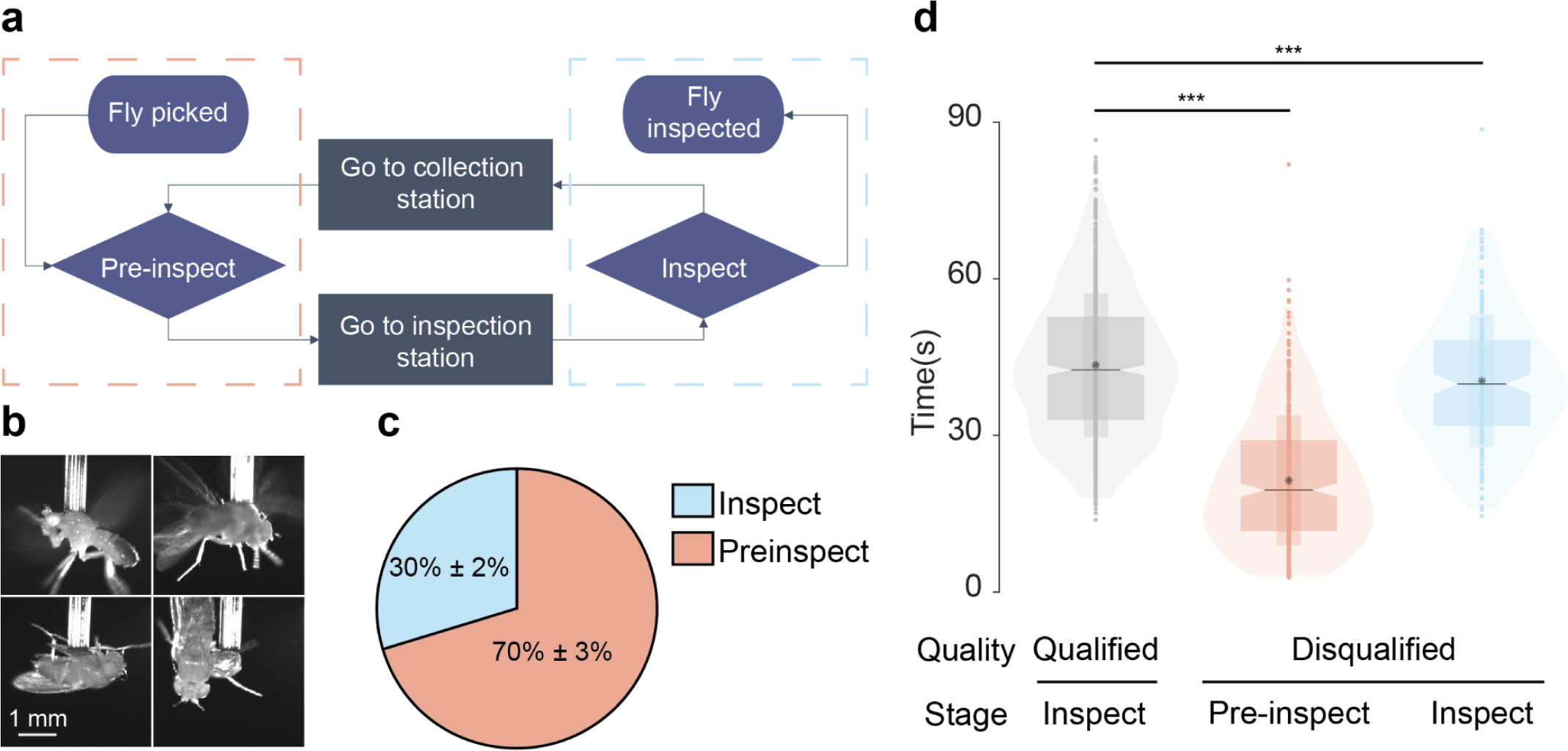
The use of pre-inspection at the collection station reduced the net time spent handling disqualified flies. **a)** A diagram showing the characteristics of pre-inspection features compared to inspection features. The dashed pink line indicates the collection station, and the dashed blue line indicates the inspection station. Note that the pre-inspection station resides physically at the collection station. This setup ensures that FlyMAX filters out a disqualified fly during the pre-inspection process. This avoids two translation processes for inspection, improving execution timing. **b)** Sample images taken by the pre-inspection camera. *Top rows*, Seemingly well thorax picked flies. FlyMAX will inspect these flies further at the inspection stage. *Bottom rows*, FlyMAX disqualified badly picked flies and discarded these flies without proceeding to the inspection stage. **c)** A pie chart illustrating the percentage of stations that filtered out disqualified flies. The total number of disqualified flies assessed was 608. Error bars: s.d. calculated as 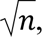 and then converted to a percentage, where *n* is the number of observations in each pie slice. **d)** Violin plot showing the distribution of time taken to disqualify flies at both pre-inspection and inspection stations. Pre-inspection significantly reduces the time spent determining disqualified flies, as indicated by the lower time distribution compared to the inspection station. This reduction suggests that pre-inspection effectively minimizes the total automation runtime by filtering out badly picked flies earlier in the process. (\*\*\**P* < 0.001; Kruskal-Wallis ANOVA with post-hoc Holm-Bonferroni correction with pairwise comparison).

**Fig. S10.**
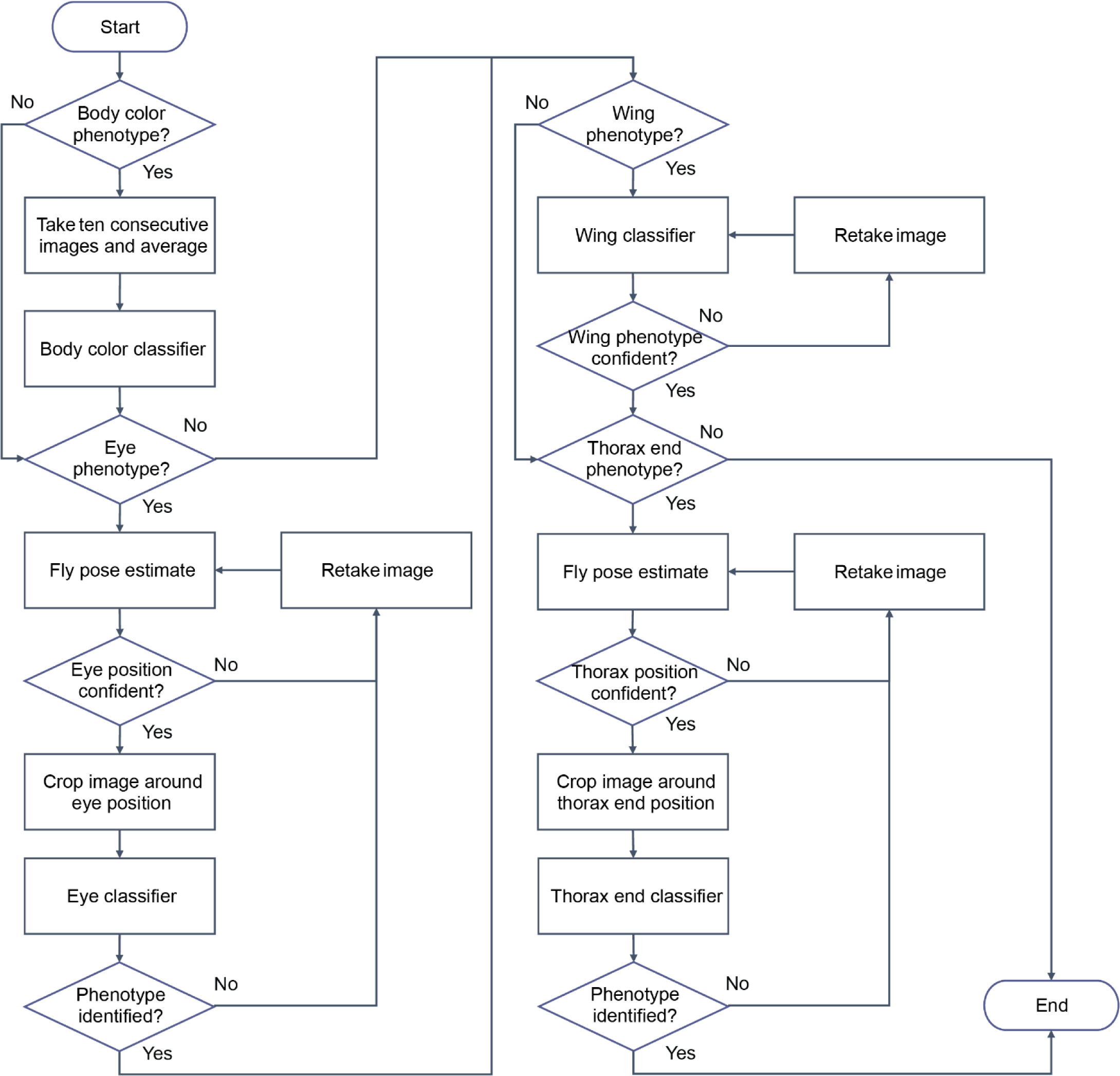
Phenotyping workflow. FlyMAX’s phenotyping workflow allows for the inclusion or exclusion of phenotyping steps for specific body parts based on research needs. *Body color.* The workflow initiates with the analysis of body color. The process begins by inputting downsampled images into the system for initial inference. To enhance reliability, the system computes the average values from ten sequential images. This approach helps mitigate variations and ensures a more stable phenotyping base for body color determination. *Eye phenotype*. The phenotyping process for the eyes employs pose estimation to pinpoint the precise location of keypoints associated with the eyes. The neural network not only identifies these keypoints but also provides a confidence level for each, reflecting the clarity and visibility of the region of interest. Challenges such as blurring or occlusion due to the dynamic nature of live flies may impact these confidence levels. The system iteratively takes new images and refines the pose estimation as needed, based on the confidence scores, until it achieves a satisfactory level of precision. For the actual classification, it analyzes a high-resolution crop focusing on the eye region to determine the specific eye phenotype. *Wing*. For wing analysis, the workflow utilizes the full image to assess wing phenotypes. Given that flies are occasionally captured in flight by the system, the clarity of the wing images can vary significantly, from crisp shape to blurred outlines. The system iteratively captures and analyzes images until it can make a clear inference, ensuring accurate phenotyping despite the challenges posed by the motion of flying. Subscutellar bristles. Similar to the eye classification process, thorax phenotyping involves capturing images and performing pose estimation to accurately locate the thorax endpoints. The system then uses a cropped, full-resolution image of the thorax area for detailed classification. The keypoint that marks the thorax position is crucial for this step, as it guides the system in focusing the analysis on the relevant area for precise phenotype classification. We engineered this phenotyping workflow to adapt to the dynamic and complex nature of live Drosophila specimens. It employs a combination of advanced imaging techniques, iterative analysis, and machine learning to ensure accurate and reliable phenotyping across various body parts.

**Fig. S11.**
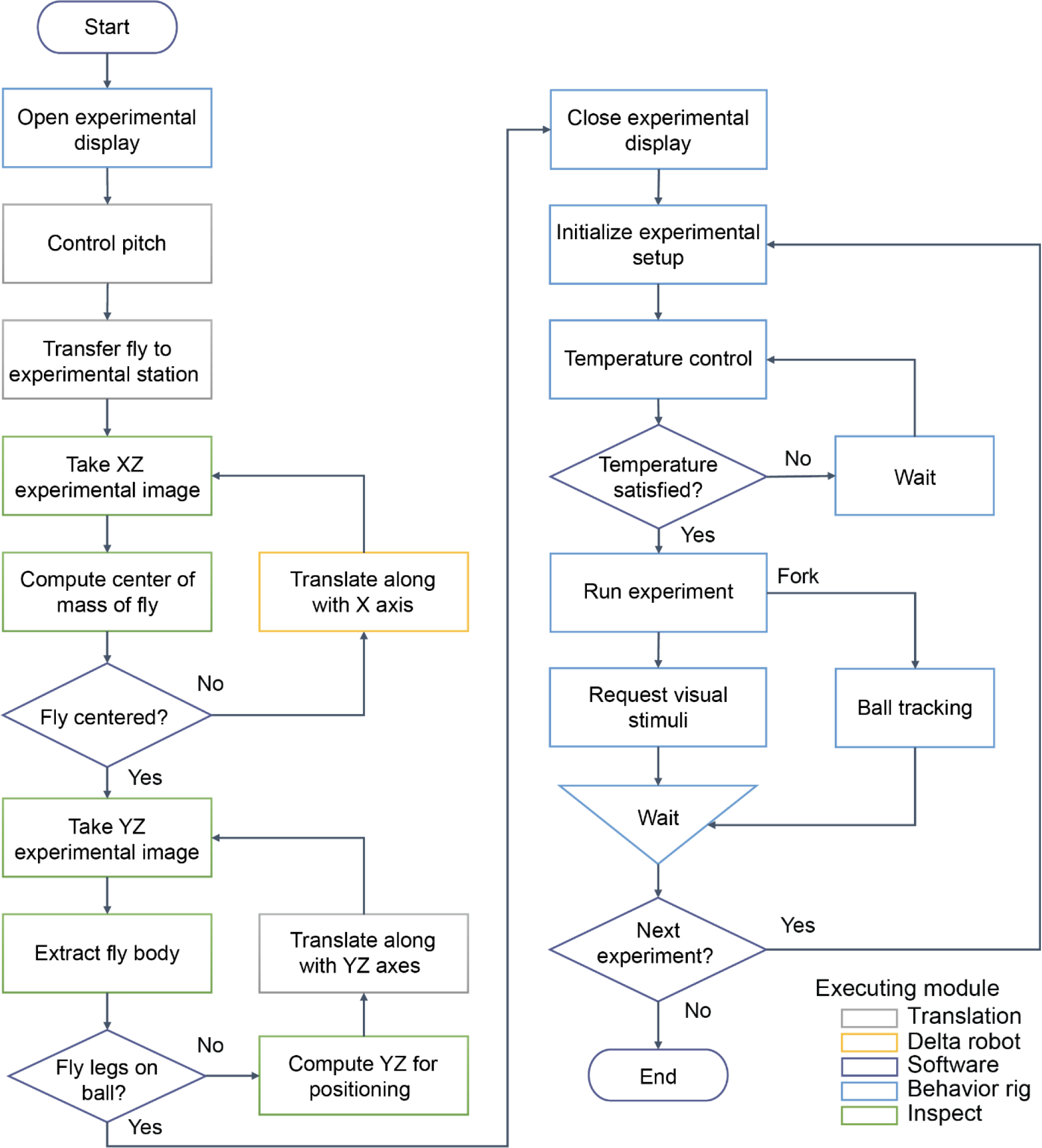
Experimentation workflows and algorithms. Automated experimentation workflow delineates the sequence of operations executed by the FlyMAX system during automated experimentation. We color-coded each step to indicate the specific component of FlyMAX responsible for its execution. Here’s a refined description of the process. *Initial setup*. The workflow commences with the adjustment of one of the experimental displays to facilitate the YZ view camera’s imaging of the ball tracking setup. This adjustment is crucial for enabling machine vision-guided control of the subsequent steps. *Fly Positioning*. FlyMAX adjusts the pitch of the fly carefully using the rotation station that is part of the gantry system. This adjustment ensures that the fly is in the optimal orientation for the upcoming steps. With the fly’s position updated, the gantry system moves, transporting the fly closer to the experimental station. This movement is crucial for positioning the fly in readiness for the central part of the experiment. *Fine-tuning fly placement*. The XZ view camera then takes over, monitoring the fly’s position to fine-tune the delta robot’s movements. This step precisely aligns the fly at the center above the ball, ensuring perfect positioning for the experiment. Utilizing the YZ view camera, the system estimates the fly’s exact position, allowing the gantry to make final adjustments. These adjustments bring the fly directly above the ball, compensating for any rotational shifts that might have occurred during the pitch control phase. *Experimental setup configuration*. Once the fly is securely on the ball, FlyMAX translates the experimental display back into place, effectively enclosing the behavior setup. This enclosure is essential for controlling the visual stimuli presented to the fly during the experiment. The system then proceeds to configure the experimental setup according to the predefined parameters, including temperature adjustments and other user-defined requirements. This configuration is pivotal for creating the desired experimental conditions. *Execution of experiments*. With the setup complete, visual stimulation and ball tracking are initiated concurrently. This simultaneous execution is a critical component of the experimental workflow, allowing for real-time observation and analysis of the fly’s responses. FlyMAX initiates a dedicated process for the continuous collection of data related to the ball’s locomotion. This data is essential for evaluating the fly’s behavior and the experiment’s overall outcomes. *Repetition of experiments*. The described steps are repeated multiple times to ensure the robustness and reliability of the experimental results. This repetition is crucial for validating the findings and drawing conclusive insights from the study.

**Fig. S12.**
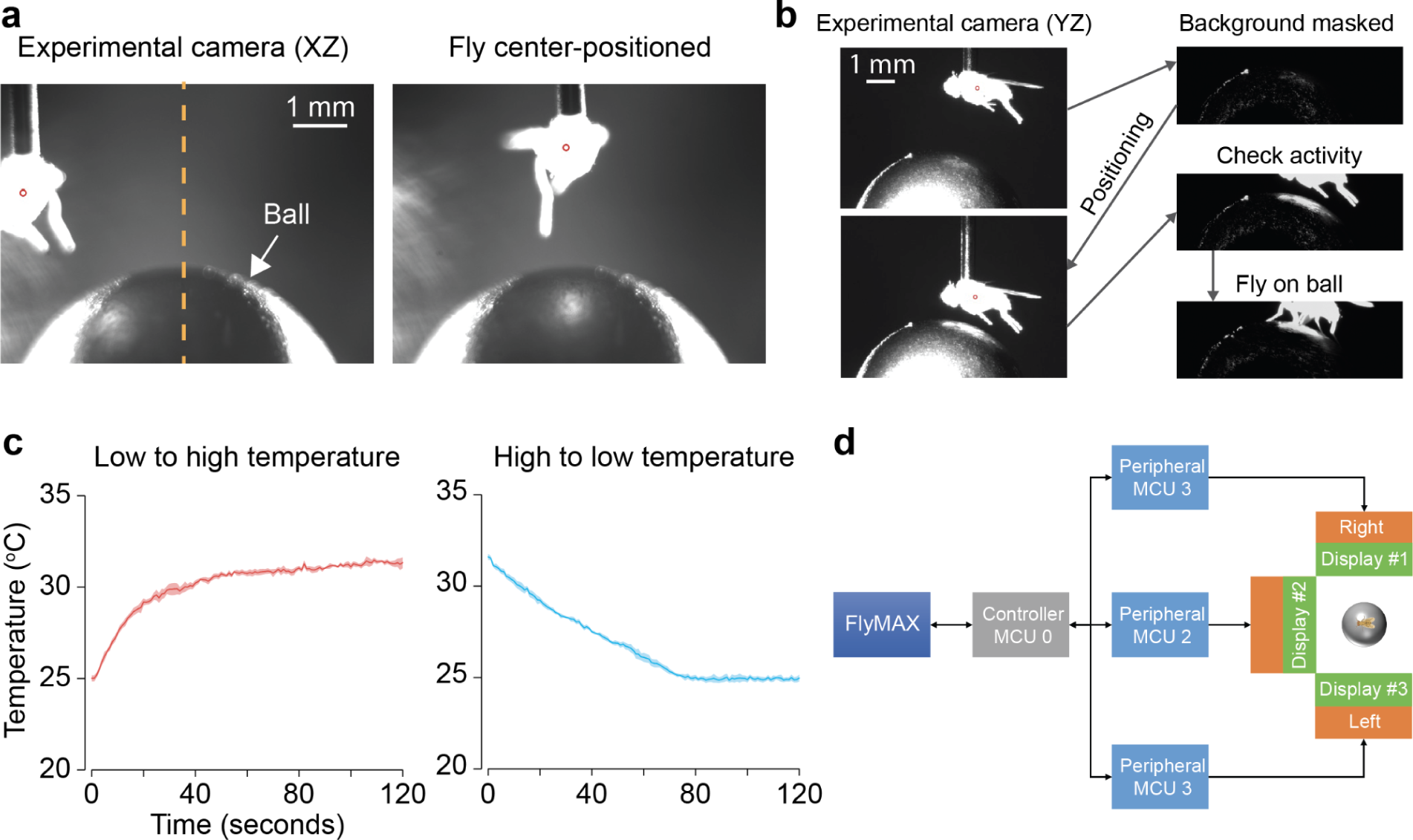
Automated manipulations in experimental stations, including positioning of tethered flies, thermogenetics, and visual stimulation. **a)** Machine vision based robotic control to position a fly. *Left*. XZ view camera captures the moment when the delta robot brings a captured fly close to the ball. For simple machine visioning to locate the fly’s position, we saturate the fly pixels to compute the center of mass of the fly. A small red open circle indicates the computed position of the fly. The orange dashed line indicates the center of the ball in the XZ view. *Right*. The fly has been moved to be positioned at the center of the ball. From the left image, the algorithm computes the distance between the red circle and the center of the ball, and the delta robot translates with the calibrated measurements to bring the fly to the center. **b)** FlyMAX uses YZ view camera to position the fly on the ball after XZ position occurred in (a). Using background masking, the algorithm extracts the fly’s body position. Similar to the method used for XZ positioning, it computes the center of mass of the fly and executes translation to position the fly close to the ball. The gantry performs Y translation to bring the fly’s body close to the ball. Then, the gantry also performs Z translation to make the fly stand on the ball. Once brought close to the ball, the robot checks if its legs are touching the ball. If the robot detects that the legs are touching the ball, it will slightly adjust the height to ensure the fly stands comfortably. **c)** Performance of thermal controlling. To test how quickly it can control temperature for thermogenetics, we measured the temperature over time after temperature control. *Left*. The temperature controller increased the temperature from 25 degrees Celsius to 32 degrees. *Right*. The controller decreased the temperature from 32 degrees Celsius to 25 degrees. For both heating and cooling, it took about a minute to reach the desired temperature. Error bars: s.e.m. across 4 trials for both low-to-high and high to low temperature control. **d)** Detailed architecture for generating synchronous visual stimulation. Communication for generating synchronous visual stimulation involves several components in FlyMAX. FlyMAX software communicates directly with the main controller via USB serial communication (MCU 0). The main controller communicates with three other peripherals via parallel communication through the direct I/O channels. Each peripheral has dedicated communication with a display panel via SPI communication. This setup ensures synchronized visual stimulation by coordinating communication between the software, main controller, peripherals, and display panels.

**Supplementary Table 1.**
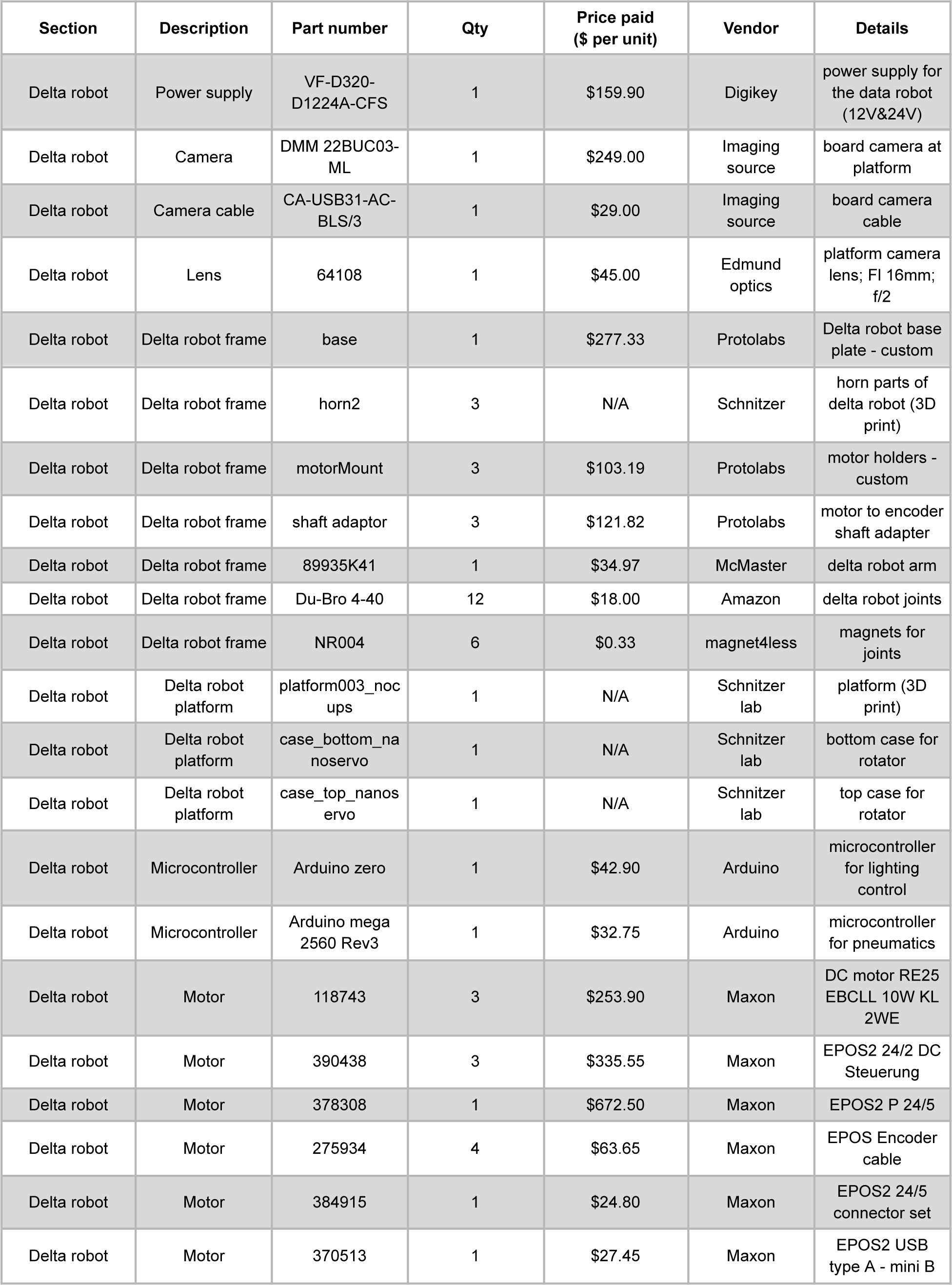
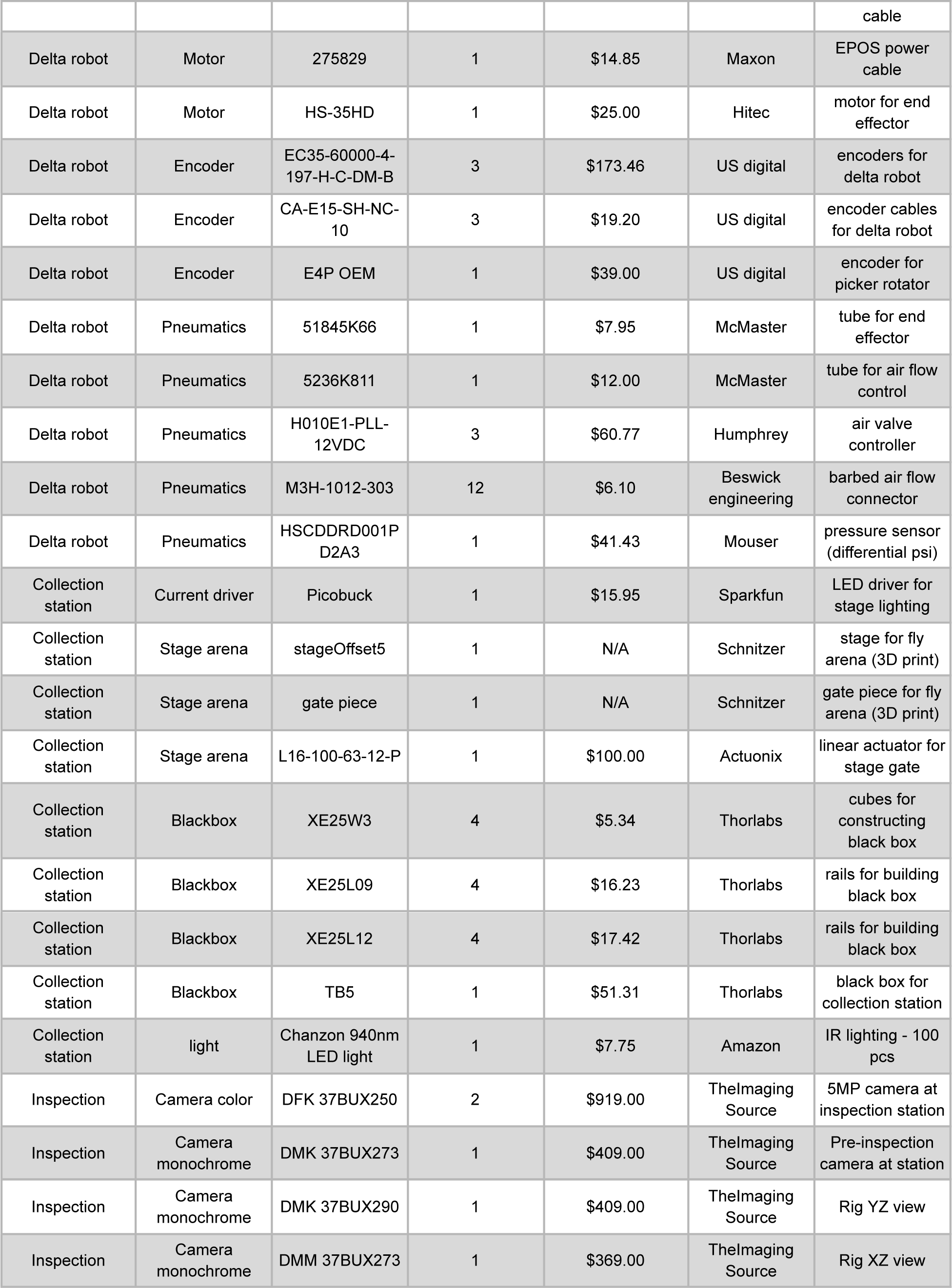
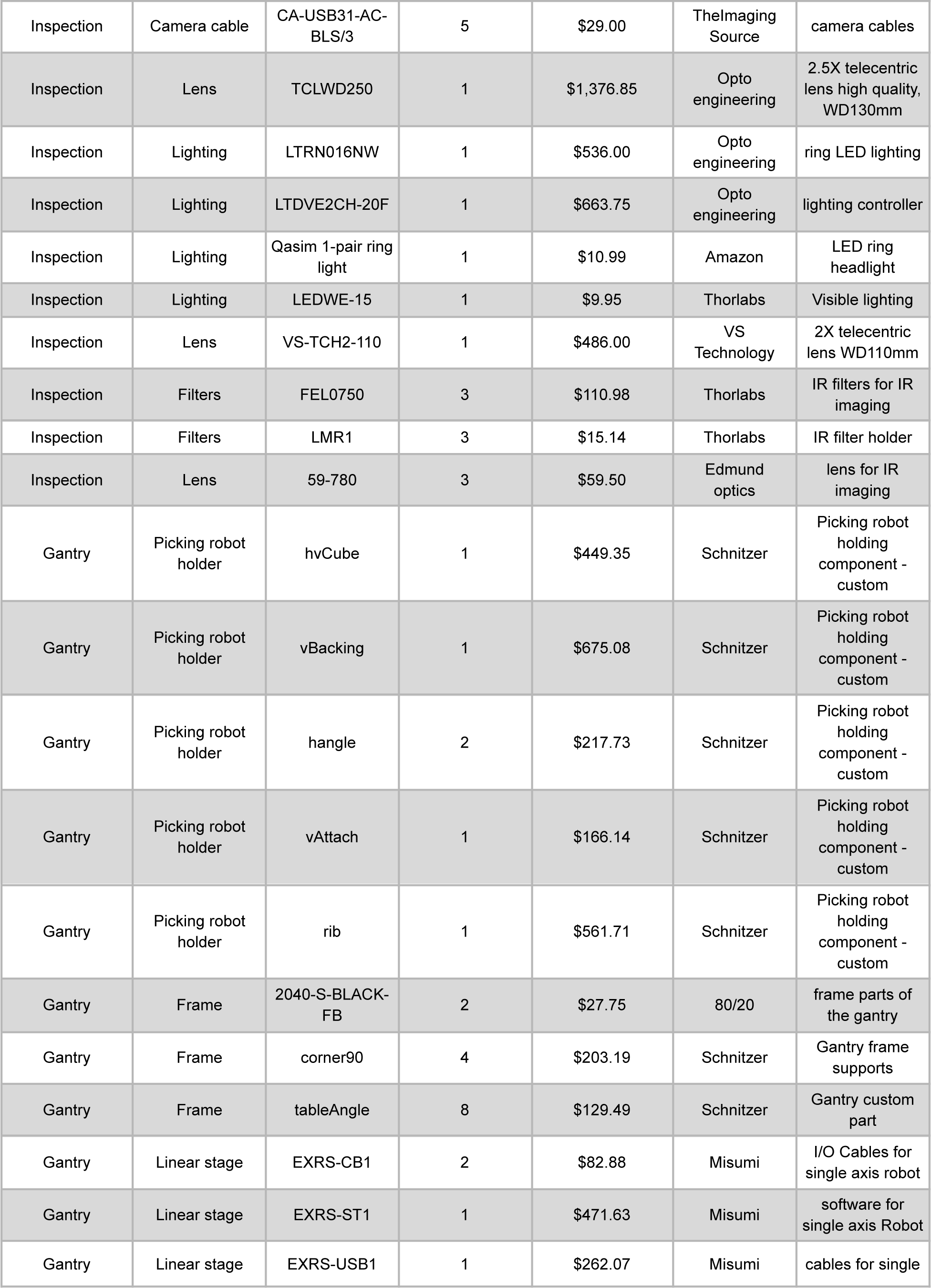
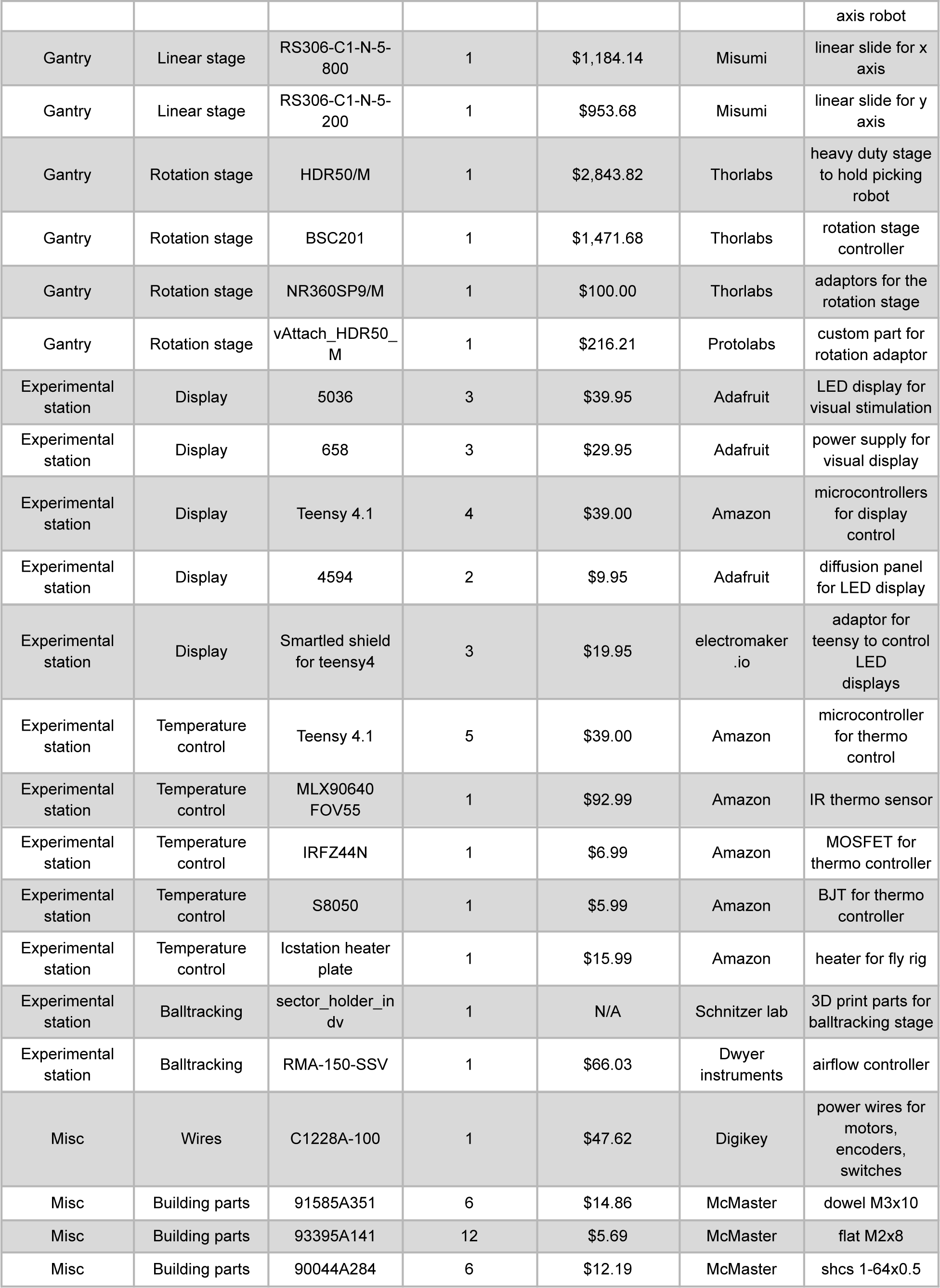
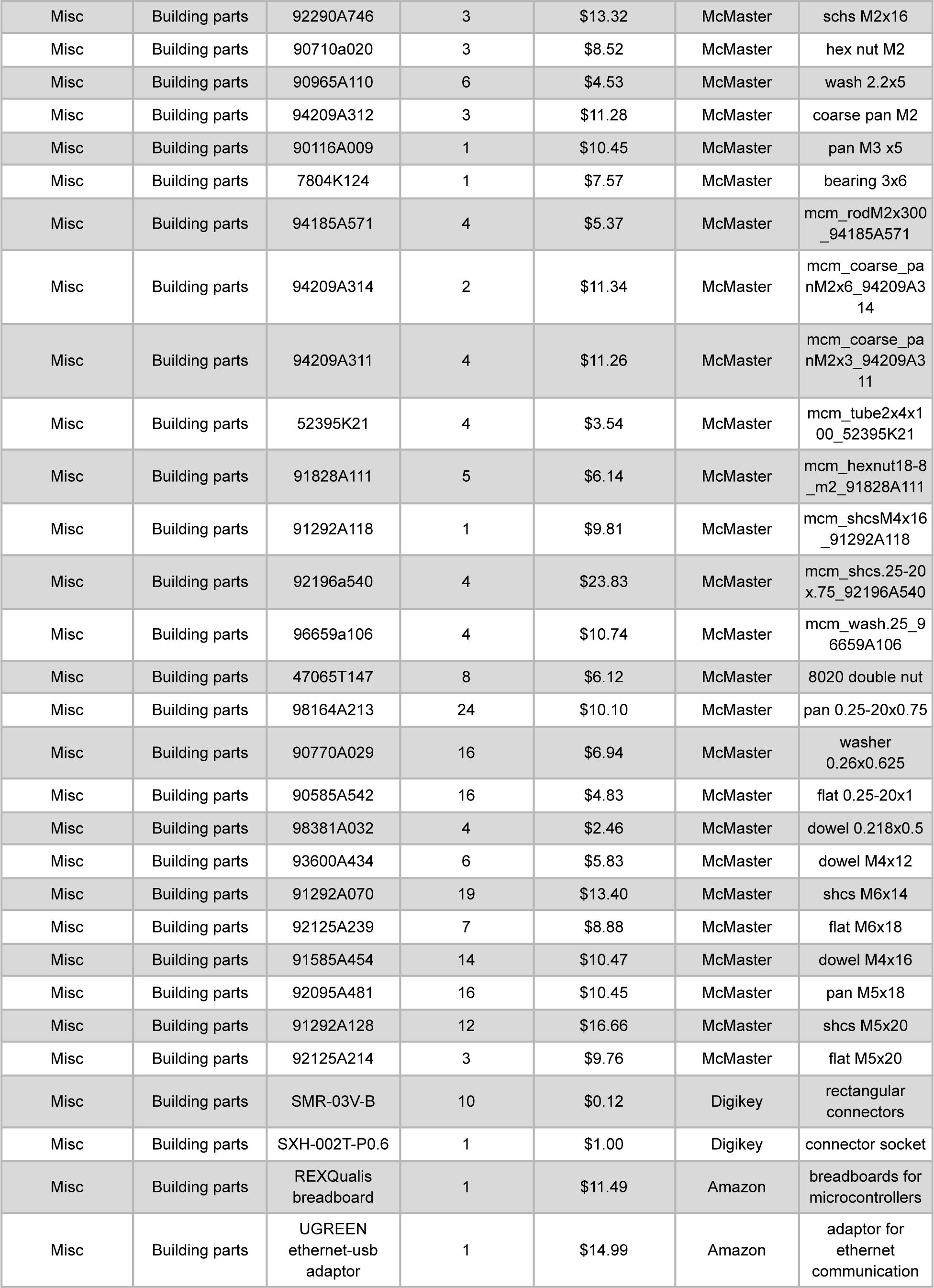
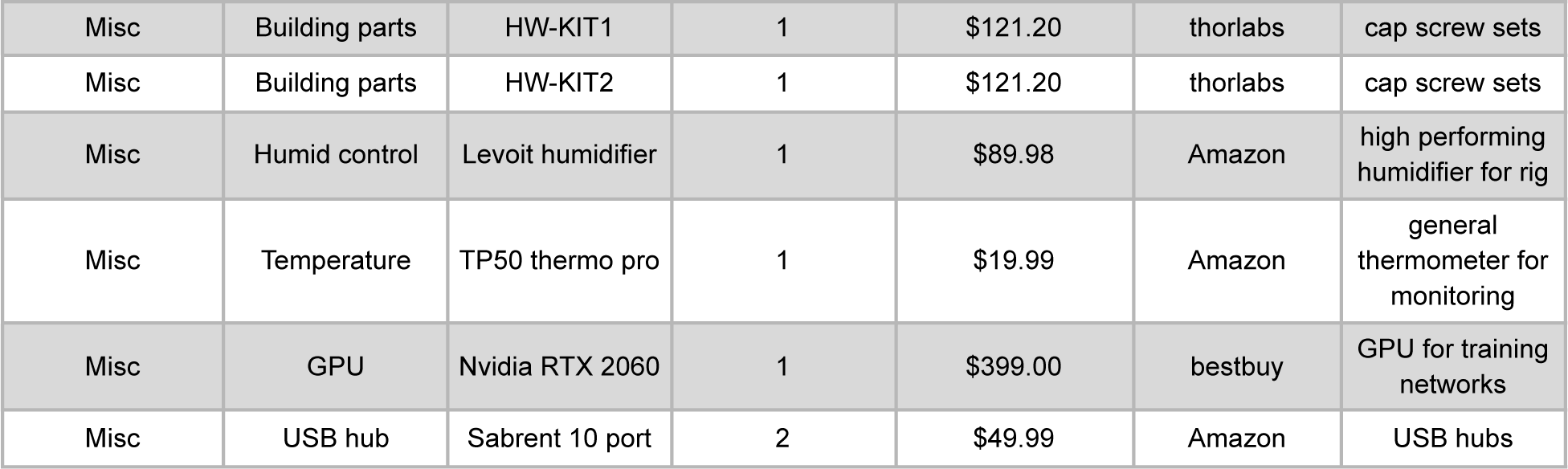
Bill of materials for the FlyMAX system. Bill of materials (BOM) for the FlyMAX system. Each row is a component that has information about the component’s section, description, part number, quantity, unit price, and vendor, along with specific details to ensure clarity and precision in replicating or understanding the setup.

**Supplementary Table 2.**
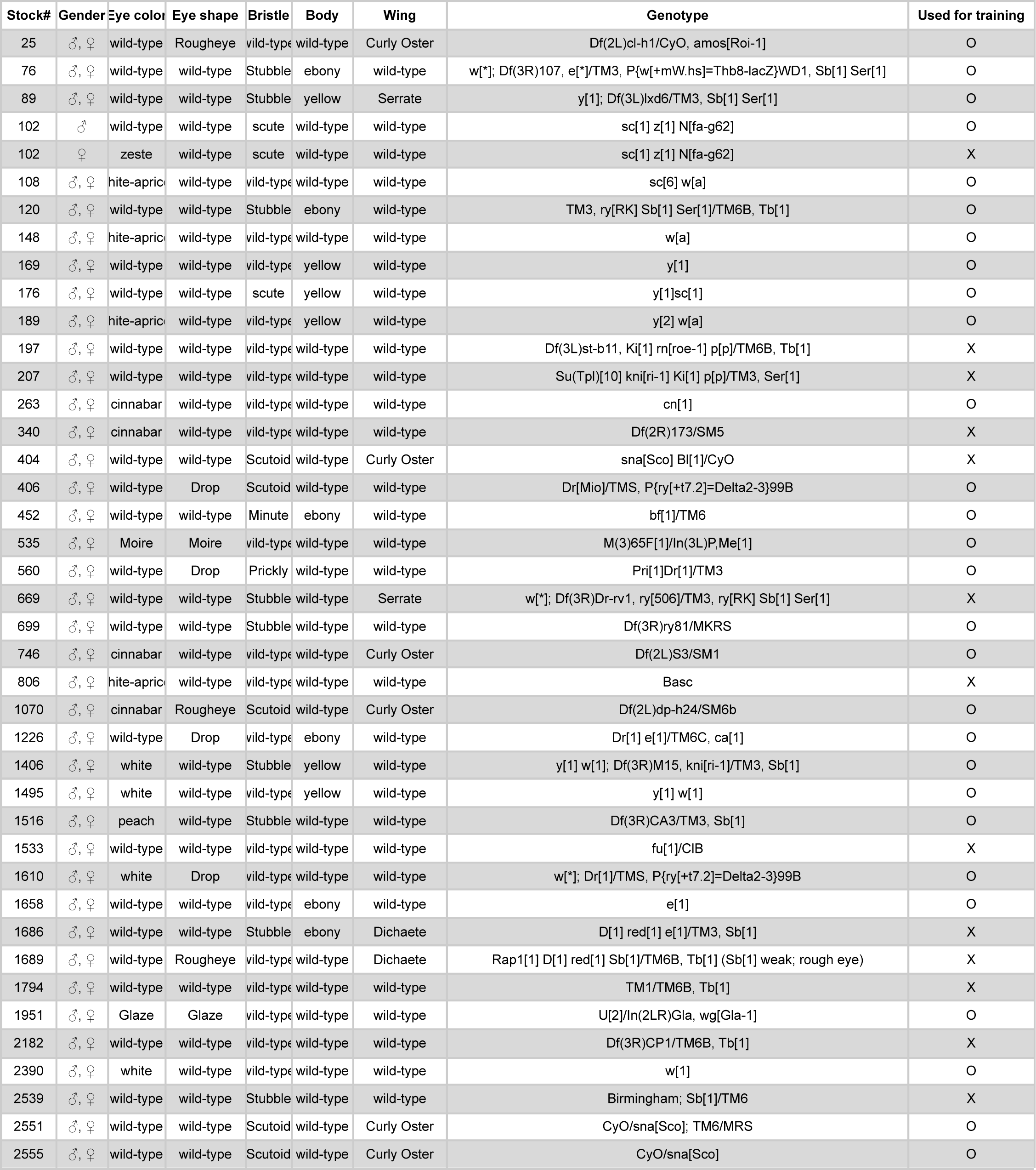
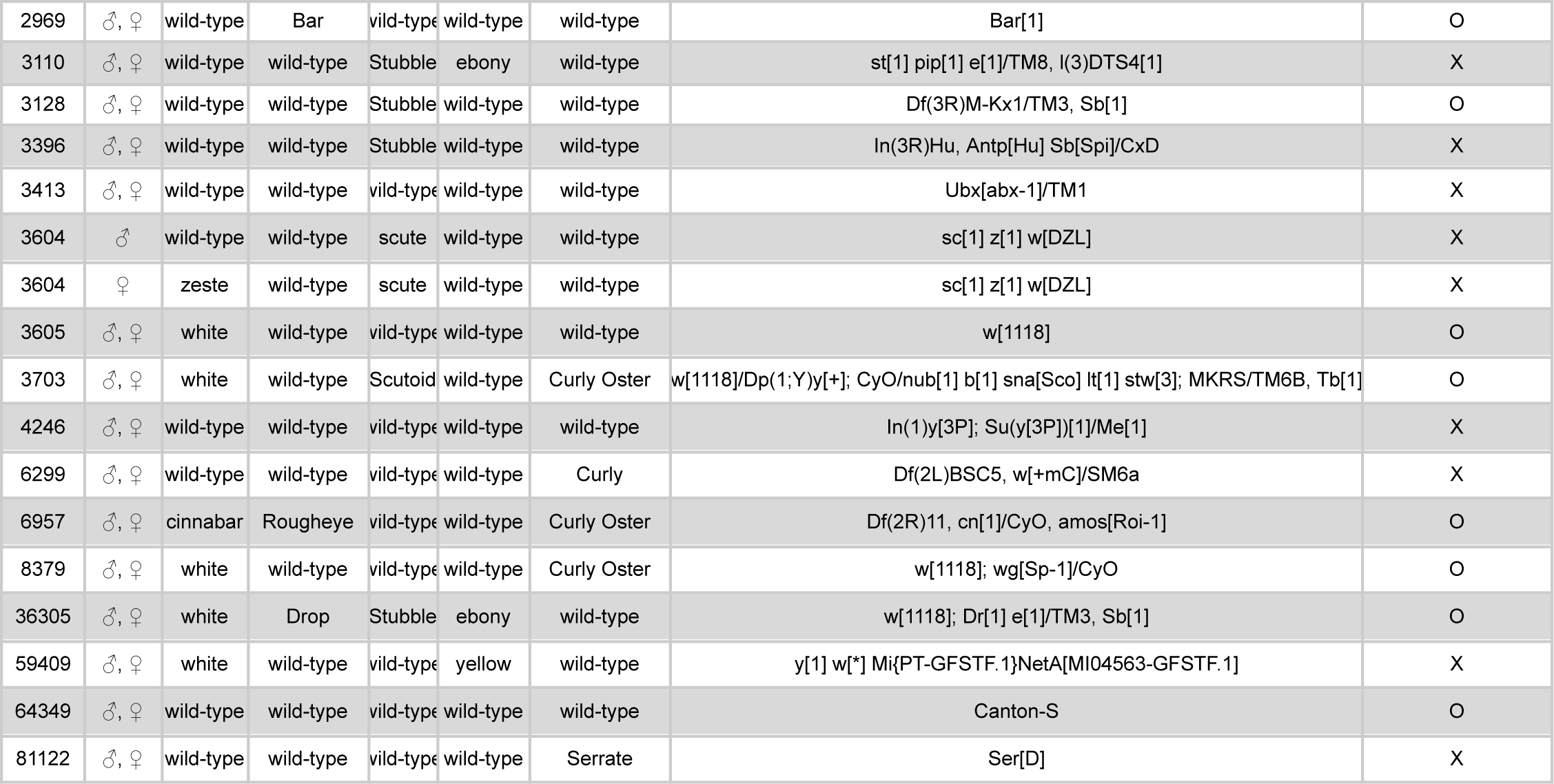
Fly lines used for phenotyping. The table lists the fly lines captured and image-processed by FlyMAX. The column ‘Used for training’ indicates those lines used for training neural networks.

**Supplementary Table 3.**
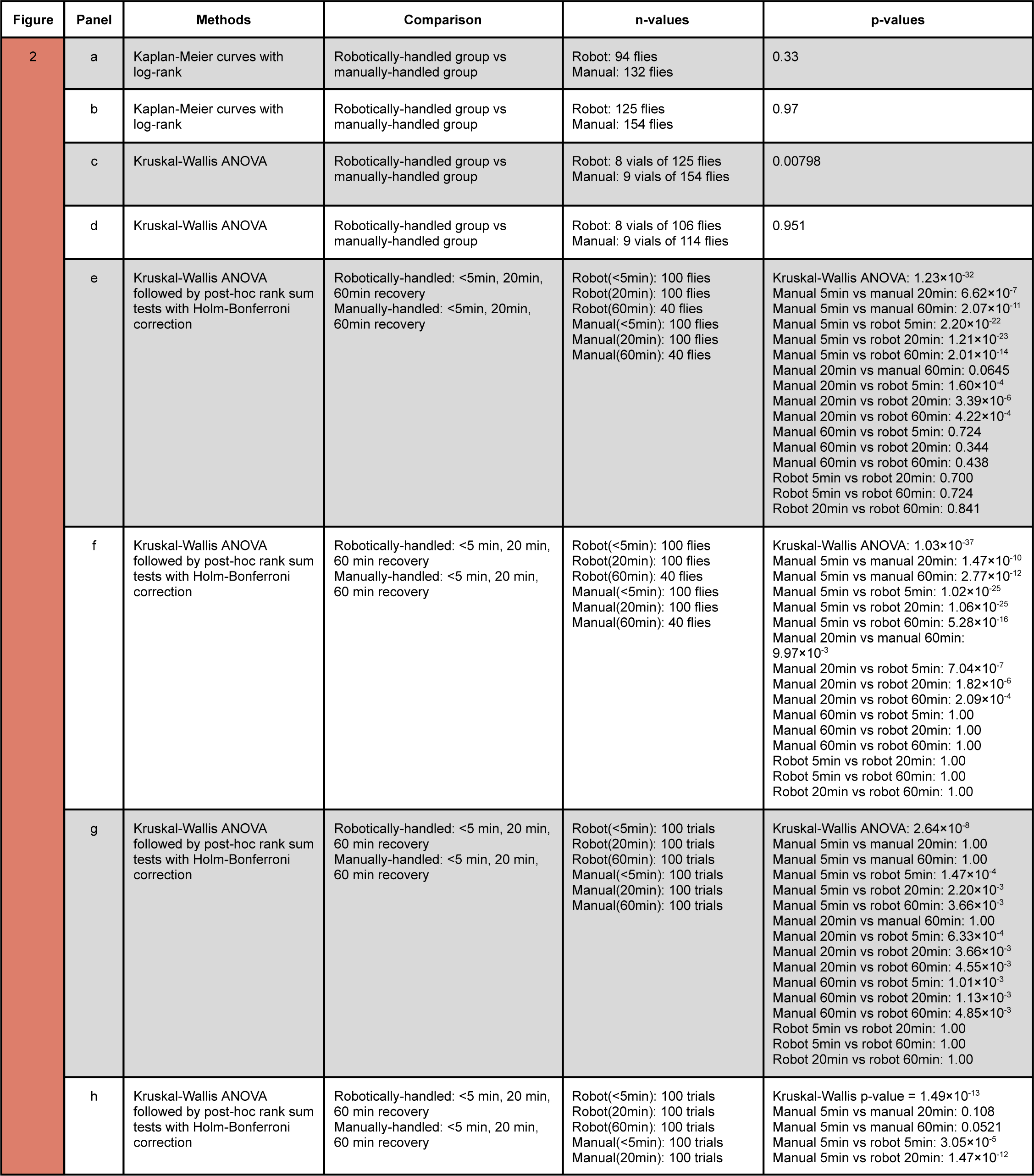
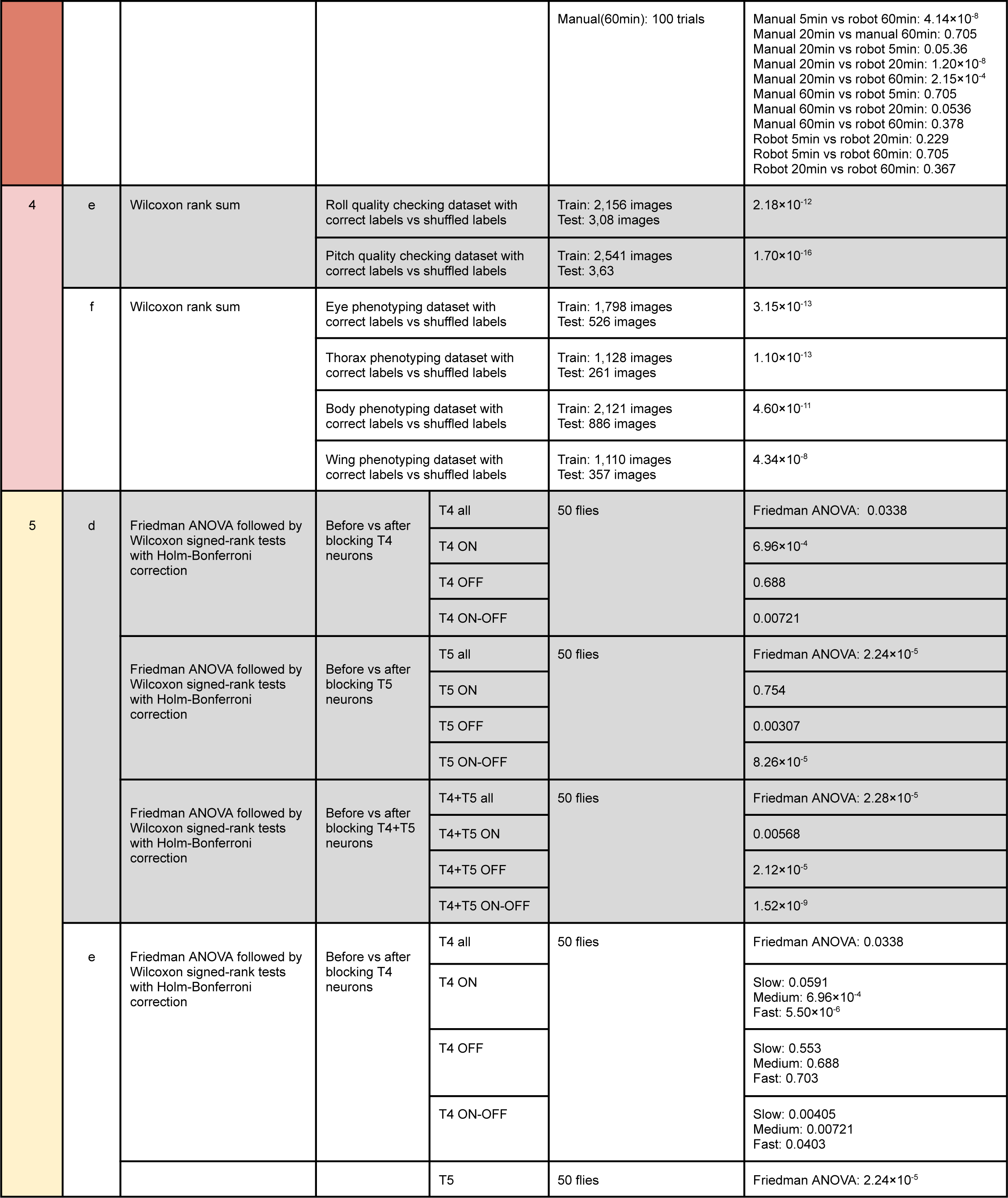
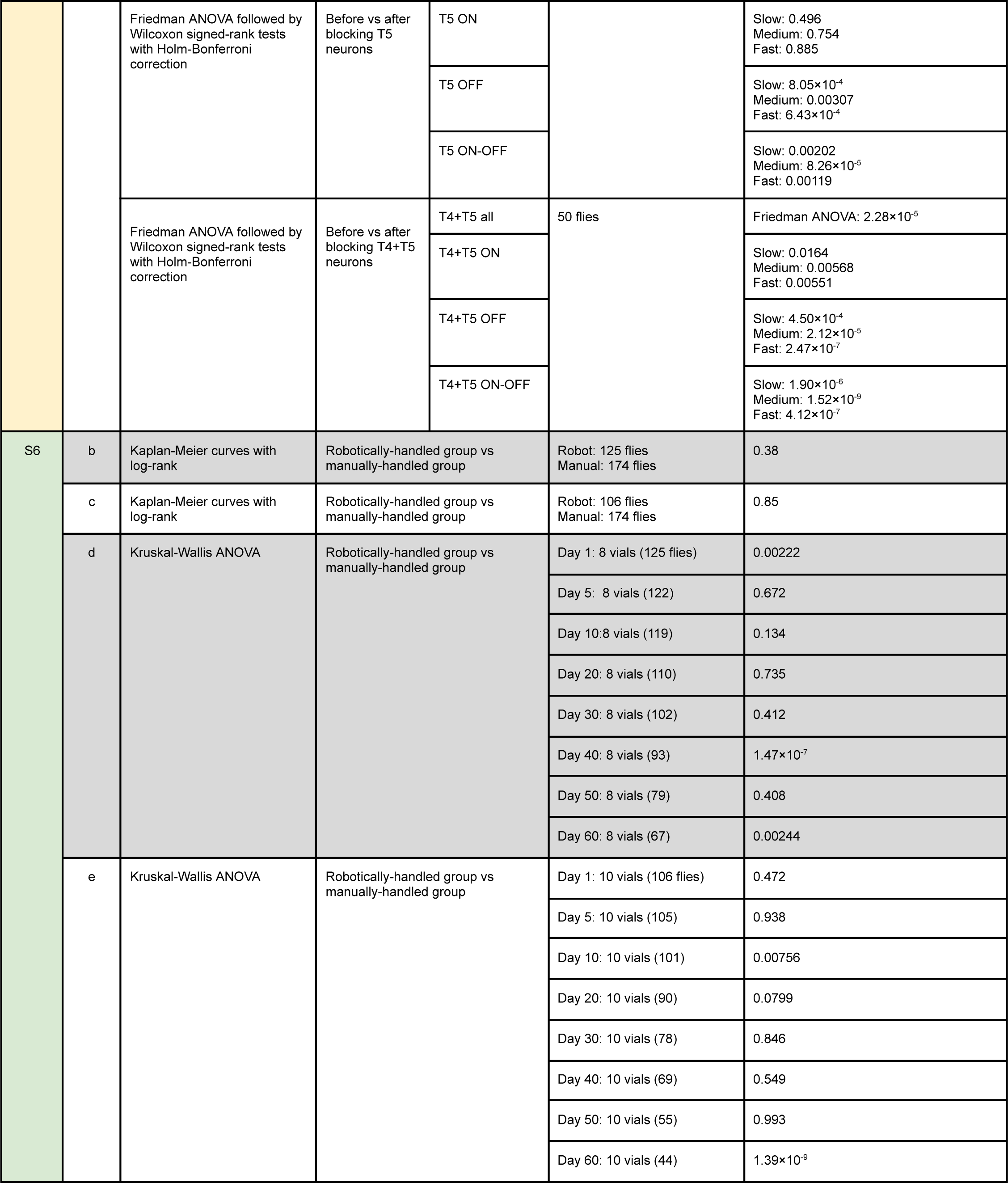
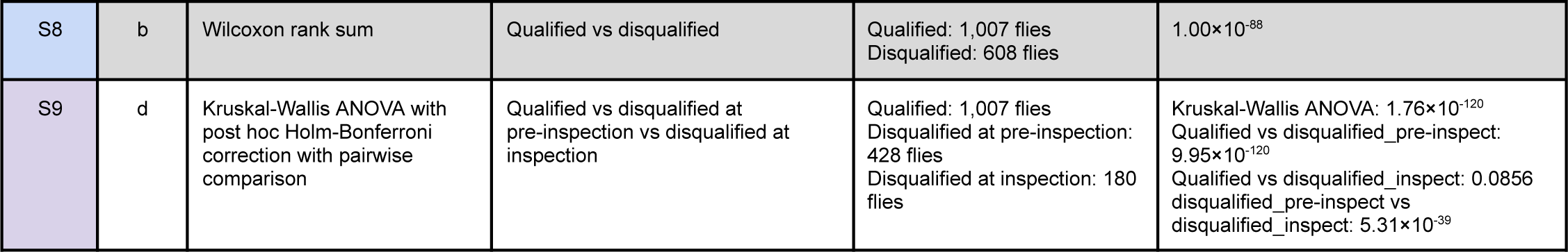
Statistical testing results. Statistical test results presented in the figures. The ‘Methods’ column specifies the statistical techniques used; the ‘Comparison’ column identifies the groups analyzed.

## Methods

### Fly stocks

Regarding health checking and high-throughput collection tasks, we sourced Canton-S (#64349) flies from the Bloomington Drosophila Stock Center (BDSC). We obtained additional fly lines from BDSC to check various phenotypes expressed in eyes, thoraxes, wings, and humerals (**Table S2)**. For *GAL4-UAS*, we used *R54A03-GAL4* (#50457), *R52H07-GAL4* (#50172), *R42F06-GAL4* (#41253), and *UAS-shi^ts1^* (#44222) from BDSC.

We raised flies on standard cornmeal agar media with a 12 hr light/dark cycle at 25℃ and 60% relative humidity. For behavioral experiments, we used flies 5–10 d old. For manual preparations of flies, we brought all flies to the cooled surface of an aluminum thermoelectric block for chilling (4ºC) and chose them in a random manner from a much larger group raised together for all studies; we did not use a formal randomization procedure for selecting flies. For automated preparations of flies, we gently inserted a vial with flies into the collection station, and FlyMAX captured them in the order of climbing up to the arena.

### FlyMAX software

We used C++ Visual Studio environment (Microsoft Visual Studio 2019 v142; ISO C++17 Standard) to develop FlyMAX software. We integrated various software development kits provided by vendors (camera control: IC Imaging Control v3.4.0.2550 from TheImagingSource; delta robot motor control: EPOS P v3.4 r1 from MaxonMotor; rotation motor control: Kinesis v1.14.23 from Thorlabs; light control: LTDV v1.0 from Opto Engineering) and communication protocols of C++ library to develop seamless communication integration for diverse robotic components within robotics stack, including camera systems, thermal sensors, motor encoders, and pressure sensors (**Fig. S4**). For computer vision, we used C++ OpenCV^66^ and its extra modules (v4.5.2) for computer vision libraries. We utilized other C++ libraries, such as Boost (v1.81.0; https://www.boost.org) and Eigen^67^ (v3.4.0), to facilitate file management, logging, matrix computation, etc.

The core software comprised the manipulation of the delta robotics, the collection station, and experimental stations, along with other dynamically linked libraries—camera, inspection, long-distance translations, and neural networks modules—that generate multiple instances at initialization (**Fig. S5**). Once the program initializes, it directs users to the interface, where they can run individual modules or autonomous tasks.

We categorized the user interface (UI) based on modular tasks to enhance user support. At the gantry UI, users can command the position of the delta robot, with options for predefined positions, such as the collection and inspection stations. The users can input global reference positions (XYZ positions of the end effector) or incremental distance to control fine positioning. Fly-picking UI has functionalities for XYZ translations with the rotation of the end effector. Users can also use this UI to perform piecewise tasks regarding picking a single fly: controlling the gate of the collection station, detecting, tracking and picking a fly. After the collection, users can further handle the fly for inspections, preparations, experiments, and so on. Camera UI provides access to individual cameras for image capture, simple processing, and configuration settings. Lighting UI allows users to control lighting setups, adjusting both the on/off states and intensity levels. Pneumatics UI manages bidirectional airflow and monitors pressure at the end effector. Experiment UI facilitates control over devices at the experimental station, managing visual displays, temperature settings, and experimental configurations.

FlyMAX supports full automation for tasks that do not involve human interventions. The collection task collects a well-tethered fly. After initialization, the software commands the picking robot to move to collection stations, pick a fly, translate to the inspection station, and inspect the picking quality (**Fig. S7a**). If standards are unmet, FlyMAX will go back to previous steps to iterate the process until it collects a qualified fly. Users can add following steps, for example, running experiments or sorting, to make the system handle multiple files with full automation. The system inspects the phenotypic details of a fly while the robot tethers it, integrating multiple steps to ensure inspection accuracy. (**Fig. S10**). The software is also capable of conducting complex experiments involving a tethered fly on a behavior rig, employing thermogenetics, and recording data seamlessly (**Fig. S11**). As FlyMAX software can collect data from all devices simultaneously, the data and log do not require further synchronization. These automatic features significantly reduce the laborious tasks traditionally performed by humans, enhancing efficiency and precision in experimental setups.

### Fly picking robotics development

We significantly modified and improved the earlier prototype of the fly-picking delta robot to facilitate robustness, accuracy, precision, and high-throughput picking performance for lab automation^27–32^. For the motor control, we reconfigured the dimensions of the delta robot to feature a 50mm diameter for the circular diameter of DC motors (Maxon 118743), a 20mm length for the robot arm’s horn, and 62 mm rods connecting the platform and the horns. This design alteration allowed the robot to translate extended distances along the Z axis relative to the XY axes.

We upgraded the robot frame from acrylic to aluminum for rigidity and durability. For each motor, we installed an optical encoder with 24,000 quadrature counts to guarantee an accuracy of 0.01mm. This enables the robot to reach various experimental stations accurately without physical constraints.

One main position controller (Maxon 378308) controlled the motors, while three peripherals (Maxon 390438) controlled individual motors through a CAN network. We installed limit switches for accurate homing. We customized the rods with a hypodermic tube (McMaster 89935K41) and inserted threaded balls (Du-Bro 884) as joints. We 3D-printed the frames of the horns and installed magnets (3D print: Veroblack; magnet: magnet4less NR004) on each end so that we could attach the rods with joints to the horns. We also installed the same magnets on the end of the platform, using them to make the robot tightly hold the platform and smoothly translate for fast motion.

For the platform, we also 3D-printed its frame similar to the previous design but removed the ring LED since we installed lighting in each station for specific purposes. For the end effector, we extended its length to 60mm to better position a tethered fly to a specific position for the experimental setup.

### Collection station setup

We designed a custom stage to facilitate the collection of flies, inducing them to come up so the picking robot can pick them up. Located in a dark room, the stage minimized visibility of the surroundings for the flies, reducing their activity levels. We simulated the picking model to determine the diameter of the stage arena that the platform camera could image and where the end effector could span (**Fig. S2**).

Furthermore, similar to prior studies aimed at limiting fly activity, we surrounded the stage with water to confine flies within the detection zone^31^. To give a good contrast for the thorax detection, we engineered a lighting angle of twenty degrees for infrared illumination (**Fig. S3a,c-d**). A linear actuator (Actuonix L16-100-63-12-P) with its dedicated controller (Linear Actuator Controller) handled the stage gate to ensure flies stayed in the vial when it did not perform the fly collection. The commands from FlyMAX software are close, half close, and open. The close command closes 90% of the space so that it does not let flies pass through and avoids risking injury to the flies by not closing too tightly. When a fly comes up to a stage, the gate initially half-closes to induce other flies to move closer to the gate in case the robot needs another fly. Additionally, during air breeze into the vial, the robot half-closes the gate to provide gentle air stimulations. We used Veroblack for 3D printing the collection station and a set of infrared LEDs (Chanzon 100F5T-IR-FS-940NM to construct custom ring lighting for the station (**Fig. 3Sb,e**). An LED current controller (Sparkfun COM-13705) regulated the lighting to provide constant illumination for accurate fly detection.

The gating and lighting controls inputted a signal from a microcontroller (Arduino Zero) that transferred a signal from the software to the related controllers. To initiate the collection process, the station opens the gate, turns on the stage light, and waits for flies to climb up to the stage. Once the camera at the delta robot detects a fly in the arena, the station closes the gate, and the picking process begins.

### Fly thorax detection and tracking for picking

We developed a fast, scale- and rotation-invariant detection algorithm to facilitate the rapid picking before a detected fly moves. We employed template matching to the thorax detection tasks, as the morphology of flies is common; however, challenges in detection lie in their variance in size and rotation from the camera view. A previous approach used template matching with rotated objects by rotating templates by 0 to 360 degrees with 10 degrees^68^, but this increased the computation time by up to 36 times and resulted in inaccurate template matching due to potential angle misalignment up to 10 degrees.

The robot uses top-down view images taken from the camera on its platform to locate the thorax position (**Fig. S1a**). To reduce the computation time for detection without sacrificing accuracy, we divided the algorithm into two steps: first, to detect the location of a fly body and then to accurately locate its thorax using template matching. To do this, we utilized adaptable thresholding to identify fly bodies and template matching to locate an accurate thorax position for picking. The algorithm downsamples the captured image for faster image processing (**Fig. S1b**). Then, it performs Otsu thresholding on the image to extract fly bodies^69^. Otsu’s method identifies a threshold value that binarizes the intensities of a given image. In monochrome images with intensity levels ranging from 0 to *L,* the pixel data is distributed in a normalized histogram where *n_i_* represents the number of pixels at level *i*, and the total number of pixels is *N*, such that:

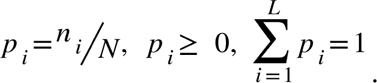

For binarization, we set two classes - presumably pixels related to background and flies - separated by a certain threshold, *k*, and defined two probabilities of class occurrence based on the threshold as

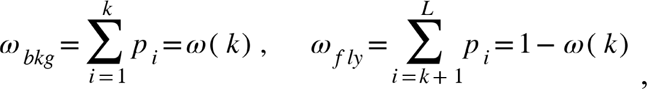

where *ω_bkg_* is a probability that a given pixel belongs to a background and *ω_fly_* is a probability that a given pixel belongs to a fly. To find the best k that separates two classes, we wanted to minimize the intra-class variations, σ^2^*_intra_*,

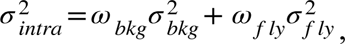

where σ^2^_bkg_ and σ^2^_fly_ are variations of each class. In other words, we wanted to maximize the inter-class variations, σ^2^*_inter_*, defined as:

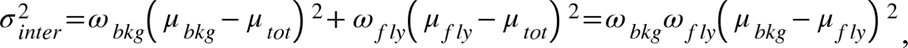

where *μ_bkg_*, *μ_fly_*, and *μ_tot_* are mean pixel values of background class, fly class, and total image, respectively, given

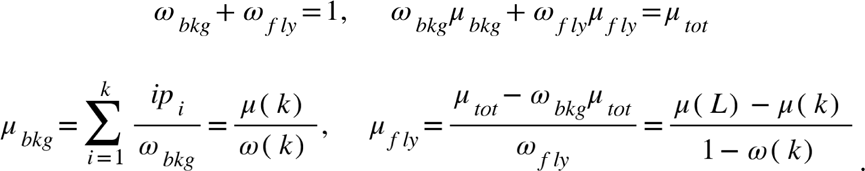

Therefore, for the optimal threshold value *k\** is

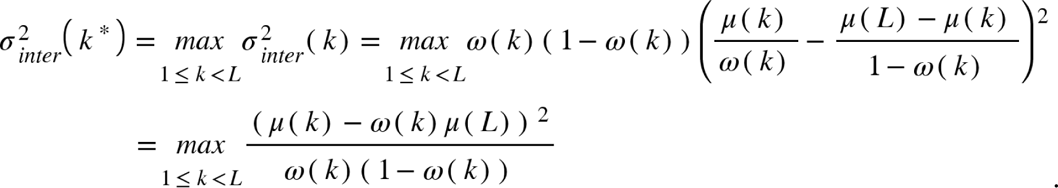

The binarization with *k\** value as a threshold reveals fly blobs (**Fig. S1c**). Of possible bodies, depending on the tracking algorithms—whether to track a body in the center or to look for the largest body size—the algorithm selects one body from possible bodies (**Fig. S1d**). From the determined fly blob, we needed the scale and orientation information for scale- and rotation-invariant template matching. As fly size varies according to age, gender, and genotype, the algorithm compares the body size of the template fly to that of the fly found in the image, scaling up or down to match the template size (**Fig. S1e**). For orientation information, the algorithm utilizes line fitting by minimizing the L2 distance between a line and the points of a detected blob, *r*, using the least squares method,

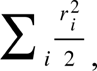

where *i* is each point of a detected blob. Once the algorithm fits the line, it computes its orientation.

For the template, we included a fly head and the thorax for them being relatively consistent compared to an abdomen and wings (**Fig. S1f**). The algorithm stores the position of the thorax relative to the (0, 0) coordinate of the template to estimate the thorax position after template matching. Based on the angular position of the blob, it rotates the image to match its orientation with that of the template (**Fig. S1g**). The algorithm uses another image with 180 degrees rotation to include the possible case when the fly head points downward. The algorithm computes the template matching by computing residuals spanning the image space, a normalized square difference, and finding the *x*, *y* coordinates with the minimum value:

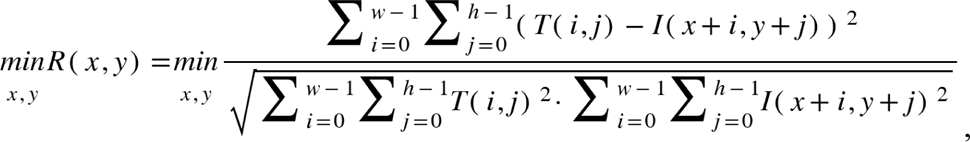

where *R*, *T*, *I* are a residual matrix, a template matrix whose width is *w* and length is *h*, and an image matrix, respectively. Once the algorithm estimates the thorax position, it transforms this position back to its original location in the robot’s space (**Fig. S1h**).

For validation of this algorithm, we compared the estimated orientations and thorax positions with manually annotated orientations and the positions (**Fig. S1i-j**). By comparing the residual values of template matching on random artifacts, we set 15 as a threshold value to determine if the best template matching position is a thorax (**Fig S1k**).

### Long distance translation development

FlyMAX offers versatile features to manipulate flies, and the long-distance translation module enables this by transporting the picking delta robot. We developed this module to work as a motorized gantry. We mounted the delta robot on a customized bridge-like frame with motorized translations of 800mm in Y axis (Misumi RS306-C1-N-5-800) and 200mm in Z axis (Misumi RS306-C1-N-5-200) with 0.01mm accuracy in both axes. A dedicated controller (Misumi EXRS-C1) controlled these motors, which were managed through direct I/O or dedicated software. To enable control by FlyMAX software, we designed a microcontroller to control 16 input channels and read 16 output channels for each controller (Arduino Mega2560). In addition, we mounted a rotation stage (Thorlabs HDR50/M) so that the picking robot can rotate to control the pitch level of the collected fly. We confirmed that the robot’s rotation remained stable for ±45 degrees during operation.

### Lifespan assay

To test the potential effects of robotic manipulations on flies, we compared the lifespans of a group of flies collected by FlyMAX with those of a control group collected manually (**Fig. 2a,b**). We sorted the flies on Days 0-1 after eclosion. For robotically handled flies, the robot first collected flies without the use of anesthesia, sorted them by gender, and then transferred them to food vials. We anesthetized manually handled flies with CO_2_ for gender sorting and then transferred them to food vials. In each vial, we placed 15-20 flies and maintained them at a temperature of 25°C and 60% relative humidity, following a 12-hour light/dark cycle. We transferred flies to fresh food vials every 2-3 days to ensure their sustenance and well-being throughout the study.

### Longitudinal climbing assay

To examine potential effects of robotic manipulations on climbing ability over time, we performed a time-lapse climbing assay, testing the flies’ climbing ability every 5 days (**Fig. 2c,d**). We reared flies at 25ºC and 60% relative humidity with a 12 h light/dark cycle. On each testing day, we put flies into a cylindrical tube, waited 60 s, tapped the tube 5 times to prompt the flies to climb from the bottom of the tube, and then counted the number of flies that climbed to a height of >175 mm on the side wall within 120 s^70^. Every 10 s, we recorded the number of flies that passed this 175 mm target line. We blocked the end of the tube with a vial cap, and after the assay, we gently put flies into a new vial of food. We performed all these steps without the use of anesthesia.

### Fly inspection setup

We installed a separate station for inspection to image flies under independent lighting conditions, allowing us to capture details of their morphology. Two cameras (TheImagingSource DFK37BUX250; IMX250) positioned at a 90-degree angle to each other, covering the XY space. Both cameras aimed at the position of a captured fly, enabling simultaneous viewing of two sides, such as the anterior and lateral aspects. Due to the trajectory of the delta robot and the long-distance translation capability, placing the lens closer to the subject would limit the translation of FlyMAX. To overcome this, we employed telecentric lenses with 2.5× (Opto Engineering TCLWD250) and 2× (VS Technology VS-TCH2-110) magnifications, with working distances of 132mm and 110mm, respectively. For the camera and lens set that inspected mainly the lateral aspects of a fly, we chose a 2.5× lens with an industrial-level LED lighting (Opto Engineering LTRN016NW) to capture details of a live fly while moving and flying. The other set of camera and 2× lens used a custom ring light (Qasim QAAE0034-100MM-W) for anterior view. A fast LED controller (Opto engineering LTDVE2CH-20F) controlled two lighting setups. The black box covering the picking robot held a background for high-contrast fly imaging (Gaffer power GP-CHROMEGREEN2x30).

### Picked quality checking

To ensure the reliability of our autonomous system in the fly collection, FlyMAX incorporated a systematic approach for assessing the accuracy of the picks, specifically evaluating whether it picked the thorax part correctly. For this, FlyMAX checked the pitch and roll of the collected fly. To ensure high throughput in checking the quality, we installed a pre-inspection at the collection station with infrared light to filter out apparent bad picks on a wing, abdomen, head, etc. (**Fig. S9**). Once a collected fly passed pre-inspection, it underwent another inspection at the inspection station for detailed checking. At the inspection station, the robot checked roll and pitch qualities with anterior and lateral views, respectively.

For the models that checked picking quality, we used LibTorch (v1.8.1+cu111), a C++ version of PyTorch, to train computational classifiers that differentiate good or bad picks based on given images taken during fly inspection. We built two classifiers for roll and pitch checking separately. To construct datasets, we ran automatic picking and inspection tasks of over a thousand flies, from which we collected 2,464 and 2,904 for roll and pitch classifiers, respectively. For the datasets, we performed 8-fold cross validation, such that the number of training and testing sets for the roll classifier are 2,156 and 308 and that the number of testing and training sets for the pitch classifier are 2,541 and 363, respectively.

For neural network architectures, we used EfficientNet (v2b0) pre-trained with ImageNet^39^ for computer vision tasks. We modified the last linear layer of EfficientNet to fit the number of labels we used for each classification task: two for the roll classifier (bad and good) and four for the pitch classifier (bad, low, middle, and high). We then added logsoftmax function as the final layer for classification, such that the output of *i^th^* class is:

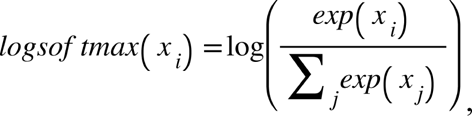

where *i* indicates ‘bad’ and ‘good’ for roll classification and ‘bad’, ‘low’, ‘middle’, and ‘high’ for classification. With these outputs, we optimized the weights of classifiers with negative log likelihood loss:

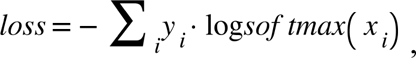

where *y_i_* is the value for an input image. We then used the Adam optimizer to update the model parameters:

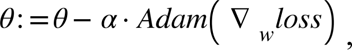

where *α* = 1 ·10^-4^ is a hyperparameter that specifies the learning rate, which we optimized using the validation dataset, and Adam(∇*_w_loss*) denotes Adam optimizer, an extension of stochastic gradient descent that provides faster convergence through adjustments of the learning rate during training. (We used the Adam optimizer’s standard parameter values for adjusting the learning rate^71^). We trained the networks by updating their matrix elements across 100 full passes through the entire training sets. We used Python and C++ for training and testing, C++ language for deploying to FlyMAX. For training and testing, we took advantage of a GPU (NVIDIA RTX2060).

### Phenotyping method

We utilized deep neural networks for classifying the phenotypes. For phenotyping body color and wings, we used EfficientNet (v2b0) pre-trained with ImageNet^39^. The phenotyping classification on eyes and thorax end consists of two steps: first, identifying the positions of interest, and then classifying the cropped image around these positions. We used a model trained from DeepLabCut (DLC) for pose estimation^38^. In the DLC, we manually labeled positions of eyes and thorax end on a hundred flies whose phenotypic features vary (list the genotype of flies in the labeling set). We included flies that flew or were out of focus to increase the robustness of the pose estimation. We used the interface provided by DLC to label, train, and validate the model accuracy. We trained the models using the Python TensorFlow frameworks and converted them to ONNX (https://onnx.ai) models, which can be loaded from the TensorFlow C++ framework. We implemented the forward layers of DLC in C++ to compute the pose estimation from the feature values and verified by comparing them with the output from Python. With the estimated position, we cropped a 500 × 500 image centered on the estimated pose, which served as the input for the second classifiers. For these classifiers, we used the same EfficientNet architecture pre-trained with ImageNet. Similar to the quality-checking classifiers, we modified the last layer of the EfficientNet to output feature values of the labels, added a final layer with logsoftmax, and used Adam as an optimizer for training with the same learning parameters. For eye phenotype, we used 9 labels, and the number of training and testing sets was 1,798 and 526, respectively (**Fig. 4d**; left). For thorax end phenotype, we used 6 labels, and the number of training and testing sets was 1,118 and 261, respectively (**Fig. 4d**; middle). For the body color, we used 3 labels, and the number of training and testing sets was 2,121 and 866, respectively (**Fig. 4d**; right). For wings, we used wild type and Curly phenotype as labels, and the number of training and testing sets was 1,110 and 357, respectively.

### Fly experimentation setup

To conduct an advanced behavioral assay with thermogenetic control, we set up a visual sensory behavior rig for an experimental stage comprising visual displays, a temperature controller, a ball tracking system, and positioning cameras.

For the visual display, we used three LED display panels (Adafruit 5036; 32 × 64 pixels per panel) that are high-performing and cost-effective. We positioned the three 80 mm-long panels (32 pixels per panel) to form 270-degree azimuth and 145-degree elevation, placing them 45 mm away from the fly’s position to benchmark prior works^34,43^. We covered the LED displays with diffusion acrylic panels to reduce glare and rescale the intensity range from 0 to 30 mW/m^2^. We made the pixel size of each LED less than 3 degrees, ensuring it was smaller than the interommatidial distance of the *Drosophila* eye. In order to position flies to the ball accurately, we installed cameras for XZ and YZ plane aspects. Since the displays surround the ball tracking setup, we used a linear stage to move one of the panels, allowing vision for the YZ plane camera.

For thermogenetic perturbation, we utilized a thermistor (HT10K; Thorlabs) to regulate the environment’s temperature. We placed an infrared remote thermosensor (MLX90640; 32x24 pixels; 55-degree field of view) 50 mm away from the fly to monitor the body temperature during experiments.

### Stimuli display

The visual stimuli consisted of three types: ON, where a light edge moved, OFF, where a dark edge moved, and ON-OFF, where a grating pattern moved. Each stimulus type could move in two directions: left to right and right to left. Each stimulus had three options for temporal frequencies: 0.4 Hz, 1.3 Hz, and 4 Hz. The refresh rate of the display was 300fps, and we used green for the color of all stimuli.

To control each panel, we utilized a microcontroller (Teensy 4.1; PJRC) with Serial Peripheral Interface (SPI) communications. As we needed three panels, we had three microcontrollers and one additional microcontroller to command the others (**Extended Data Fig. 10d**). To generate light patterns on the displays, we programmed the main controller (MCU0) to monitor and command the peripherals (MCU1-3) via direct I/O. For instance, when generating an ON stimulus from left to right, MCU0 sends a signal to MCU1 to execute the ON stimulus and waits for confirmation of execution. Then, MCU0 sends the signal to MCU2, and then MCU3. Each communication channel with MCU0 is separate, allowing MCU0 to send simultaneous signals to all peripherals for the ON-OFF grating pattern execution.

For automatic protocol, we adopted prior work on generating visual stimuli^34^. During experiments, the program randomly chose a type, speed, and direction for each trial. We set 33.3% for the contrast (Michelson contrast: (I_max_-I_min_)/(I_max_+I_min_)) for all types of stimuli. The stimuli appeared on four windows, each with an angle of 67.5 degrees. With this window angle, the speed of the moving edge corresponds to 30 deg/s, 90 deg/s, and 270 deg/s for the frequencies of 0.4 Hz, 1.3 Hz, and 4 Hz, respectively.

### Thermogenetic control

Leveraging the data read from the remote thermosensor, FlyMAX utilized thermal values extracted from a designated bounding box encompassing the fly’s position to compute the average temperature of the fly’s body. We verified the temperature measured by the remote sensor using a tactile thermosensor. We developed a microcontroller to function as a Proportional-Integral-Derivative (PID) controller for building a temperature controller. With feedback from the remote thermosensor, we attained the desired temperatures in around one minute (**Extended Data Fig. 10c**). The PID parameters we used were the following:

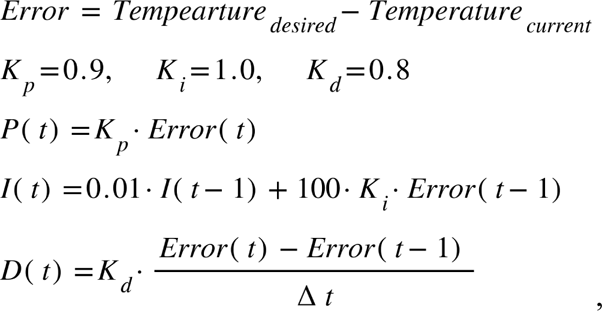

where K_p_, K_i_, and K_d_ are proportional, integral, and derivative gains, respectively. We modified the integral terms to prevent windup issues, such as error accumulation over time, by limiting the accumulation to a specific time frame.

### Measurements of fly locomotion

To track the locomotion of individual flies walking on a trackball (**Fig. 2e-h**; **Fig. 5d-g; Fig. S12a,b**), we used a setup similar to that of prior studies^49–51^. Specifically, two optical USB pen mice (i-pen mouse; Finger System) were aimed at the equator of an air-suspended, hollow high-density polyethylene ball (6.35-mm diameter; ∼80-mg mass; Precision Plastic Ball Co.). The pen mice were positioned 2.3 cm away from the ball and tracked its rotational motion (120 Hz readout). We configured FlyMAX software to collect the readouts from the pen mice during experiments. We computed the fly’s pitch, roll, and yaw velocities using code written in MATLAB^72^ (v2022b; Mathworks).

### Visual sensory motor behavior assay

To check the visual sensory motor behavior of flies, we set up a ball tracking system to record locomotive measures given visual stimuli described in the stimuli display section. To assess the activity of robotically versus manually tethered flies (**Fig. 2e-h**), FlyMAX ran randomized trials but only used locomotive measures of ON-OFF stimuli with 1.3 Hz frequency to compute activity levels. Each fly underwent three trials of the same stimuli, and we averaged the activities. To manually prepare flies for the control group, we initially put a fly on a cold metal plate to immobilize it and then used a thin tungsten rod (A-M Systems #718500) to tether its thorax with UV-cured glue (Kemxert E036). After tethering, we waited for the specified recovery time for each group. We immediately put the group with less than 5 minutes of recovery time into a behavioral experimental rig and began the experiments. We put the groups with recovery times of 20 minutes and 60 minutes into the behavioral rig after the designated waiting times, respectively, before beginning the experiments. For the fly behavior experiments (**Fig. 5**), we randomized the order of stimulus sequences, conducting five trials for each type, speed, and direction of the stimuli, and averaged the results. We used the same flies before and after blocking neurotransmission, varying only the ambient temperature varying between conditions. For the post-processing of the readout data, we used MATLAB after experiments.

We used the measured velocities to compute a turning index^5^ (TI), defined as the unitless ratio of yaw velocity to pitch and roll velocities of the ball:

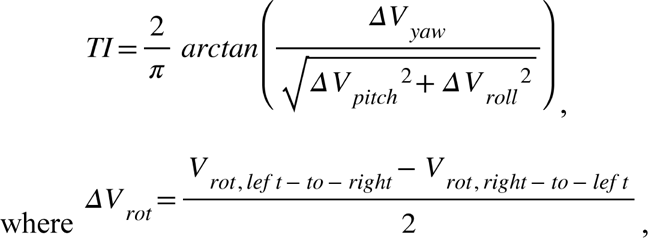

*rot* can represent any type of rotation, including yaw, pitch and roll, and *left-to-right* and *right-to-left* indicate the direction of the stimuli (not the direction of the ball). During all experiments, flies responded to visual stimuli that moved either from left to right or right to left. For each type of rotation, we computed *ΔV_rot_* by changing the sign of *V_rot,right-to-left_*, adding it to *V_rot,left-to-right_*, and dividing the sum by 2. This approach allowed us to handle the stimuli moving in both directions. For the turning index, only the direction of yaw velocity is important, as it indicates the direction of turning in response to the stimulus. For yaw, we set the ball’s counter clockwise direction as positive, and thus *V_yaw,left-to-right_*is positive when a fly follows the direction of a left-to-right stimulus and thus turns the ball right to left, or counterclockwise. Therefore, the turning index reflects both the direction of the response and the relative magnitude of yaw velocity compared to other rotational velocities.

### Autonomous high-throughput fly collection

To check the performance of fly collection, we performed about a thousand fly picking experiments (**Fig. 3**). The process follows the workflow described for fly collection. We included some other functions to make FlyMAX run continuously without interruption. During the detection process, if a fly did not appear for 30 s, the delta robot stimulated the flies inside the vial at the collection station by gently pushing the airflow through the passage. The gate was half-closed to prevent overstimulation that could cause the flies to avoid climbing up. The air pressure we used was gentle, and depending on the fly’s height, picking at the same position might not successfully capture the fly. To overcome this, FlyMAX adjusted the picking height by estimating the target fly’s size through trial and error. We put hundreds of flies in a vial to encourage climbing to the collection station, and whenever the number of flies in the vial became low, we rapidly swapped it with another batch. This process was swift and does not require stopping FlyMAX on the fly. We also implemented systems to handle unexpected errors, such as camera failures, by detecting anomalies in real-time and responding appropriately without manual intervention. There were some occasions when the end effector was clogged due to sticky food, or a fly got stuck that mere air pressure could not clean it. To resolve this issue, we installed a water fountain and programmed the picking robot to clean itself by sucking up the water and flushing it. With all automation systems in place, FlyMAX was able to run continuously for more than 15 hours.

### Real-time phenotyping

After the phenotyping models were validated, we commanded FlyMAX to check fly phenotypes in real-time. We mixed vials containing wild-type flies (#64349) with flies with white eyes, curled wings, and scutoid thorax ends (#3703). We did not control the proportions of different fly types for this task. Once FlyMAX picked up a fly and used pre-inspection to filter out badly picked flies, it proceeded with autonomous phenotyping (**Fig. S10**). We used a hundred identified flies for each group to run real-time classification for the eye, wing and thorax end phenotypes. We utilized classifiers trained on the data shown in **Fig. 4f** to perform binary classification between two fly groups. After the phenotyping, FlyMAX took multiple images of the fly from different angles for comparison and post hoc analysis. For each phenotype, we considered the first hundreds of trials for the validation process.

### Statistical analyses

We performed all statistical analyses using MATLAB (v2022b; Mathworks) software. FlyMAX writes all the loggings and data in .json format using the Boost logging feature. The MATLAB code can read and process these JSON files. We chose sample sizes using our own and published empirical measurements to estimate effect magnitudes.

For statistical testing, we employed various methods tailored to the nature of our data and the specific hypotheses we aimed to test. For the survival assay, we used the log-rank test, a non-parametric test to compare survival distributions between robotically and manually handled groups. We performed non-parametric Kruskal-Wallis and Friedman ANOVAs to avoid making assumptions about normal distributions or equal variances across groups. To perform post-hoc pairwise comparisons, we used the Wilcoxon rank sum test and Wilcoxon signed-rank test with Holm-Bonferroni correction for multiple comparisons. In box-and-whisker plots, the boxes cover the middle two quartiles of each dataset, horizontal lines inside the boxes indicate median values, whiskers extend to 1.5 times the interquartile range, and individual points indicate data points.

### Data availability

The data that support the findings of this study are available from the corresponding authors upon reasonable request.

### Code availability

For FlyMAX software, we used C++ Visual Studio environment (Microsoft Visual Studio 2019 v142; ISO C++17 Standard) and other detailed libraries listed in FlyMAX Software subsection. For neural network frameworks, we used PyTorch^73^ (v1.7.1) and TensorFlow^74^ (v2.10.0) for classification analysis. MATLAB codes underlying the data analysis and the computation model are available upon request from the corresponding authors.

## Acknowledgments

We gratefully acknowledge research grants from the NIH Director’s Pioneer Program, the NSF NeuroNex Program, and a Samsung Scholarship for their support of our work. Special thanks to Tony H. Kim and Radoslaw Chrapkiewicz for their invaluable technical assistance.

## References

1. Loesche, F. & Reiser, M. B. An Inexpensive, High-Precision, Modular Spherical Treadmill Setup Optimized for Drosophila Experiments. Front. Behav. Neurosci. 15, 689573 (2021).

2. Manjila, S. B. & Hasan, G. Flight and Climbing Assay for Assessing Motor Functions in Drosophila. Bio Protoc 8, e2742 (2018).

3. van der Bliek, A. M. & Meyerowitz, E. M. Dynamin-like protein encoded by the Drosophila shibire gene associated with vesicular traffic. Nature 351, 411–414 (1991).

4. Aso, Y. et al. The neuronal architecture of the mushroom body provides a logic for associative learning. Elife 3, e04577 (2014).

5. Bjedov, I. et al. Mechanisms of life span extension by rapamycin in the fruit fly Drosophila melanogaster. Cell Metab. 11, 35–46 (2010).

6. Jennings, B. H. Drosophila – a versatile model in biology & medicine. Mater. Today 14, 190–195 (2011).

7. Branson, K., Robie, A. A., Bender, J., Perona, P. & Dickinson, M. H. High-throughput ethomics in large groups of Drosophila. Nat. Methods 6, 451–457 (2009).

8. Toma, D. P., White, K. P., Hirsch, J. & Greenspan, R. J. Identification of genes involved in Drosophila melanogaster geotaxis, a complex behavioral trait. Nat. Genet. 31, 349–353 (2002).

9. Rand, M. D., Montgomery, S. L., Prince, L. & Vorojeikina, D. Developmental toxicity assays using the Drosophila model. Curr. Protoc. Toxicol. 59, 1.12.1–20 (2014).

10. Flood, T. F., Gorczyca, M., White, B. H., Ito, K. & Yoshihara, M. A large-scale behavioral screen to identify neurons controlling motor programs in the Drosophila brain. G3 3, 1629–1637 (2013).

11. Munnik, C., Xaba, M. P., Malindisa, S. T., Russell, B. L. & Sooklal, S. A. Drosophila melanogaster: A platform for anticancer drug discovery and personalized therapies. Front. Genet. 13, 949241 (2022).

12. Hawker, C. D. Laboratory automation: total and subtotal. Clin. Lab. Med. 27, 749–70, vi (2007).

13. Boyd, J. Tech.Sight. Robotic laboratory automation. Science 295, 517–518 (2002).

14. Yorozu, S. et al. Distinct sensory representations of wind and near-field sound in the Drosophila brain. Nature 458, 201–205 (2009).

15. Kim, A. J., Fenk, L. M., Lyu, C. & Maimon, G. Quantitative Predictions Orchestrate Visual Signaling in Drosophila. Cell 168, 280–294.e12 (2017).

16. Kim, S. S., Hermundstad, A. M., Romani, S., Abbott, L. F. & Jayaraman, V. Generation of stable heading representations in diverse visual scenes. Nature 576, 126–131 (2019).

17. Robie, A. A. et al. Mapping the Neural Substrates of Behavior. Cell 170, 393–406.e28 (2017).

18. Takemura, S.-Y. et al. A connectome of a learning and memory center in the adult Drosophila brain. Elife 6, (2017).

19. Zheng, Z. et al. A Complete Electron Microscopy Volume of the Brain of Adult Drosophila melanogaster. Cell 174, 730–743.e22 (2018).

20. Scheffer, L. K. et al. A connectome and analysis of the adult Drosophila central brain. Elife 9, (2020).

21. Huang, C. et al. Dopamine-mediated interactions between short- and long-term memory dynamics. Nature (2024) doi:10.1038/s41586-024-07819-w.

22. Hindmarsh Sten, T., Li, R., Otopalik, A. & Ruta, V. Sexual arousal gates visual processing during Drosophila courtship. Nature 595, 549–553 (2021).

23. Bartholomew, N. R., Burdett, J. M., VandenBrooks, J. M., Quinlan, M. C. & Call, G. B. Impaired climbing and flight behaviour in Drosophila melanogaster following carbon dioxide anaesthesia. Sci. Rep. 5, 15298 (2015).

24. MacMillan, H. A., Nørgård, M., MacLean, H. J., Overgaard, J. & Williams, C. J. A. A critical test of Drosophila anaesthetics: Isoflurane and sevoflurane are benign alternatives to cold and CO2. J. Insect Physiol. 101, 97–106 (2017).

25. Mackenzie, K. & Muller, H. J. Mutation effects of ultra-violet light in Drosophila. Proc. Biol. Sci. 129, 491–517 (1940).

26. Stark, W. S., Walker, K. D. & Eidel, J. M. Ultraviolet and blue light induced damage to the Drosophila retina: microspectrophotometry and electrophysiology. Curr. Eye Res. 4, 1059–1075 (1985).

27. Savall, J., Ho, E. T. W., Huang, C., Maxey, J. R. & Schnitzer, M. J. Dexterous robotic manipulation of alert adult Drosophila for high-content experimentation. Nat. Methods 12, 657–660 (2015).

28. Alisch, T., Crall, J. D., Kao, A. B., Zucker, D. & de Bivort, B. L. MAPLE (modular automated platform for large-scale experiments), a robot for integrated organism-handling and phenotyping. Elife 7, (2018).

29. Grover, D., Katsuki, T. & Greenspan, R. J. Flyception: imaging brain activity in freely walking fruit flies. Nat. Methods 13, 569–572 (2016).

30. Bath, D. E. et al. FlyMAD: rapid thermogenetic control of neuronal activity in freely walking Drosophila. Nat. Methods 11, 756–762 (2014).

31. Williamson, W. R., Peek, M. Y., Breads, P., Coop, B. & Card, G. M. Tools for Rapid High-Resolution Behavioral Phenotyping of Automatically Isolated Drosophila. Cell Rep. 25, 1636–1649.e5 (2018).

32. Flores-Valle, A., Honnef, R. & Seelig, J. D. Automated long-term two-photon imaging in head-fixed walking Drosophila. J. Neurosci. Methods 368, 109432 (2022).

33. Maisak, M. S. et al. A directional tuning map of Drosophila elementary motion detectors. Nature 500, 212–216 (2013).

34. Strother, J. A. et al. The Emergence of Directional Selectivity in the Visual Motion Pathway of Drosophila. Neuron 94, 168–182.e10 (2017).

35. Benzer, S. BEHAVIORAL MUTANTS OF Drosophila ISOLATED BY COUNTERCURRENT DISTRIBUTION. Proc. Natl. Acad. Sci. U. S. A. 58, 1112–1119 (1967).

36. Huang, C. et al. Long-term optical brain imaging in live adult fruit flies. Nat. Commun. 9, 872 (2018).

37. Krizhevsky, A., Sutskever, I. & Hinton, G. E. ImageNet classification with deep convolutional neural networks. Commun. ACM 60, 84–90 (2017).

38. Mathis, A. et al. DeepLabCut: markerless pose estimation of user-defined body parts with deep learning. Nat. Neurosci. 21, 1281–1289 (2018).

39. Tan, M. & Le, Q. EfficientNetV2: Smaller Models and Faster Training. in Proceedings of the 38th International Conference on Machine Learning (eds. Meila, M. & Zhang, T.) vol. 139 10096–10106 (PMLR, 18--24 Jul 2021).

40. Chyb, S. & Gompel, N. Atlas of Drosophila Morphology: Wild-Type and Classical Mutants. (Academic Press, 2014).

41. Kaufman, T. C. A Short History and Description of Drosophila melanogaster Classical Genetics: Chromosome Aberrations, Forward Genetic Screens, and the Nature of Mutations. Genetics 206, 665–689 (2017).

42. Borst, A., Haag, J. & Mauss, A. S. How fly neurons compute the direction of visual motion. J. Comp. Physiol. A Neuroethol. Sens. Neural Behav. Physiol. 206, 109–124 (2020).

43. Fisher, Y. E., Silies, M. & Clandinin, T. R. Orientation Selectivity Sharpens Motion Detection in Drosophila. Neuron 88, 390–402 (2015).

44. Haag, J., Mishra, A. & Borst, A. A common directional tuning mechanism of motion-sensing neurons in the ON and in the OFF pathway. Elife 6, (2017).

45. Kitamoto, T. Conditional modification of behavior in Drosophila by targeted expression of a temperature-sensitive shibire allele in defined neurons. J. Neurobiol. 47, 81–92 (2001).

46. Bourbeau, P. P. & Ledeboer, N. A. Automation in clinical microbiology. J. Clin. Microbiol. 51, 1658–1665 (2013).

47. Armbruster, D. A., Overcash, D. R. & Reyes, J. Clinical Chemistry Laboratory Automation in the 21st Century - Amat Victoria curam (Victory loves careful preparation). Clin. Biochem. Rev. 35, 143–153 (2014).

48. Bailey, A. L., Ledeboer, N. & Burnham, C.-A. D. Clinical Microbiology Is Growing Up: The Total Laboratory Automation Revolution. Clin. Chem. 65, 634–643 (2019).

49. Ubbens, J. R. & Stavness, I. Corrigendum: Deep Plant Phenomics: A Deep Learning Platform for Complex Plant Phenotyping Tasks. Front. Plant Sci. 8, 2245 (2017).

50. Esteva, A. et al. Corrigendum: Dermatologist-level classification of skin cancer with deep neural networks. Nature 546, 686 (2017).

51. Diez-Hermano, S., Ganfornina, M. D., Vegas-Lozano, E. & Sanchez, D. Machine Learning Representation of Loss of Eye Regularity in a Drosophila Neurodegenerative Model. Front. Neurosci. 14, 516 (2020).

52. Button, K. S. et al. Power failure: why small sample size undermines the reliability of neuroscience. Nat. Rev. Neurosci. 14, 365–376 (2013).

53. Prinz, F., Schlange, T. & Asadullah, K. Believe it or not: how much can we rely on published data on potential drug targets? Nat. Rev. Drug Discov. 10, 712–712 (2011).

54. Collaboration, O. S. Estimating the reproducibility of psychological science. Science 349, aac4716 (2015).

55. Jean David, R., et al. Cold stress tolerance in Drosophila: analysis of chill coma recovery in D. Melanogaster. J. Therm. Biol. 23, 291–299 (1998).

56. Shen, J. et al. CO2 anesthesia on Drosophila survival in aging research. Arch. Insect Biochem. Physiol. 103, e21639 (2020).

57. Sinha, S. et al. High-speed laser microsurgery of alert fruit flies for fluorescence imaging of neural activity. Proc. Natl. Acad. Sci. U. S. A. 110, 18374–18379 (2013).

58. Newly installed fly-flipping robot up and running at the Stowers Institute. Stowers Institute for Medical Research https://www.stowers.org/news/newly-installed-fly-flipping-robot-and-running-stowers-institute (2011).

59. Prüßing, K., Voigt, A. & Schulz, J. B. Drosophila melanogaster as a model organism for Alzheimer’s disease. Mol. Neurodegener. 8, 35 (2013).

60. Şentürk, M. & Bellen, H. J. Genetic strategies to tackle neurological diseases in fruit flies. Curr. Opin. Neurobiol. 50, 24–32 (2018).

61. Daborn, P. J. et al. A single p450 allele associated with insecticide resistance in Drosophila. Science 297, 2253–2256 (2002).

62. Huang, J. & Lee, Y. The power of Drosophila genetics in studying insect toxicology and chemical ecology. Crop Health 1, 12 (2023).

63. Gu, S., Holly, E., Lillicrap, T. & Levine, S. Deep Reinforcement Learning for Robotic Manipulation with Asynchronous Off-Policy Updates. arXiv [cs.RO] (2016).

64. OpenAI et al. Learning Dexterous In-Hand Manipulation. arXiv [cs.LG] (2018).

65. Deng, Y., et al. Deep Reinforcement Learning for Robotic Pushing and Picking in Cluttered Environment. arXiv [cs.RO] (2023).

66. Bradski, G. The OpenCV Library. Dr. Dobb’s Journal of Software Tools (2000).

67. Guennebaud, G., Jacob, B. & Others. Eigen v3. http://eigen.tuxfamily.org (2010).

68. Zhang, Z. & Shang, H. Low-cost Solution for Vision-based Robotic Grasping. in 2021 International Conference on Networking Systems of AI (INSAI) 54–61 (IEEE, 2021).

69. Otsu, N. A threshold selection method from gray-level histograms. IEEE Trans. Syst. Man Cybern. 9, 62–66 (1979).

70. Madabattula, S. T. et al. Quantitative Analysis of Climbing Defects in a Drosophila Model of Neurodegenerative Disorders. J. Vis. Exp. e52741 (2015).

71. Kingma, D. P. & Ba, J. Adam: A Method for Stochastic Optimization. arXiv [cs.LG] (2014).

72. Seelig, J. D. et al. Two-photon calcium imaging from head-fixed Drosophila during optomotor walking behavior. Nat. Methods 7, 535–540 (2010).

73. Paszke, A. et al. PyTorch: An imperative style, high-performance deep learning library. Adv. Neural Inf. Process. Syst. 32, (2019).

74. Abadi, M., et al. TensorFlow: Large-Scale Machine Learning on Heterogeneous Distributed Systems. arXiv [cs.DC] (2016).

